# Quantitative dissection of the metastatic cascade at single colony resolution

**DOI:** 10.64898/2026.02.20.706841

**Authors:** Chris Roberts, Andy Xu, Xiangwei Fang, Adrienne Visani, Chien-Wei Peng, Xulei Qin, Irenaeus C. C. Chan, Madeline Dunterman, David A. Giles, Yaoxuan You, Isabella Guppy, Zhang Yang, Timothy Woodiwiss, Albert H. Kim, Alexander H. Stegh, Guolan Lu, Feng Chen, Li Ding, Rui Tang

**Affiliations:** Taylor Family Department of Neurosurgery, Washington University School of Medicine, St. Louis, MO, USA; The Brain Tumor Center, Alvin J. Siteman Comprehensive Cancer Center, Washington University School of Medicine, St. Louis, MO, USA; Cancer Biology Graduate Program, Washington University School of Medicine, Saint Louis, MO, USA.; Department of Medicine, Washington University in St. Louis, St. Louis, MO, USA.; Department of Surgery, Washington University School of Medicine, St. Louis, MO, USA; Department of Surgery, Stanford University, Stanford, CA, USA; Department of Urology, Stanford University, CA, USA; Department of Genetics, Washington University School of Medicine, St. Louis, MO, USA

## Abstract

Metastasis is the leading cause of cancer-related deaths. However, the core determinants and mechanistic principles underlying the metastatic cascade remain elusive. Small cell lung cancer (SCLC) is a highly aggressive malignancy with exceptional metastatic potential and limited therapeutic options. Here, we present Metastasis Originated Barcode Sequencing (MOBA-seq), a high-throughput *in vivo* platform that systematically maps genetic regulators across the metastatic cascade at single-colony resolution. MOBA-seq integrates scalable barcode-based lineage tracing with a computational pipeline that quantitatively deconvolutes genotype-specific effects on metastatic seeding, dormancy, and clonal expansion across hundreds of thousands of metastatic events. Applying this approach to more than 400 candidate regulators of SCLC, we uncovered tissue-specific metastatic suppressors and universal metastatic essential genes. We identified metastatic seeding as the predominant determinant of metastasis. Comparative analysis across recipient mice of distinct genetic backgrounds further revealed that innate immune surveillance constrains metastatic progression by reducing metastatic seeding and enforcing dormancy, with additional modulation by sex and tissue context. We validated the frequently mutated gene *CREBBP* as a key metastasis suppressor whose loss enhances SCLC metastasis through both tumor-intrinsic and immune-modulatory mechanisms. This work establishes a scalable and quantitative platform for mapping the metastatic fitness landscape at single-colony resolution across hundreds of thousands of *in vivo* data points. Our approach offers a broadly applicable framework for dissecting the interactions between cancer-intrinsic and microenvironmental factors governing tumor initiation, progression, and therapeutic response.

## Introduction

Metastasis is the terminal and most lethal phase of cancer progression, accounting for the vast majority of cancer-related deaths. In 2025 alone, an estimated 290,000 new cases of metastatic cancer will be diagnosed, leading to over 240,000 deaths^1^. Despite its prevalence and lethality, the cellular and molecular mechanisms for cancer metastasis are still poorly understood. Metastasis is a complicated multi-step process encompassing primary cancer cell dissemination, survival through the circulatory and lymphatic systems, metastatic seeding in a distant site, dormancy escape, and clonal expansion to form tissue destructive metastatic lesions^2^. At each stage of the metastatic cascade, cancer-intrinsic factors, including various transcriptional and epigenetic regulators, co-operate with dynamic extrinsic factors. This includes immune surveillance, tissue-specific microenvironments, and patient-specific variables including sex and age, to shape metastatic fitness^3,4^. A systematic framework that integrates these genetic and environmental influences across the metastatic cascade will offer comprehensive insights into the biology of metastasis and identify actionable targets for early intervention and better treatment of this lethal disease.

Small Cell Lung Cancer (SCLC) is a recalcitrant and highly metastatic cancer with approximately 300,000 new cases annually worldwide^5^. Approximately 70% of SCLC patients are diagnosed with extensive stage disease (ED) with distant metastases, most frequently in the liver and brain, resulting in a five-year survival rate below 3%^6–8^. SCLC freely travels through the circulatory system, resulting in an unusually high circulating tumor cell (CTC) burden in patients^9^. The highly metastatic nature, clear organotropic pattern, and high CTC content of SCLC present an opportunity to study the metastatic cascade with precision and detail. Despite the identification of a few frequently mutated genes and their mechanistic insights, no targeted therapies are yet available for SCLC patients^10,11^. Intriguingly, some ED-SCLC patients respond well to a combination of chemo-/radiation– and immunotherapies, suggesting metastatic SCLC may have a unique cancer immune microenvironment^12^. Studying SCLC metastasis in preclinical models to decode cancer intrinsic and extrinsic environmental factors of the metastatic cascade could nominate new avenues for targeted therapies in ED-SCLC patients.

Current approaches for studying metastasis, such as histological staining and imaging modalities including bioluminescence and other whole-body techniques, have provided valuable insights into the metastatic cascade^13,14^. However, these methods are often limited in their ability to provide sufficient quantitative information about the metastatic cascade. Histological analysis of metastatic colony size and amount can lead to sampling concerns due to the limited regions of tissue profiled on slides. Whole body imaging technologies do not allow for analysis of different steps across the metastatic cascade and are usually limited by their resolution. Most importantly, *in vivo* genetic studies utilizing these methods are often limited by low multiplexing capacity and insufficient statistical power when profiling large gene sets^15,16^. Genetically heritable DNA barcoding has emerged as a powerful strategy for lineage tracing and for revealing cellular state transitions during cancer evolution^17–19^. Yet, most studies employing barcoding have focused on reconstructing the trajectories of a few dominant tumor clones, overlooking the potential of colony-level barcoding to quantitatively resolve tumor fitness across diverse metastatic populations. Given the high mutational burden and heterogeneity of SCLC, there is a pressing need for scalable, high-throughput, and quantitative technologies capable of delineating cell-intrinsic determinants at each stage of the metastatic cascade ^20^.

In this study, we developed the <u>M</u>etastasis-<u>O</u>riginated <u>Ba</u>rcode <u>Seq</u>uencing (MOBA-seq) technology to generate hundreds of thousands of high-resolution *in vivo* data to study genetic drivers of SCLC metastasis. MOBA-seq utilizes a combination of tumor colony barcoding and CRISPR-Cas9 screening to deconvolute genotype-specific effects on metastatic seeding, dormancy, and clonal expansion in a high-throughput quantitative manner at single colony resolution. In conjunction with MOBA-seq, we also developed a computational pipeline designed to analyze sequencing outputs from MOBA-seq (https://github.com/tanglab-2024). By measuring colony size of hundreds of thousands of uniquely barcoded metastases per genotype at near 10 cells resolution, we identified and validated novel metastatic drivers and uncovered new mechanisms underlying SCLC metastasis, organotropism, and immune response. We functionally validated a frequently mutated SCLC gene coding for CREB-binding protein (CREBBP), as a potent metastatic suppressor. In the SCLC metastatic cascade, CREBBP functions to suppress dissemination, seeding, and clonal outgrowth. However, CREBBP deficiency also facilitates dormancy and heightens vulnerability to immune surveillance. We envision that MOBA-seq can be broadly applied for quantitative measurements of tumor fitness effects in other implantation-based preclinical mouse models for other primary and metastatic cancer types.

## Results

### Establish the MOBA-seq pipeline

To quantify genotype-specific effects on SCLC metastasis at single-colony resolution, we developed Metastasis-Originated Barcode Sequencing (MOBA-seq, **Figure 1a**), an integrated platform combining pooled CRISPR screening with colony barcode sequencing in implantation-based mouse models^21–23^. We adopted a published design where a 17-nucleotide random barcode was inserted into the –22 to –5 – nucleotide region of a bovine U6 (bU6) promoter, preserving transcriptional activity while enabling highly diverse barcode-sgRNA pairings^16,24^. We cloned sgRNA oligo pool into this barcoded backbone and established more than 2×10^6^ bacteria colonies that contain unique bU6-barcode-sgRNA constructs (**Figure S1a**). We transduced Cas9-expressing SCLC cell lines with the lenti-bU6-barcode-sgRNA library at a low multiplicity of infection (MOI = 0.1), followed by puromycin selection (**Figure S1b**). This generated a log-normal distributed SCLC cell library with over 1×10^5^ unique barcode-sgRNA pairs at an average of 40 cells per barcode-sgRNA pair (**Figure S1c**).

**Figure 1.**
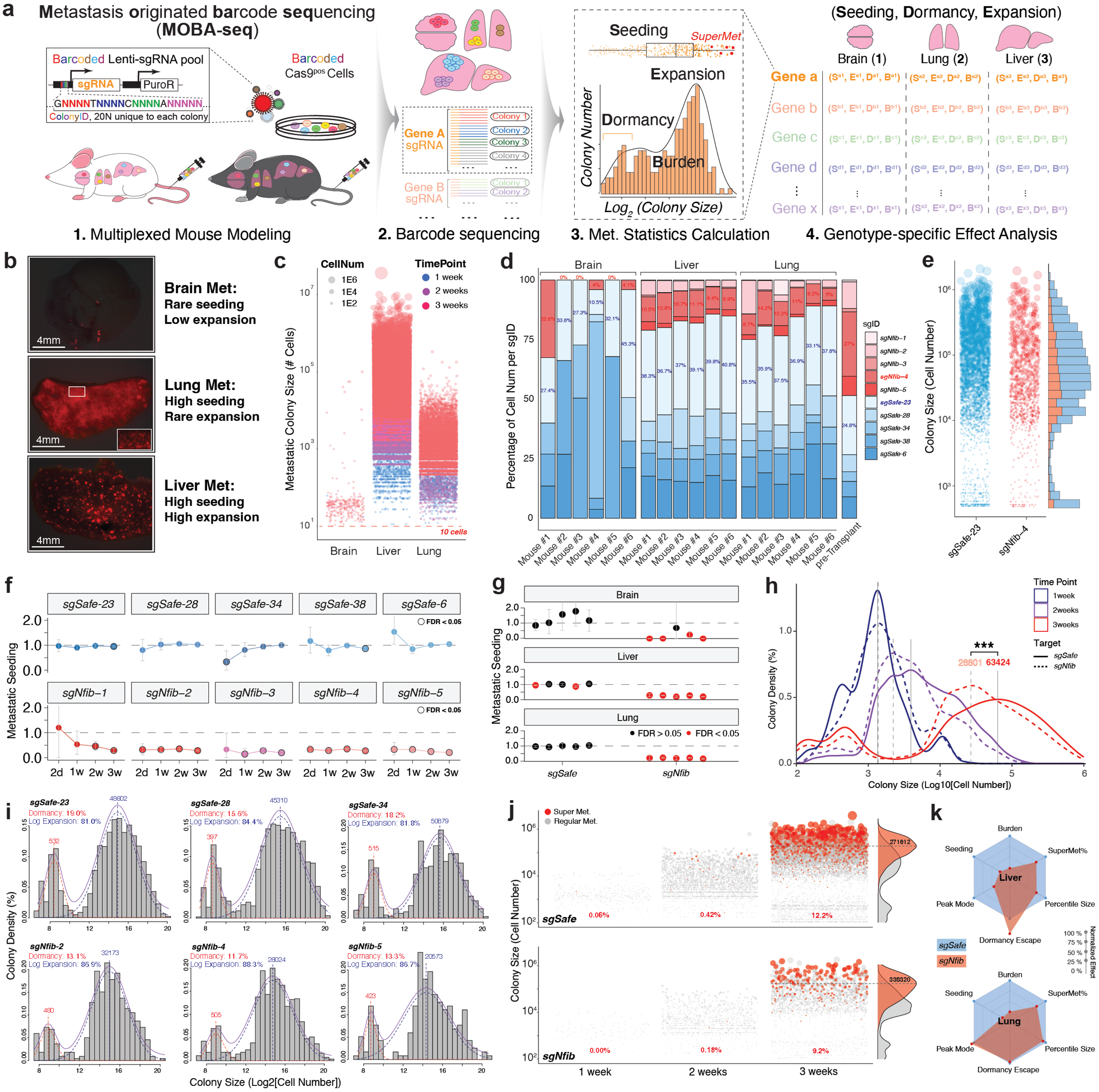
MOBA-seq quantitatively deconvolutes genotype-specific metastatic effects at single colony resolution. **a.** Schematic representation of the MOBA-seq pipeline. Cas9^+^ cancer cells are transduced with a randomly barcoded sgRNA lentiviral library before transplantation into recipient mice. After tumor initiation, target tissues are collected for barcode sequencing and analyzed to determine genotype-specific effects on metastatic seeding, dormancy, clonal expansion, and overall metastatic burden. **b.** Tissue-specific SCLC metastasis in tail vein transplantation mouse model. Fluorescent images showing surface metastases from mCherry^+^ RP48 cells transplanted into the tail-vein at three weeks. Scale bar indicated 4 mm, white box shows enlarged region of lung micro-metastases. **c.** MOBA-seq digitally reconstitutes metastatic pattern across different tissues in mice. Barcoded RP48 cells are transplanted into the tail-vein of NSG mice. Brain, lung, and liver tissues were harvested at one, two, and three weeks for MOBA-seq analysis. Each individual dot represents a uniquely barcoded metastatic colony, with dot size correlating to the size of the colony, and each color represents a different time point. **d.** MOBA-seq consistently measures *Nfib*-specific metastatic effects across tissues and mice. RP48-Cas9 cells are transduced with a lenti-sgRNA library containing five sgRNAs targeting *Nfib* and five control sgRNAs before transplanting into six NSG mice for three weeks. The total metastatic cell representation of each sgRNA is shown across tissues for each mouse, as well as a pre-transplantation representation. The percentage of *sgNfib-4* and *sgSafe-23* expressing RP48 cells across each sample is marked. **e.** Inactivation of *Nfib* suppresses RP48 metastatic seeding. Metastatic colonies from six mice livers for *sgSafe-23* and *sgNfib-4* are plotted. The plot on the left shows individual tumor sizes. The bar plot on the right shows colony size density for both genotypes. **f.** MOBA-seq reliably and consistently measures *Nfib*-specific metastatic seeding over time and across five sgRNAs. Relative metastatic seeding of *sgSafe*– and *sgNfib*-expressing RP48 cells was measured at two days, and one, two, and three weeks. Each dot is shown as fold change ± CI. FDR<0.05 is signified with a bold outline. **g.** MOBA-seq consistently measures metastatic seeding in brain, liver and lung. Relative metastatic seeding of *sgSafe*– and *sgNfib*-expressing cells was compared between tissues. Each dot is a sgRNA and is shown as fold change ± CI. FDR <0.05 are marked in red. **h.** MOBA-seq reveals changes in metastatic colony size distributions in the liver over time. Density plot showing colony size distributions for *sgNfib* and *sgSafes* over three weeks with a spectrum from 1×10^2^ to 1×10^6^ cells. Color represents time points and line type represents genotype. PeakMode of major colony population is designated with a vertical line, and a significant difference between PeakMode of *sgSafe* and *sgNfib* at 3 weeks is detected and marked. *** FDR < 0.0001 **i.** MOBA-seq quantifies genotype-specific dormancy in metastasis. A Gaussian mixture model is used to demultiplex the dormancy and lognormal expansion populations in SLCL liver metastasis. The percentages of dormancy and log expansion metastatic colonies are marked. Note that the inactivation of *Nfib* suppressed dormancy while also suppressing log expansion peak mode. **j.** MOBA-seq identified colonies that can disseminate circulating SCLC cells into the blood. Blood cells from liver-metastases bearing mice were sequenced via MOBA-seq. Matching colony barcodes between blood and liver SCLC were used to identify liver metastatic colonies that disseminate circulating SCLC, termed SuperMet, and highlighted in red. SuperMets over three weeks from control and *Nfib*-inactivated colonies are shown. Plotted to the right is the size distribution of SuperMet colonies after three weeks. Note that inactivation of *Nfib* increased the PeakMode and reduced the frequency of liver SuperMet colonies. **k.** Radar plots showing *Nfib*-specific metastatic effects in the liver (*Top*) and lung (*Bottom*) on overall metastatic burden, metastatic seeding, PeakMode of log expansion, Dormancy escape, 90th percentile colony size, and SuperMet percentage.

To establish the MOBA-seq pipeline, we used a SCLC metastasis model by tail vein injection of mCherry-expressing murine (RP48) and human (H82) cells into *NOD/Scid/γC* (*NSG*) mice^6,25,26^. Via flow cytometry analysis, we found that transplanted H82 cells were initially detected in the lung and gradually redistributed to the liver over two days post-transplantation (**Figure S1d**). Three weeks post-injection, fluorescent images of tissue surface RP48 metastases showed distinct, organ-specific metastatic profiles: the brain exhibited rare metastatic seeding with low clonal expansion, the lung showed high metastatic seeding but minimal clonal expansion, and the liver displayed both high metastatic seeding and robust clonal expansion (**Figure 1b**). In parallel, we also transplanted barcoded syngeneic murine RP48 cells into NSG mice and applied MOBA-seq for metastases analysis in the liver, lung, and brain after a period of three weeks^27^. To quantify barcoded metastatic colony size, a predefined number of spike-in cells was added to each tissue prior to genomic DNA extraction and sequencing. We then built a linear regression using the spike-in barcode read counts and their expected cell numbers from each sample to enable quantitative recovery of metastatic colonies at high accuracy (**Figure S1e-g**). We developed <u>L</u>ineage <u>E</u>valuation and <u>T</u>racing <u>T</u>hrough <u>U</u>nique <u>C</u>ellular <u>E</u>vents (LETTUCE), a dedicated MOBA-seq data analysis package (**Figure S1h**), which extracts barcode-sgRNA reads, applies spike-in–based normalization, and reconstructs metastatic colony sizes. LETTUCE also supports statistical modeling of genotype-specific effects on metastatic seeding, dormancy escape, clonal expansion, circulating cancer cell (CTC) dissemination, and metastatic burden. Tail vein transplantation of barcoded RP48 cells seeded throughout the mouse tissues and expanded over time, resulting in 5,000-20,000 genetically diverse barcoded metastatic colonies per mouse with the vast majority detected in only a single mouse (**Figure S1i-j**). MOBA-seq reliably calculated genotype-specific metastatic seeding profiles across mice (**Figure S1k-l**). MOBA-seq also detects CTC in both C57BL/6 and NSG mice with high-sensitivity (**Figure S1m**).

With hundreds of thousands of barcoded metastatic colonies, MOBA-seq digitally reconstituted SCLC metastatic profiles with single colony resolution that recapitulated the organ-specific metastatic patterns observed via fluorescent imaging. Importantly, by leveraging high depth sequencing, MOBA-seq supports detection of metastatic colonies as small as 10 cells, allowing for identification of rare, early-stage metastases across organs (**Figure 1c**). This ultra-high sensitivity allows for comprehensive characterization of the metastatic cascade, from seeding to clonal outgrowth throughout the metastatic cascade.

### MOBA-seq quantifies genotype-specific impact on SCLC metastatic profiles

To benchmark the robustness of MOBA-seq in quantifying genetic effects on metastasis, we targeted the well-established SCLC metastatic driver *Nfib*^11^. *NSG* mice were transplanted with an RP48-Cas9 MOBA-seq library containing five sgRNAs targeting *Nfib* and five control sgRNAs targeting *safe* harbor loci of mouse genome (**Figure S2a**)^28^. Metastatic burden was quantified based on barcode-derived cell counts across the brain, liver, and lung over a period of three weeks post-injection, alongside a representative pre-transplantation library (**Figure 1d, Figure S2b-c**). In both liver and lung, sgRNA representations are reliably consistent across biological replicates, while brain samples have a higher inter-mouse diversity, likely due to the stochastic nature of rare metastatic seeding events. In all tissues and mice, inactivation of *Nfib* suppressed overall metastatic burden, in agreement with prior studies^11,29^. To directly visualize the effect of *Nfib* inactivation on metastatic seeding, we compared liver metastases carrying *sgNfib#4* versus *sgSafe#23* –two sgRNAs with matched representation in the pre-transplantation pool. MOBA-seq revealed a reduction in seeding by *sgNfib*#4, reflected by fewer uniquely barcoded colonies in the liver and other tissues (**Figure 1e**). To quantify sgRNA-specific effects, we computed the fold change of metastatic seeding for each sgRNA targeting *Nfib* in the liver relative to the mean seeding of all control sgSafe guides over time. All five sgRNAs targeting *Nfib* show consistent effects on the suppression of seeding over 2 days, 1 week, 2 weeks, and 3 weeks timepoints (**Figure 1f, Figure S2d**). At the 3-week endpoint, *sgNfib*-specific metastatic seeding patterns remained highly reproducible in the liver and lung, with greater variability observed in the brain due to its rare and stochastic seeding pattern (**Figure 1g**). Interestingly, time-course analysis revealed organ-specific dynamics of *Nfib*-dependent seeding. The liver maintained a stable ratio between *sgNfib* and *sgSafe* metastatic colonies over time, whereas this ratio steadily declined in the lung, suggesting selective elimination of *Nfib*-deficient colonies in the lung, likely mediated by local innate immune surveillance^30^.

To assess the effect of *Nfib*-deficiency on metastatic outgrowth, we calculated SCLC colony sizes at different percentiles across genotypes^21,23^. While *Nfib*-inactivated metastases were consistently smaller than corresponding controls, statistical power varied across percentile-based comparisons (**Figure S2e**). To more comprehensively quantify genotype-specific clonal expansion, we plotted colony size distribution on log-scale and defined PeakMode as the most frequently observed colony size within each genotype. PeakMode emphasizes exponentially expanding macro-metastases. Inactivation of *Nfib* significantly reduced PeakMode colony size in the liver but not in the lung, indicating tissue-specific suppression of metastatic outgrowth (**Figure 1h, Figure S2f**). To quantify dormancy by analyzing micro-metastases, we modeled the log-scaled colony size distribution across liver, lung, and brain under the assumption that dormant colonies fall below a defined size threshold (“valley model”) (**Figure S2g**). A Gaussian Mixture Model (GMM) was applied to unmix dormant from proliferative colonies within each genotype, allowing us to quantify the proportion of clones remaining in a dormant state. Surprisingly, *Nfib* inactivation reduced proliferative expansion while also increasing dormancy escape in the liver (**Figure 1i, Figure S2h**).

Finally, leveraging barcode-derived lineage tracing across tissues within the same mouse, we were able to identify a subset of metastatic clones capable of re-dissemination via the circulation, termed “SuperMets”. In wild-type liver metastases, SuperMets predominantly originated from large, clonally expanded lesions and only emerged at the three-week time point (**Figure 1j upper)**. This suggests SuperMet represents a late stage of SCLC clonal expansion. *Nfib* inactivation reduced the frequency of SuperMet colonies but increased the PeakMode size of the remaining SuperMet colonies as compared to controls, indicating a suppression of CTC dissemination (**Figure 1j lower, Figure S2i**). These results suggest that *Nfib*-deficiency restrained the late-stage transition of metastatic clones into a CTC-releasing state^31^. Importantly, MOBA-seq offers an indirect yet quantitative measure of genotype-specific CTC output in mouse models with single CTC sensitivity. To summarize, by profiling 16,738 individual metastatic colonies across three tissues, we achieved a single-colony resolution landscape of *Nfib*-dependent effects on metastatic seeding, expansion, dormancy, and re-dissemination (**Figure 1k)**. We identified markedly distinct tissue-specific effects, suggesting the role of organ microenvironments in shaping SCLC metastasis.

### MOBA-seq identifies known and novel genetic effectors in SCLC metastasis

To systematically identify genetic drivers of SCLC metastasis, we constructed a CRISPR-MOBA-seq library (“MOBA500”). This library targeted ∼250 genes recurrently mutated in SCLC patients, ∼200 SCLC cell line-specific candidate genes from prior CRISPR viability screens, and 50 control sgRNAs that were either non-targeting or targeting safe harbor loci in the mouse genome (**Table S1, Figure S3a**)^20,28,32^. Barcoded RP48 cells were transplanted into the tail vein of nine C57BL/6 mice and analyzed after three weeks (**Figure 2a**). From the harvested tissues, we recovered over 75,000 unique metastatic colonies. Comparison of sgRNA representation before and after metastasis identified tissue-specific shifts in genotype composition across liver, lung, and brain (**Figure 2b**). The overall metastasis of each genotype can be further de-multiplexed by the colony barcode to quantify metastatic colony size per genotype (**Figure 2c, Figure S3b**). As expected, sgRNAs targeting canonical tumor suppressors, such as *Tsc1, Tsc2,* and *Pten* generate the largest average metastatic colonies in the liver, confirming the sensitivity of MOBA500 in detecting pro-metastatic effects of genetic perturbations^33^.

**Figure 2.**
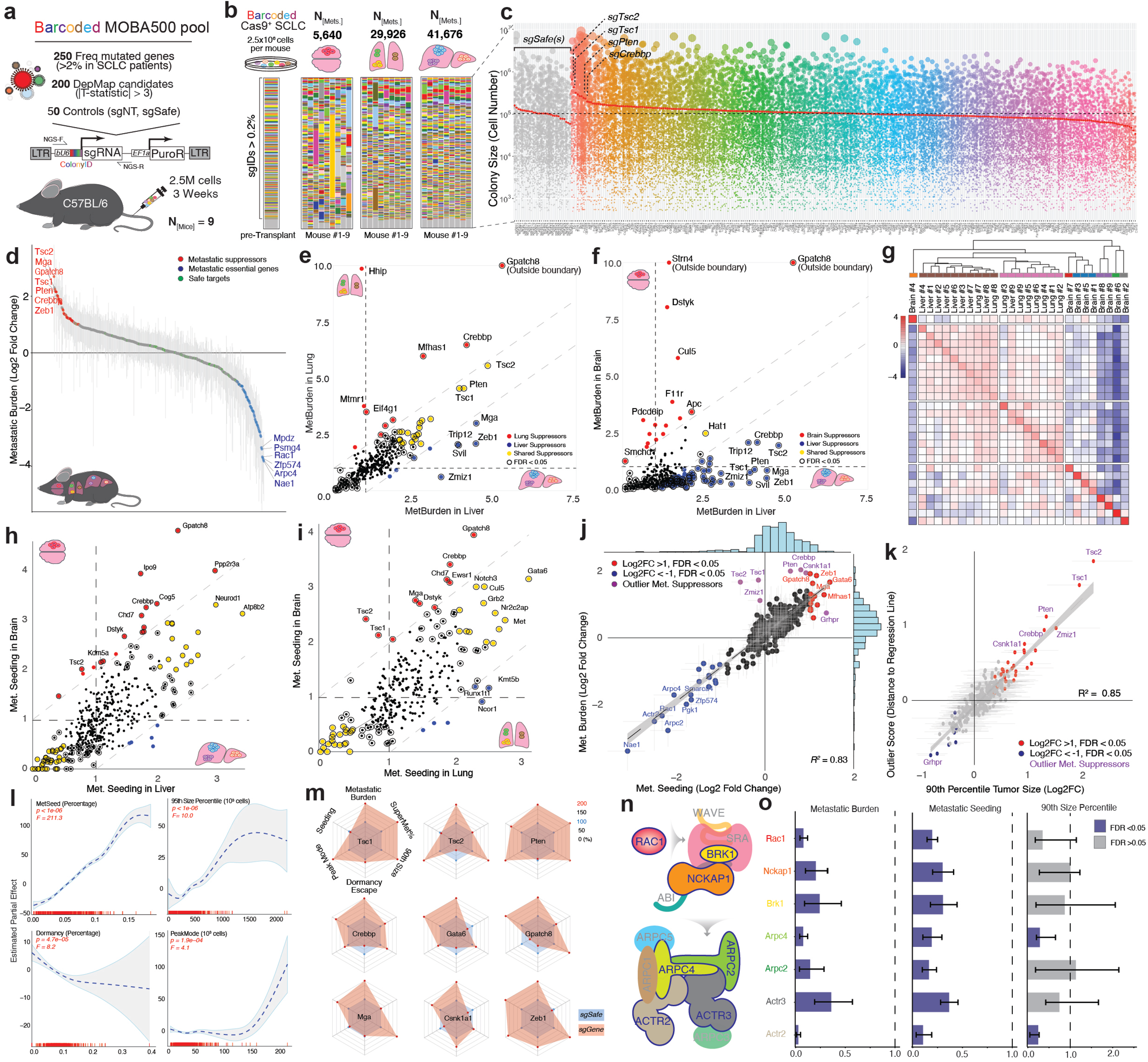
MOBA-seq identified genotype-specific metastatic effects in SCLC. **a.** Schematic of the MOBA500 screen targeting ∼450 candidate genes. A lenti-sgRNA-PuroR library contains 250 sgRNAs targeting homologs of frequently mutated genes in SCLC patients, 200 sgRNAs targeting homologs of significantly enriched genes from human cell line CRISPR screen, and 50 control sgRNAs targeting safe regions on the mouse genome. 2.5 x 10^6^ lenti-transduced RP48-Cas9 cells are tail-vein transplanted into nine C57BL/6 mice and tumors are allowed to expand for three weeks. **b.** Representation of all SCLC genotypes from three target organs across nine mice. Total metastatic colonies recovered from each target organ are marked. Cell library samples are collected before transplantation as a quantification baseline. Genotypes representing more than 0.2% of the library were labeled with color. Genotype color-mapping is the same across all samples. Note that genotype enrichment profiles among liver metastases are consistently measured across mice. **c.** MOBA-seq identifies well-known tumor suppressors in SCLC liver metastasis. Liver metastatic colonies from each genotype are ranked by their mean colony size, which is indicated by a red dot. The dashed line represents the mean colony size of all control colonies combined (*sgSafes*). Note that sgRNAs targeting known tumor suppressors *Tsc1*, *Tsc2*, and *Pten* were re-discovered to increase mean colony size in this screen. **d.** MOBA-seq identifies known and novel genetic regulators in SCLC liver metastasis. Ranked plot of each genotype highlights top candidate suppressors (red, Log2FC>1, FDR<0.05) and drivers (blue, Log2FC<-1, FDR<0.05) of SCLC liver metastatic burden. Grey error bars represent 95% confidence interval. sgRNAs that generate wild-type metastases are shown in green. **e.** MOBA-seq identifies liver– and lung-specific metastatic suppressors. Scatter plot represents fold change in metastatic burden of sgIDs in the liver and lung. Lung-specific metastatic suppressors are shown in red, liver-specific metastatic suppressors are shown in blue, and shared suppressors are shown in yellow. Dashed lines indicate a fold change difference one of one between two organs. Genotypes with FDR<0.05 are outlined in black. **f.** MOBA-seq identifies liver– and brain-specific metastatic suppressors. Similar to Figure 2e. Note that MOBA-seq identifies unique suppressors from brain metastases. **g.** Tissue-specific metastatic seeding showing clustering of liver and lung samples. Brain samples do not cluster well due to lower sample size. Genotype-specific metastatic seeding profiles across samples were clustered by tissue of origin. Color indicates the correlation coefficient among samples, a higher correlation is shown in red. **h.** Cross-comparison of genotype-specific metastatic seeding in brain and liver. Brain-specific metastatic seeding suppressors are shown in red and dashed lines indicate a fold change difference of one between two organs. Genotypes with FDR<0.05 are outlined in black. **i.** Cross-comparison of genotype-specific metastatic seeding in brain and lung. Similar to Figure 2e. Note that MOBA-seq identifies shared metastatic seeding suppressors in the brain compared to both liver and lung. **j.** Metastatic seeding is the primary factor driving overall metastatic burden. Plot of genotype-specific metastatic seeding and metastatic burden for all genotypes revealed a high linear correlation (R^2^=0.83). Note that metastatic drivers mainly function through metastatic seeding, while metastatic suppressors have more diverse effects. Outlier suppressors from linear regression are marked in purple. Grey error bars represent 95% confidence interval. **k.** The 90th percentile metastatic colony size determines outlier suppressors. The orthogonal distance of each genotype from Figure 2j is plotted against 90^th^ percentile metastatic colony size with a high linear correlation (R^2^=0.85). Red dots indicate a positive outlier distance, while blue dots indicate a negative outlier distance. Grey error bars represent a 95% confidence interval. **l.** Metastatic burden is co-determined by multiple metastatic metrics in a generalized additive model (GAM). Genotype-specific metastatic burden was modeled using metastatic seeding, 90th-percentile colony size, dormancy escape and PeakMode. Metastatic seeding accounted for the largest variation in burden, whereas 90th-percentile colony size and PeakMode contributed preferentially at different ranges of metastatic burden. Dormancy escape was inversely associated with metastatic burden. **m.** MOBA-seq identified diverse patterns for metastasis suppression. Radar plots stratify genetic suppressors by their effects on the different steps throughout the metastatic cascade. **n.** Rac1-WRC-ARP2/3 pathway cartoon for actin assembly. Genes on this pathway that the MOBA500 library targets are highlighted with a border and bolded (7/13 genes). **o.** Targeting the Rac1-WRC-ARP2/3 pathway consistently affects metastatic burden and seeding but has little effect on colony size. Each bar represents fold change of aggregated data from all nine mice. The error bars represent 95% confidence interval.

To quantify genotype-specific SCLC metastatic effects, we first calculated the relative metastatic burden for each genotype by normalizing to the mean burden of wild-type (*sgSafe*) metastases. To identify genes that significantly modulate metastatic burden, we ranked the total metastatic burden of each sgRNA in the liver, the most frequent site of metastasis in SCLC. This analysis revealed *Crebbp* as the strongest liver metastasis suppressor, alongside known tumor suppressors *Tsc1*, *Tsc2*, and *Pten*. Conversely, we identified essential metastatic driver genes such as *Nae1* and *Rac1* whose loss severely impaired SCLC liver metastasis (**Figure 2d, Figure S3c**). We ranked the total metastatic burden associated with each sgRNA across all three tissues and compared these rankings with either the homologous gene mutation frequencies in SCLC patients or cell viability data from publicly available CRISPR and RNAi screens in human SCLC cell lines (**Figure S3d-f**). This analysis identified conserved tumor suppressors, such as *PTEN* and *CREBBP*, as well as conserved essential genes, including *ASCL1*^34^. To explore tissue-specific metastatic effects, we compared the normalized metastatic burdens between the liver and lung. Despite wild-type metastases exhibiting tissue-specific differences, most sgRNAs displayed consistent genotype-specific effects across both tissues (**Figure 2e**). This suggested an unbiased suppression in the lung metastatic tumor microenvironment (TME). In contrast, comparison of liver and brain metastatic burdens revealed distinct brain-specific suppressors, indicating a unique set of genetic requirements for SCLC adaptation within the brain TME (**Figure 2f**)^35^. The presence of a unique brain metastatic pattern was further supported by clustering analyses of metastatic seeding across all samples. Liver and lung samples clustered together based on their overall metastatic seeding profiles, while brain samples formed a distinct group with lower similarity to other tissues (**Figure 2g**). To systematically identify genes regulating tissue-specific seeding, we compared genotype-specific metastatic seeding between the brain and other tissues. This analysis revealed a set of brain-specific metastatic suppressors whose inactivation promote SCLC seeding to the brain, including *Gpatch8*, *Cdh7*, *Crebbp*, and *Tsc2* (**Figure 2h-i**). These results highlight a mechanistic divergence between brain and peripheral metastases, suggesting that metastatic seeding is a critical bottleneck in the SCLC metastatic cascade.

### Metastatic phenotypes are mainly driven by metastatic seeding

Our results suggested that genetic suppressors in metastatic burden also suppressed metastatic seeding in the same tissue (**Figure 2e-f and 2h-i**). This observation led us to hypothesize that metastatic seeding is a primary determinant of overall metastatic burden. To test this, we first established that metastatic seeding is a robust and reproducible metric of metastasis that is independent of both pre-transplantation cell proliferation *in vitro* and subsequent clonal expansion *in vivo* (**Figure S4a-d**). We then compared relative metastatic seeding and metastatic burden across all genotypes in the liver from *NSG* mice and observed a strong positive correlation, suggesting that seeding frequency largely accounts for metastatic outcome. (**Figure 2j, Figure S5a**). In contrast, the correlation between the metastatic burden and all other statistics were substantially weaker (**Figure S5b-c**). We also observed that metastatic essential genes were closely aligned with the seeding-burden regression, whereas suppressors exhibited greater variability, suggesting the presence of a hidden layer of secondary determinants influencing metastatic outcome. Notably, although colony size did not directly correlate with metastatic burden, we identified a strong correlation between the 90^th^ percentile colony size and the orthogonal residual (Outlier Score) from the seeding-burden regression (**Figure 2k**). In addition, we compared the size percentiles and liver metastatic burden or seeding efficiency across different percentile groups and observed a progressive increase in fitness (R²) with distance, which plateaued beyond the 90th percentile (**Figure S5d**). We further modeled metastatic burden using a Generalized Additive Model (GAM) to dissect the relative contributions of distinct metastatic statistics. As expected, metastatic seeding emerged as the dominant determinant of genotype-specific metastatic burden, while the 90th percentile tumor size and PeakMode contributed preferentially to small and large metastatic lesions, respectively (**Figure 2l**). Collectively, these data demonstrate that metastatic seeding is the primary driver of SCLC metastatic outcome, while the outgrowth of exceptionally large colonies serves as a secondary contributor, particularly upon inactivation of select metastatic suppressors.

When comparing metastatic burden and other metastatic statistics, we found a significant difference in the burden-dormancy escape relationship between the liver and the lung, suggesting tissue-specific regulation of dormancy (**Figure S5b-c**). We therefore further examined the relationship between dormancy escape and other metastatic statistics in both tissues. In the lung, dormancy escape was strongly positively correlated with overall metastatic burden and 90th percentile colony size, but showed little correlation with metastatic seeding. In contrast, in the liver, dormancy escape exhibited a negative correlation with both metastatic burden and 90th percentile colony size, particularly among strong tumor suppressors (**Figure S5e**). This is also consistent with our previous observations of *Nfib*-dependent dormancy escape (**Figure 1i, Figure S2h**). Together, these findings indicate the existence of tissue– and genotype-specific mechanisms governing dormancy escape.

### Diversified mechanisms of metastatic suppression but a unified requirement for metastasis

Our results revealed diversified effects of metastatic suppressors on overall metastatic burden (**Figure 2j, Figure S6a-f**). We thus constructed a landscape of SCLC liver metastasis by integrating six metastatic metrics, including metastatic burden, seeding, PeakMode colony size, dormancy escape, 90th percentile colony size, and SuperMet percentage for nine metastatic suppressors (**Figure 2m**). Notably, top-ranked suppressors exhibited distinct mechanistic signatures. Canonical tumor suppressors such as *Tsc1*, *Tsc2*, and *Pten* primarily impacted PeakMode and 90th percentile colony size, with minor influence on metastatic seeding. Loss of *Crebbp*, *Gata6*, and *Gpatch8* strongly promoted metastatic seeding but also reduced dormancy escape as side effect. Most metastatic suppressors regulate multiple steps of the metastatic cascade, underscoring the mechanistic heterogeneity during metastatic regulation. Beyond suppressors, we also examined seven putative regulators of metastasis in the Rac–WAVE regulatory complex–ARP2/3 (Rac-WRC–ARP2/3) actin assembly pathway (**Figure 2n**). We were able to identify significant and ubiquitous levels of reduction in metastatic burden and metastatic seeding in recipient mice for all seven out of 13 targets in the pathway (**Figure 2o, left and middle**). Interestingly, once colonies were established and expanded, most exhibited no significant differences in 90th percentile colony size compared to controls (**Figure 2o, right**). These findings suggest that actin assembly components are essential for early metastatic seeding, but dispensable for subsequent clonal expansion. Together, these results highlight the diversity of genetic control mechanisms governing SCLC metastasis and pinpoint pathway-specific vulnerabilities across distinct steps of the metastatic cascade.

### Immune surveillance in SCLC liver metastasis mainly suppressed metastatic seeding via innate immunity

Immune surveillance is known to be one of the major determining factors in cancer metastasis^36^. To understand how the immune system shapes distinct steps of the metastatic cascade, we performed MOBA-seq by transplanting barcoded RP48 cells into both immunocompetent C57BL/6 and immunodeficient NSG mice. Compared to C57BL/6 recipients, NSG mice exhibited a markedly elevated liver metastatic burden, as evidenced by gross liver surface metastases and liver-to-body weight ratios (**Figure 3a-b**). MOBA-seq enables direct quantification of individual metastatic colonies and revealed that NSG livers harbored approximately fivefold more colonies than those of C57BL/6 mice, indicating that immune surveillance robustly eliminates metastatic seeding. (**Figure 3c**). Recent studies implicate natural killer (NK) cells as key mediators of immune control in SCLC liver metastases^26,37^. To dissect the relative contributions of adaptive and innate immune cell populations to SCLC metastatic seeding, we transplanted RP48 cells into C57BL/6, Rag2-KO, and NSG mice. Metastatic seeding events quantified per mouse by MOBA-seq revealed a substantial increase in NSG mice and only a subtle increase in Rag2-KO mice relative to C57BL/6 controls (**Figure 3d**). We further compared relative liver metastatic seeding and burden across genotypes. NSG mice clustered tightly together with low inter-mouse variation, whereas C57BL/6 and Rag2-KO mice exhibited significantly higher variability with a larger coefficient of variation (CV). (**Figure 3e**). Notably, Rag2-KO mice phenocopied C57BL/6 mice across multiple metrics. These results suggested that innate immunity, such as NK cells, acts as the primary barrier to successful SCLC metastatic seeding ^26,38^.

**Figure 3.**
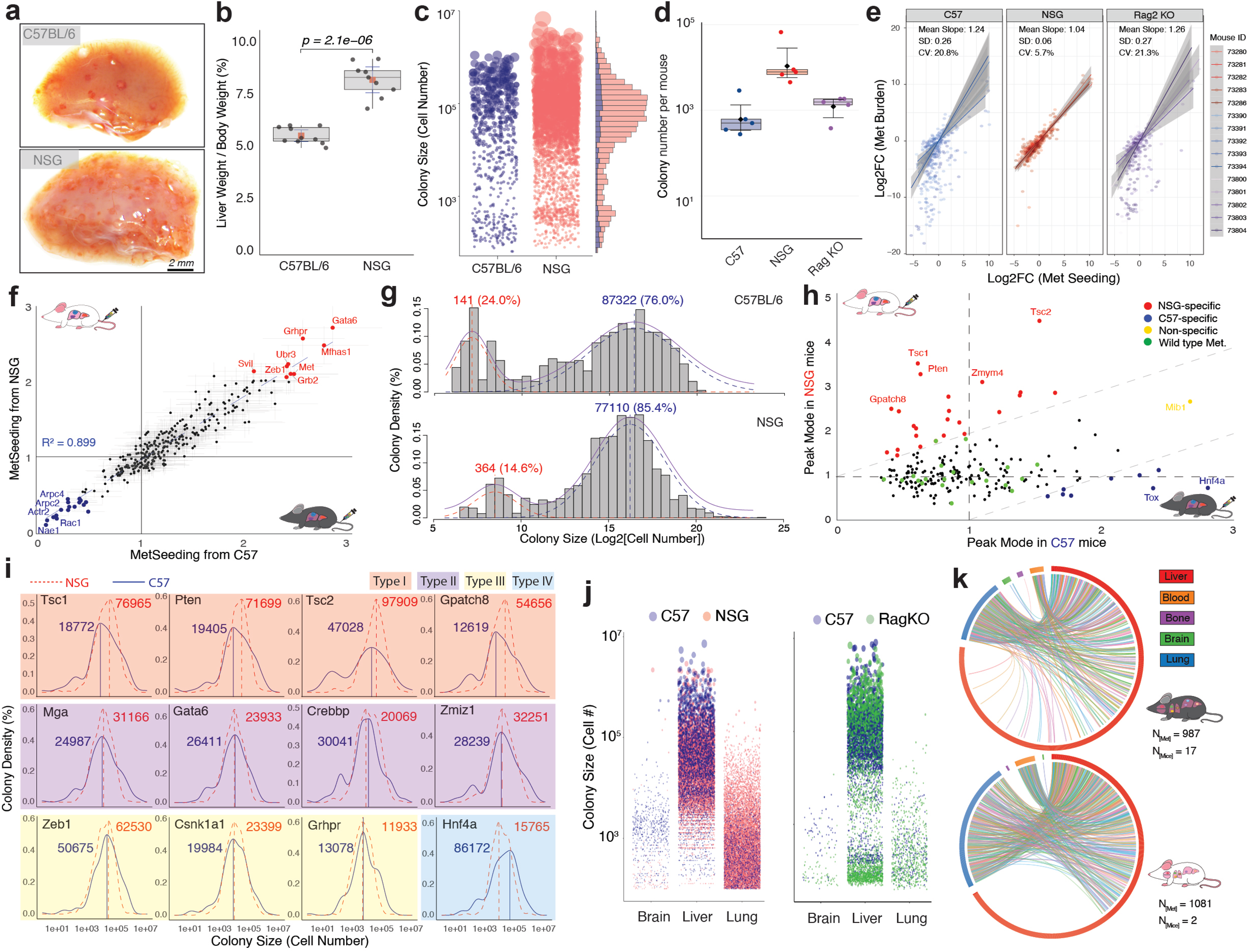
MOBA-seq identifies key factors of immune surveillance in SCLC metastasis. **a.** NSG mice have a higher SCLC liver metastatic burden compared to C57BL/6 mice 3 weeks after tail-vein transplantation. Representative brightfield images of the left lateral lobe are shown. Scale bar: 2 mm **b.** NSG mice have a significantly higher liver weight-to-body weight ratio compared to C57BL/6 mice 3 weeks after tail-vein SCLC transplantation. Each dot is a mouse (n = 10 mice). **c.** MOBA-seq analysis identified more liver metastatic colonies from NSG mice compared to C57BL/6 mice. Each dot represents uniquely barcoded wild-type metastatic colony. Bar graph on the right shows absolute numbers of colonies binned by size. **d.** The innate immune system suppresses SCLC liver metastatic seeding. This box plot shows the number of unique metastatic colonies detected per mouse for C57BL/6, NSG, and Rag2-KO mice. Each dot is a mouse (n = 5 mice). Note that Rag2-KO mice generate only slightly higher metastatic colonies than C57BL/6 mice. **e.** Immune surveillance of SCLC liver metastasis is less dependent on B and T cell activities. Plot of genotype-specific liver metastatic seeding and metastatic burden effects for each mouse genotype in each mouse. Note that Rag2-KO mice are more like C57BL/6 mice. **f.** Immune surveillance broadly suppresses metastatic seeding. For each genotype, the fold change of metastatic seeding compared to wild-type SCLC from both C57BL/6 and NSG mice is plotted. Note there is a high correlation between the two mouse backgrounds (R² = 0.89). **g.** Immune surveillance impacts both dormancy escape and clonal expansion. A colony size histogram showing a bimodal distribution of wild-type liver metastases in both C57BL/6 and NSG mice. C57BL/6 mice have more dormant colonies but also a higher peak mode of exponential growth phase colonies. **h.** MOBA-seq defines immune-sensitive genotypes in their clonal expansion. For each genotype, the fold change of peak mode compared to wild-type SCLC from both C57BL/6 and NSG mice is plotted. Genotypes that grow better in NSG and C57BL/6 mice are highlighted in red and blue, respectively. Wild-type metastases are labeled in green. **i.** MOBA-seq defines four patterns of genotype-specific immune response. Type I genes increase dormancy and reduce clonal expansion; Type II genes increase dormancy without significant change in clonal expansion; Type III genes are resistant to immune response; and Type IV genes increase clonal expansion with the help of the immune system. **j.** Tissue-specific immune response in SCLC metastatic expansion. Comparing metastatic colonies between NSG and C57BL/6 mice (*Left*) and Rag2-KO and C57BL/6 mice (*Right*) across brain, liver, and lung at three weeks after tail-vein transplantation. Immune surveillance on metastatic clonal expansion is tissue dependent. In the liver, the immune systems have little effect on clonal expansion. In the lung, the immune system suppresses clonal expansion, mainly through the innate immune system. In the brain, the adaptive immune system promoted clonal expansion. **k.** Immune-dependent restriction of cross-tissue metastatic dissemination revealed by barcode lineage tracing. Chord diagrams depict shared colonies as connections between liver, blood, bone marrow, brain, and lung in C57BL/6 (*Top*) and NSG (*Bottom*) mice. Chord width is proportional to the log10-transformed cell number of the corresponding colony in each tissue, whereas the outer ring represents all detected colonies within each tissue, with segment width scaled by log10 cell number. The reduced extent of cross-tissue clonal sharing in immunocompetent C57BL/6 mice indicates immune-mediated restriction of secondary metastatic dissemination.

### Genotype-specific responses to immune surveillance in metastatic clonal expansion and dormancy

To identify genotype-specific metastasis in response to immune surveillance, we transplanted the RP48 MOBA500 cell library into both NSG mice and C57BL/6 mice (**Figure S7a-c**). Because metastatic seeding is predominantly constrained by innate immunity, although the absolute number of metastatic colonies differed markedly between NSG and C57BL/6 mice liver, we observed no major genotype-specific divergence in metastatic seeding. Relative metastatic seeding of each genotype in NSG mice and C57BL/6 mice is highly correlated between the two immune contexts (**Figure 3f**, R^2^=0.899), consistent with the prior notion that adaptive immunity has a limited impact in the initial phase of SCLC metastatic seeding. By leveraging the multidimensional resolution of MOBA-seq, we uncovered immune-dependent differences in a subset of metastatic metrics. Specifically, wild-type liver metastases in C57BL/6 mice exhibited a significantly higher dormancy fraction and surprisingly an even slightly larger PeakMode colony size compared to NSG mice (**Figure 3g**). These data suggest that the immune system primarily acts in the early stages of the metastatic cascade to inhibit colony seeding and promote dormancy during early stages, while exerting minimal pressure on the outgrowth of established metastatic clones. We next examined genotype-specific responses to immune surveillance in metastatic clonal expansion (**Figure S7d-e**). We identified genotypes that are sensitive to immune surveillance, characterized by the colonies harboring loss of known tumor suppressors, including *Tsc1*, *Tsc2*, and *Pten* (**Figure 3h**). These results align with prior reports linking *TSC1/TSC2* mutations to increased immunotherapy responsiveness^39^. Intriguingly, we also identified sgRNAs targeting stromal regulators, such as *Tox* (a T cell exhaustion factor) and *Hnf4a* (a liver-specific transcription factor). Inactivation of those genes paradoxically promoted clonal expansion in immune-competent hosts, suggesting potential non-cell autonomous effects of the immune microenvironment. To classify immune-related genotype responses, we compared clonal expansion and dormancy across both immune contexts and defined four response archetypes. The inactivation of Type I genes was associated with a significant reduction in PeakMode colony size and an increase in the percentage of dormant colonies in the immunocompetent hosts, marking tumors harboring mutations in these genes as strong immunotherapy candidates. Type I genes include well known tumor suppressors, such as *Tsc1*, *Tsc2*, and *Pten*. The inactivation of Type II genes exhibited increased dormancy without changes in clonal expansion, identifying these genes as regulators of immune-modulated dormancy programs. This group of genes comprises both established and previously unrecognized metastatic suppressors whose loss enhances metastatic seeding, including *Gata6* and *Crebbp*. The inactivation of Type III genes remained unaffected by immune status, indicating immune evasion or resistance. The inactivation of Type IV genes paradoxically showed better clonal growth in immunocompetent mice, potentially implicating them in the regulation of stromal interactions in metastatic progression (**Figure 3i**). Together, these results illustrate how MOBA-seq enables high-resolution dissection of genotype-specific responses to immune surveillance and highlight key candidates for immunotherapeutic targeting in metastatic SCLC.

### Immune surveillance of SCLC metastasis is tissue-context dependent

Recent single-cell profiling studies have identified tissue-specific immune microenvironments across cancer types^40^. To dissect the contribution of innate and adaptive immunity to tissue-specific metastatic progression in SCLC, we performed MOBA-seq in immunocompetent C57BL/6 mice and immune-deficient NSG and Rag2-KO mice. This analysis uncovered striking tissue-specific immune responses across the liver, lung, and brain. In the liver, C57BL/6 mice exhibited a marked reduction in metastatic colony number compared to NSG mice, while PeakMode colony sizes remained unchanged, suggesting that liver immune surveillance primarily restricts metastatic seeding. In contrast, lung metastases in C57BL/6 mice showed a profound reduction in both colony number and size relative to NSG mice, with Rag2-KO mice exhibiting intermediate reductions in colony size (**Figure 3j left**). These results point to a critical role for innate immunity in constraining both seeding and clonal expansion in the lung metastatic microenvironment. Surprisingly, we noticed a significant increase in both the number and size of tumor colonies in the brain of the C57 mice as compared to both NSG mice and Rag2-KO mice, indicating that the presence of an intact adaptive immune system in the brain may promote metastatic outgrowth (**Figure 3j right**). This immune-dependent pattern of brain metastasis was further supported by our validation of a brain-specific metastasis suppressor *Gpatch8*, whose suppressive effect was observed in immunocompetent C57BL/6 mice but was lost in immunodeficient NSG mice, likely depending on immune-mediated regulation of metastatic colony expansion in the brain TME (**Figure S8a-b**). Finally, immune-mediated suppression of metastasis was also evident in the cross-tissue dissemination of SuperMets. Using liver colonies as the reference population, lineage tracing based on shared barcodes across tissues within individual mice revealed an immune-dependent constraint on secondary metastatic dissemination, with substantially fewer clonally related metastatic populations detected across multiple organs in immunocompetent than in immunodeficient mice. By contrast, loss of specific metastatic suppressors, including *Crebbp*, *Gpatch8*, and *Pten*, increased the frequency of SuperMets shared across multiple tissues. These findings suggest that loss of metastatic suppressors can enable disseminated clones to overcome immune-mediated constraints on systemic metastatic spread. (**Figure 3k, Figure S9a-b**).

Sex, as an important biological variable, profoundly influences immune responses in cancer and other diseases^41^. Leveraging the high-resolution capabilities of MOBA-seq across >5,000 wild-type metastatic colonies, we quantified sex-specific immune response in SCLC liver metastasis by transplanting male-derived RP48 and female-derived RP116 cells into both male and female recipient mice from immune-proficient (*C57BL/6*) and immune-deficient (*NSG*) backgrounds. In the liver, we observed that male RP48 and female RP116 cells exhibited distinct metastatic growth phenotypes; however, because only one cell line of each sex was examined, these differences cannot be attributed specifically to tumor-cell sex. In contrast, female recipient mice consistently showed stronger suppression of metastatic seeding and a higher dormant fraction than male recipients (**Figure S10a-d**). These results are consistent with previous findings that female immune systems are generally more suppressive than those of males and underscore the importance of accounting for sex in mechanistic studies of metastasis and immune surveillance in cancer research^42,43^. Taken together, our high-resolution clonal analysis via MOBA-seq reveals that the impact of immune surveillance on SCLC metastasis is highly tissue-contextual and shaped by both innate and adaptive immune compartments.

### CREBBP is a bona fide suppressor of SCLC liver metastasis

To validate candidate genetic drivers of SCLC metastasis, we selected 30 genes for a targeted MOBA-seq screen. A custom CRISPR library comprising three sgRNAs per gene was transduced into two syngeneic SCLC cell lines, RP48 (male origin) and RP116 (female origin), each tagged with a unique four-nucleotide static barcode. The pooled cell libraries were injected into cohorts of five male and five female recipient mice from both NSG and C57BL/6 backgrounds. Barcode analysis at three weeks post-transplantation revealed a significant enrichment of *Crebbp*-deficient metastases in the liver relative to their pre-transplant representation (**Figure 4a, Figure S11a**). Cell line-specific barcode analysis confirmed that all three independent sgRNAs targeting *Crebbp* consistently and significantly increased liver metastatic burden in both murine cell lines (**Figure 4b**). Temporal profiling at one and three weeks demonstrated that *Crebbp* loss enhanced metastatic efficiency across tissues, sexes, and host genotypes (**Figure 4c, Figure S11b-h**). Colony size distribution analysis at three weeks further revealed that *Crebbp*-deficient metastases displayed both an increased PeakMode of expanding colonies and a higher incidence of dormant colonies (**Figure 4d**), consistent with our prior findings in *Nfib*-deficient clones, where clonal expansion and dormancy are inversely linked during SCLC liver colonization (**Figure 1i**). Notably, *Crebbp*-deficient colonies more frequently gave rise to circulating metastatic populations, as reflected by increased SuperMet scores (**Figure 4e**). Consistent with this finding, Crebbp loss was associated with broadly increased metastatic seeding across the liver, lung, brain, and bone marrow in both NSG and C57BL/6 mice (**Figure S12a-d**). Epigenetic regulation emerged as a dominant axis of metastatic control: among 13 commonly altered SCLC-associated chromatin modifiers, loss of *Crebbp* consistently promoted metastasis across liver, lung, and brain (**Figure 4f**). Comparative analyses across murine and human SCLC cells demonstrated that CREBBP is a conserved metastatic suppressor across species (**Figure 4g**). Together, these data validate CREBBP as a potent and conserved suppressor of SCLC metastasis.

**Figure 4.**
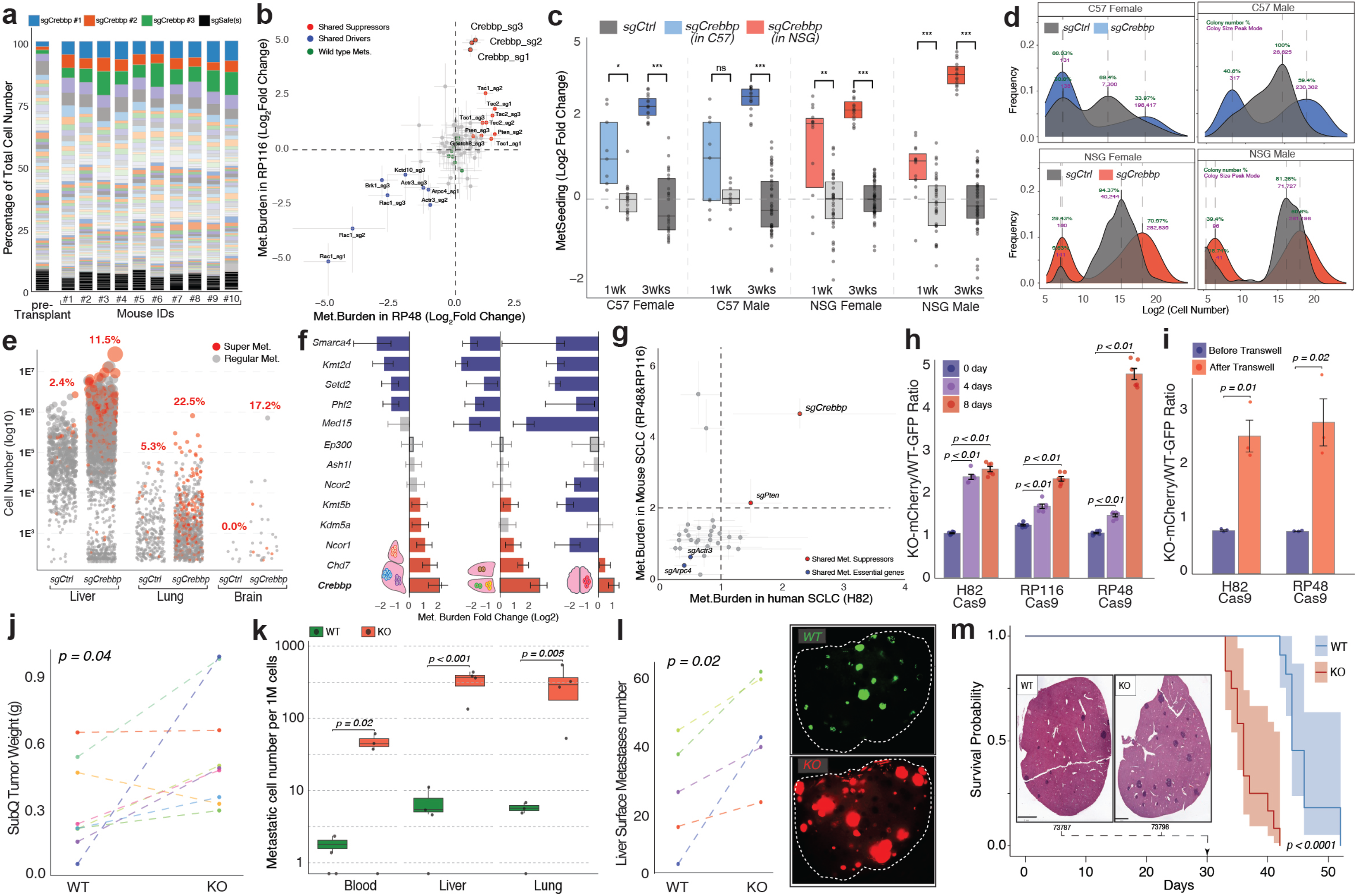
Loss of CREBBP drives a pro-metastatic phenotype *in vitro* and *in vivo*. **a.** *Crebbp* loss markedly enhances SCLC liver metastases with high consistency across recipient mice and sgRNA. Two syngeneic SCLC cell lines transduced with the MOBA-validation library were co-transplanted into ten recipient NSG mice. Shown is the representation of SCLC genotypes recovered from liver tissue of ten mice after three weeks of growth, compared with the pre-transplantation sample. Three *sgCrebbp* genotypes are highlighted. **b.** *Crebbp* loss increases metastatic burden in two syngeneic SCLC models. Tumor burden from the MOBA-validation screen showing relative metastatic burden of all genotypes normalized to wild type SCLC across two syngeneic SCLC cell lines. **c.** *Crebbp* loss enhances SCLC metastatic seeding across mouse genotypes, sexes, and time points. Metastatic seeding analysis of *sgCtrl* (gray) and *sgCrebbp* (colored) colonies in RP116 metastases after one week (left) and three weeks (right), stratified by mouse genotype and sex. Each dot is a mouse. **d.** *Crebbp* loss increases metastatic colony size and alters immune sensitivity. Distribution of colony sizes from the MOBA-validation screen for *sgCtrl* (gray) and *sgCrebbp* (colored) colonies in RP116 metastases after three weeks of growth. Note that *sgCrebbp* colonies exhibit a higher PeakMode together with an increased dormant fraction. **e.** *Crebbp* loss promotes more frequent CTC dissemination from metastatic colonies. Shown are all metastatic colonies in NSG mice for *sgCtrl* and *sgCrebbp* genotypes. SuperMet colonies with barcodes detectable in blood cells are highlighted in red. The percentage of SuperMet colonies for each genotype is indicated above each plot. **f.** *Crebbp* loss increases metastatic burden across multiple organs compared with inactivation of other epigenetic regulators. Bar graph showing relative fold change in metastatic burden for each genotype in liver, lung, and brain after three weeks. Notably, loss of *Crebbp*, but not its paralog *Ep300*, promoted multi-organ metastasis. **g.** *Crebbp* loss increases liver metastatic burden in both human and mouse SCLC models. Metastatic burden from the MOBA-validation screen of all genotypes in human (H82) and murine (RP48 and RP116) SCLC cell lines. Crebbp and Pten are the only metastasis suppressors conserved across species. **h.** *Crebbp* loss increases SCLC *in vitro* growth rate. Three Cas9-expressing SCLC cell lines, murine RP48 and RP116, and human H82, were transduced with GFP-*sgControl* or mCherry-*sgCrebbp* constructs. Equal numbers of GFP (WT) and mCherry (KO) cells were plated at Day 0. The relative proportions of GFP and mCherry cells were quantified by flow cytometry over eight days, and the ratio of mCherry/GFP cells was plotted. n=5 wells **i.** *Crebbp* loss enhances SCLC *in vitro* migration. Equal numbers of GFP-WT and mCherry-KO H82 and RP48 cells were seeded in the upper chamber of a transwell assay. After 48 hrs, the relative proportions of GFP and mCherry cells that migrated to the lower chamber were quantified by flow cytometry. The ratio of mCherry/GFP cells was plotted. n=3 wells **j.** *Crebbp* loss increases subcutaneous SCLC tumor growth. Equal numbers of GFP-WT and mCherry-KO RP48 cells were co-injected into opposite flanks of the same NSG mouse. Tumors were collected after four weeks, and tumor weights were quantified. n=9 mice **k.** *Crebbp* loss increases dissemination of SCLC cells to distant organs. GFP-WT and mCherry-KO RP48 cells were co-transplanted subcutaneously into NSG mice at a 2:1 ratio to achieve comparable primary tumor sizes at endpoint. After four weeks, liver, lung, and whole blood were collected, and the numbers of GFPD and mCherryD SCLC cells per million cell events were quantified by flow cytometry. n=4 mice **l.** *Crebbp* loss increases SCLC liver metastasis. Equal numbers of GFP-WT and mCherry-KO RP48 cells were co-injected into the tail vein of recipient NSG mice and allowed to metastasize for three weeks. Livers were harvested, and tumor colonies were visualized by fluorescence imaging in GFP and mCherry channels. A representative liver image was shown. n=5 mice **m.** *Crebbp* loss decreases survival following SCLC transplantation. C57BL/6 mice were subcutaneously transplanted with *Crebbp-*WT or *Crebbp-*KO RP48 cells. Kaplan–Meier survival curves show reduced survival in mice bearing Crebbp-deficient tumors. Representative H&E staining of liver tissue at day 30 illustrates metastatic seeding. n =13 mice

To further validate CREBBP loss as a driver of the pro-metastatic phenotype in SCLC, we established color-coded co-cultures of GFP-labeled wild-type (WT-GFP) and mCherry-labeled CREBBP-knockout (KO-mCherry) cells in both human (H82) and murine (RP48 and RP116) models. Beginning with equal proportions, KO-mCherry cells progressively outcompeted WT-GFP cells over time, demonstrating a clonal expansion advantage conferred by CREBBP loss (**Figure 4h**). Transwell migration assays of these mixed cultures further revealed a higher proportion of KO-mCherry cells migrating through the membrane after 48 hours, indicating enhanced migratory capacity in both human and murine CREBBP-deficient SCLC cells (**Figure 4i**).

We next subcutaneously transplanted equal numbers of WT-GFP and KO-mCherry RP48 cells into opposite flanks of the same NSG recipient mice. After four weeks, tumors derived from KO cells were significantly larger than those from WT cells (**Figure 4j**). To control for primary tumor size differences, we also performed subcutaneous injections with a 2:1 ratio of WT-GFP to KO-mCherry RP48 cells, yielding comparable tumor sizes at four weeks. Subsequent analysis of liver, lung, and blood samples from these mice revealed increased dissemination of mCherry-KO SCLC cells across all tissues examined (**Figure 4k**). Furthermore, tail vein co-injection of equal number of WT-GFP and KO-mCherry RP48 cells into NSG mice led to a higher proportion of mCherry-positive metastatic lesions on liver surfaces compared to GFP-positive lesions (**Figure 4l**). Consistently, recipient C57BL/6 mice bearing WT RP48 metastases exhibited significantly fewer liver metastases with prolonged survival relative to those receiving KO metastases (**Figure 4m**). Collectively, these data establish CREBBP as a potent suppressor of SCLC liver metastasis.

### CREBBP regulates SCLC cell metastatic changes via regulating CDX2 expression

To investigate CREBBP-mediated cell state regulation during metastasis, we transplanted *Crebbp*-WT and *Crebbp*-KO RP48 cells via tail vein injection into C57BL/6 mice and isolated liver metastases for single-nucleus RNA sequencing at four weeks^44^. From ∼55,000 profiled nuclei, we identified four major clusters of SCLC cells (**Figure 5a, Figure S14a**). *Crebbp*-KO metastases were dominated by cluster 4 SCLC cells, which exhibited strong enrichment of MYC signaling (**Figure 5b-c**). Comparative signature analysis revealed that Crebbp loss drove a loss of neuroendocrine identity and adoption of a more proliferative state (**Figure 5d**).

**Figure 5.**
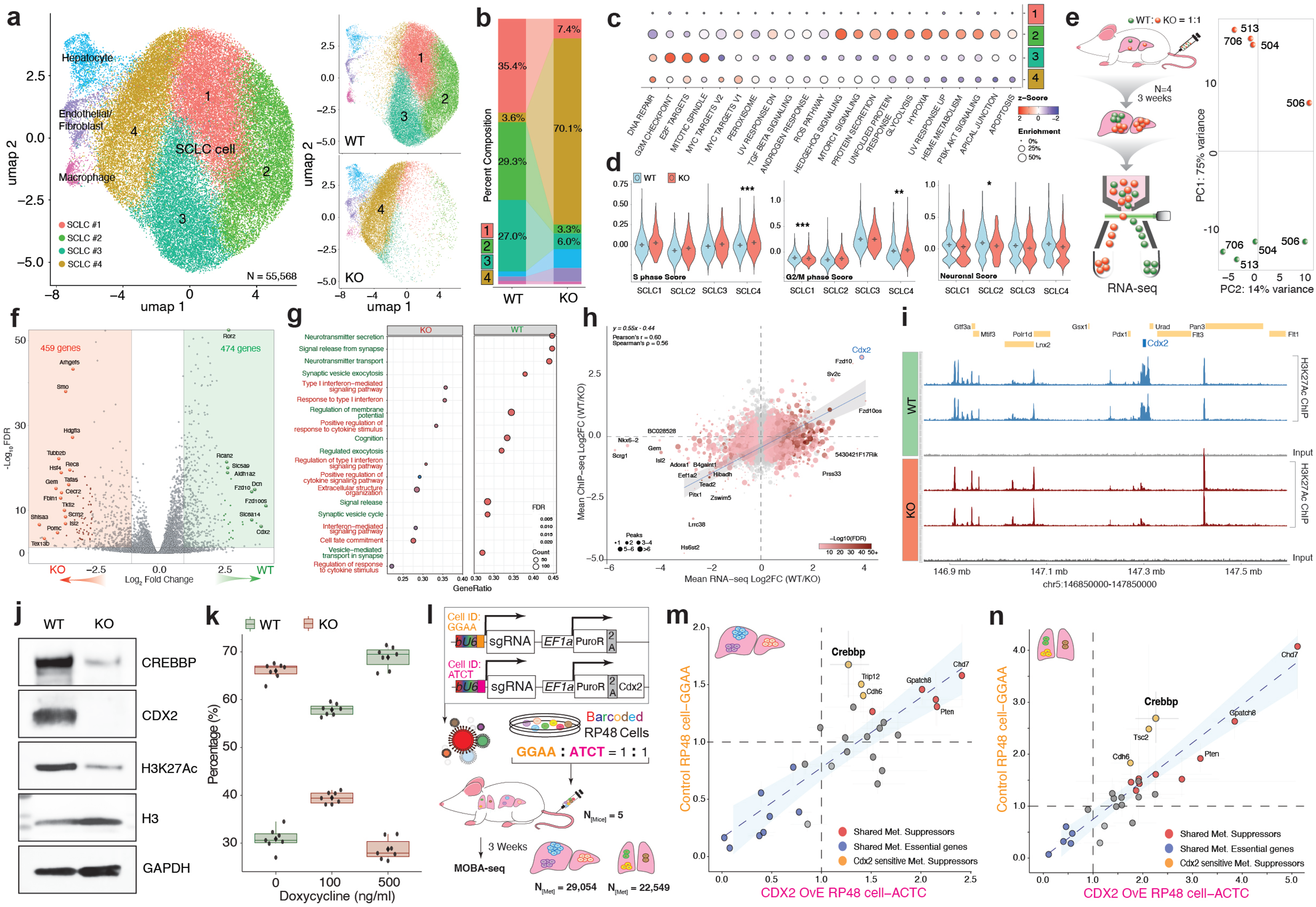
CREBBP suppresses SCLC metastasis through regulating Cdx2 expression. **a.** UMAP of single-nucleus RNA-seq from *Crebbp-*WT and *Crebbp-*KO liver metastases showing four Ascll SCLC clusters. *Crebbp-*KO metastases are enriched in cluster 4 SCLC. Left, UMAP for combined samples with all identified cell types. A total of 55,558 nuclei were analyzed. Right-Top, UMAP for SCLC clusters identified in *Crebbp-*WT sample. Right-Bottom, UMAP for SCLC clusters identified in *Crebbp*-KO sample. **b.** Alluvial plot showing cluster composition of *Ascll* □ SCLC cells from *Crebbp-*WT and *Crebbp-*KO liver metastases. *Crebbp-*KO samples are dominated by Cluster 4 SCLC cells, comprising over 70% of the population. **c.** Cluster 4 SCLC displays MYC signaling activation. Gene set enrichment analysis (GSEA) of *Ascll □* SCLC clusters, with circle color indicating enrichment z-score and size reflecting the percentage of genes enriched in each pathway. **d.** *Crebbp-*KO SCLC cells exhibit increased proliferation and loss of neuroendocrine identity. Violin plots showing pairwise comparisons of S– and G2/M-phase scores and neuronal scores across the four SCLC clusters in *Crebbp-*WT and *Crebbp-*KO metastases. **e.** Schematic of co-transplantation experiment for bulk RNA-seq. GFP-labeled *Crebbp-*WT and mCherry-labeled *Crebbp-*KO RP48 cells were mixed at a 1:1 ratio and transplanted via tail vein injection. After 3 weeks, liver metastases were dissociated, and GFPD and mCherryD SCLC cells were isolated by FACS for bulk RNA-seq analysis. PCA shows clear separation between WT and KO samples. **f.** Volcano plot showing differentially expressed genes between *Crebbp-*WT and *Crebbp*-KO metastatic SCLC cells. A total of 459 genes were upregulated (>2-fold) in *Crebbp-*KO cells, while 474 genes were upregulated (>2-fold) in *Crebbp-*WT cells, indicating balanced transcriptional activation and repression upon *Crebbp* loss. **g.** GSEA analysis of bulk RNA-seq from sorted *Crebbp-*WT and *Crebbp-*KO SCLC cells showing loss of neuroendocrine programs and activation of type I interferon signaling in *Crebbp-*KO cells. Circle color represents enrichment FDR, and circle size reflects the number of genes contributing to each pathway. **h.** Integration of H3K27ac ChIP-seq and RNA-seq identifies *Cdx2* as a CREBBP-dependent epigenetic target. Differential H3K27ac peak signals between *Crebbp-*WT and *Crebbp*-KO SCLC cells (two biological replicates) were cross-compared with differential gene expression from RNA-seq. Circle color represents –login DFDR, and circle size reflects the total number of differential peaks per gene. *Cdx2* is among the most significantly enriched and shared targets in WT cells compared with KO cells. **i.** *Crebbp* loss specifically reduces H3K27ac modification at the *Cdx2* locus. H3K27ac ChIP-seq tracks from *Crebbp-*WT and *Crebbp-*KO SCLC cells are shown for the indicated chromosome 5 region. *Crebbp-*KO cells display a selective loss of H3K27ac at the *Cdx2* locus, while neighboring gene loci remain unaffected. **j.** *Crebbp l*oss diminished CDX2 protein expression in SCLC cells. Western blot analysis of CREBBP, CDX2, and H3K27ac in *Crebbp-*WT and *Crebbp-*KO RP48 cells. Total H3 and GAPDH were used as controls. **k.** Doxycycline-induced CDX2 re-expression in *Crebbp-*KO RP48 cells suppresses proliferation and growth advantage in co-culture with *Crebbp-*WT cells. *Crebbp-*WT-TetON-CDX2 and *Crebbp-*KO-TetON-CDX2 RP48 cells were mixed at a 1:1 ratio and treated with 0, 100, or 500 ng/mL doxycycline in co-culture for 6 days. GFP/mCherry cell ratios were quantified by flow cytometry. n= 7 wells **l.** Schematic of the MOBA-seq screen using control or CDX2-overexpressing RP48 cells. Lentiviral sgRNA libraries were constructed using control (PuroR) or CDX2-overexpression (PuroR-2A-CDX2) backbones. 4N-barcoded control and CDX2-overexpressing RP48 cell libraries were mixed at a 1:1 ratio and transplanted via tail vein into five NSG mice. Three weeks post-transplantation, metastasis-bearing livers and lungs were collected, and over 20,000 barcoded metastases were identified from both tissues for MOBA-seq analysis. **m.** Crebbp-mediated metastatic suppression in the liver is CDX2-dependent. MOBA-seq analysis deconvoluted control and CDX2-overexpressing RP48 cells using their 4N-static barcodes. Genotype-specific metastatic burden was calculated as fold change relative to the mean of all control colonies (*sgCtrl*). The blue regression line with confidence interval represents genotypes unaffected by CDX2 expression, while those preferentially suppressed under CDX2-overexpressing conditions are highlighted in yellow. *Crebbp* inactivation increased liver metastatic burden only in control RP48 cells, but not in CDX2-overexpressing RP48 cells. **n.** *Crebbp*-mediated metastatic suppression in the lung is CDX2-dependent. MOBA-seq analysis of genotype-specific metastatic burden in control and CDX2-overexpressing RP48 cells that metastasized to the lung as described in m. *Crebbp* inactivation markedly increased lung metastatic burden in control RP48 cells but had a reduced effect in CDX2-overexpressing RP48 cells.

To determine whether *Crebbp* loss alters SCLC subtypes, we applied consensus non-negative matrix factorization (cNMF) to transcriptional profiles of *Crebbp*-WT and *Crebbp*-KO cells. Based on solution stability and reconstruction error, we resolved four recurrent gene-expression programs (GEPs) (**Figure S13a-b**). Comparison with canonical SCLC subtype markers revealed that GEP1 was enriched for the neuroendocrine SCLC-A marker Ascl1, whereas GEP3 was strongly enriched for SCLC-I-associated mesenchymal genes, including *Vim*, *Axl* and *Zeb2* (**Figure S13c**). Notably, *Crebbp*-KO cells showed increased usage of GEP3 and GEP4, together with reduced usage of GEP2, consistent with acquisition of a more SCLC-I-like, non-neuroendocrine state. These findings suggest that *Crebbp* loss promotes metastatic fitness in part by reshaping tumor-cell state associated with SCLC-I (**Figure S13d**). To compare transcriptional profiles in a shared *in vivo* environment, we co-transplanted GFP-labeled WT and mCherry-labeled *Crebbp-KO* cells into the same mice (as described in **Figure 4k**). Metastatic SCLC cells were subsequently sorted by fluorescence and analyzed by bulk RNA-seq (**Figure 5e**). Differential expression analysis identified a comparable number of genes up– and downregulated in *Crebbp-KO* cells (**Figure 5f, Figure S14b-c**), consistent with CREBBP functioning as both a transcriptional activator and repressor in cancer^45^. GSEA confirmed loss of neuroendocrine programs in Crebbp-deficient cells and unexpectedly revealed a robust activation of type I interferon signaling (**Figure 5g**). Together, these findings indicate that Crebbp suppresses SCLC metastasis by constraining both intrinsic cancer cell plasticity and microenvironmental signaling responses.

CREBBP functions as a lysine acetyltransferase and epigenetic regulator. To identify CREBBP-dependent epigenetic targets in SCLC, we performed H3K27ac ChIP-seq in Crebbp-WT and Crebbp-KO RP48 cells and integrated this data with RNA-seq profiles (**Figure S14d**). This analysis identified Caudal Type Homeobox 2 (*Cdx2*) as a shared target across both datasets (**Figure 5h**). Cdx2 is an intestine-specific transcription factor known to function as a tumor suppressor in colorectal cancer progression and metastasis^46,47^. *Crebbp* loss selectively reduced H3K27ac occupancy at the Cdx2 locus, accompanied by decreased *Cdx2* transcript abundance and CDX2 protein expression (**Figure 5i-j**). Doxycycline-induced re-expression of *Cdx2* in Crebbp-KO RP48 cells suppressed proliferation and migration in co-culture competition assays relative to Crebbp-WT controls (**Figure 5k, Figure S14e-f**). To quantify genetic epistasis between *Crebbp* and *Cdx2* in SCLC metastasis, we barcoded control and CDX2-overexpressing RP48 cells and introduced an additional 30-gene perturbation library using the MOBA-seq backbone. Three weeks after tail vein transplantation into NSG mice, over 20,000 barcoded liver and lung metastases were recovered (**Figure 5l**). Comparative analysis of metastatic burden revealed that most gene perturbations exerted similar effects across control and *Cdx2*-overexpressing backgrounds. In contrast, inactivation of Crebbp markedly increased liver and lung metastatic burden in control cells but not in *Cdx2*-overexpressing cells (**Figure 5m-n, Figure S14g-h**). These results establish *Cdx2* as a CREBBP-dependent epigenetically regulated target and functional effector whose repression via epigenetic modification underlies the pro-metastatic phenotype of Crebbp-deficient SCLC.

### CREBBP loss remodels the liver microenvironment by modulating endothelial–immune crosstalk

Our prior results showed that *Crebbp* inactivation promotes metastatic seeding while simultaneously increasing the fraction of dormant colonies (**Figure 3i**). Transcriptomic profiling further revealed activation of the type I interferon pathway in Crebbp-KO cells, implicating a pro-inflammatory immune response within the Crebbp-deficient metastatic TME (**Figure 5g**). To spatially resolve these effects, we applied the Xenium platform that enables spatial mapping of 5100 genes to formalin-fixed, paraffin-embedded (FFPE) sections of Crebbp-WT and Crebbp-KO RP48 liver and lung metastases from C57BL/6 mice. This approach yielded high-resolution expression maps with a median of 1,462 transcripts per cell and transcript densities up to 1,194 per 100 µm² (**Figure 6a, Figure S15a-c**). Using six signature genes of each cluster, we classified 11 major cell types within the liver metastatic niche (**Figure 6b**). Cancer cells engage in close interactions with endothelial cells (ECs) throughout the metastatic cascade^48^. To assess the effect of CREBBP loss in EC dynamics, we performed proximity analysis in liver metastases derived from Crebbp-WT and Crebbp-KO tumors. Crebbp-KO metastases exhibited a marked expansion of EC populations compared with Crebbp-WT counterparts (**Figure 6c**). H&E staining revealed that these expanded EC populations in Crebbp-KO liver metastases displayed pronounced vascular abnormalities, characterized by highly vascularized lesions with irregular and leaky vessel architecture (**Figure 6d, Figure S15d**). Vascular abnormalities during tumor progression are known to increase geometric resistance to blood flow and impair vascular function, leading to reduced perfusion and enhanced hypoxia^49,50^. Consistent with this, SCLC cells in Crebbp-KO liver metastases exhibited hyperactivation of hypoxia response, glucose and lipid metabolism pathways (**Figure 6e, Figure S15e-f**). To systematically identify endothelial cell (EC) state changes associated with Crebbp loss in liver metastases, we performed differential gene expression analysis of ECs within tumor microregional (TMR) ROIs. ECs in Crebbp-KO TMRs showed elevated expression of inflammatory markers, whereas ECs in Crebbp-WT TMRs preferentially expressed stromal and extracellular matrix (ECM) remodeling genes (**Figure 6f**). The endothelium forms the primary interface mediating immune-tumor communication. We therefore performed cell–cell interaction analysis to identify genotype-specific ligand–receptor modules between ECs and immune cells. Crebbp-deficient metastases exhibited pronounced activation of leukocyte recruitment pathways, indicative of enhanced immune cell engagement in the Crebbp-KO microenvironment (**Figure 6g, Figure S15g-h**). This enhanced immune cell infiltration in Crebbp-KO tumors was validated by both spatial TMR analysis and immunofluorescence staining, each revealing increased T and NK cell presence within Crebbp-KO metastases (**Figure 6h-i, Figure S15i-k**). Strikingly, intra-tumoral T cells in Crebbp-KO lesions were enriched for exhaustion markers relative to peritumoral T cells, a pattern absent in WT tumors (**Figure 6j-k**). To validate these observations, we performed flow cytometry on metastasis-bearing livers, which confirmed increased frequencies of PD1^+^ CD8 and CD4 T cells in Crebbp-KO tumors compared with WT controls (**Figure 6l**). The distinct immune remodeling observed in Crebbp-deficient metastases suggested a therapeutic vulnerability to immune checkpoint inhibition. To test this hypothesis, we analyzed clinical outcomes from the Memorial Sloan Kettering–Integrated Mutation Profiling of Actionable Cancer Targets (MSK-IMPACT) cohorts^51^. Across cancer types, patients harboring CREBBP mutations exhibited significantly higher tumor mutational burden (TMB) compared with CREBBP-wild-type counterparts (**Figure S15l**). Importantly, among patients treated with immune checkpoint inhibitors (ICI), those with *CREBBP* mutations demonstrated significantly prolonged overall survival compared with patients harboring *CREBBP*-wild-type tumors (**Figure 6m, Figure S16**). Together, these findings indicate that CREBBP inactivation remodels the SCLC metastatic microenvironment by promoting endothelial-immune cell interactions and driving T cell exhaustion, thereby establishing a hypoxic and immunosuppressive niche that supports metastatic progression and may predict increased responsiveness to immune checkpoint blockade.

**Figure 6.**
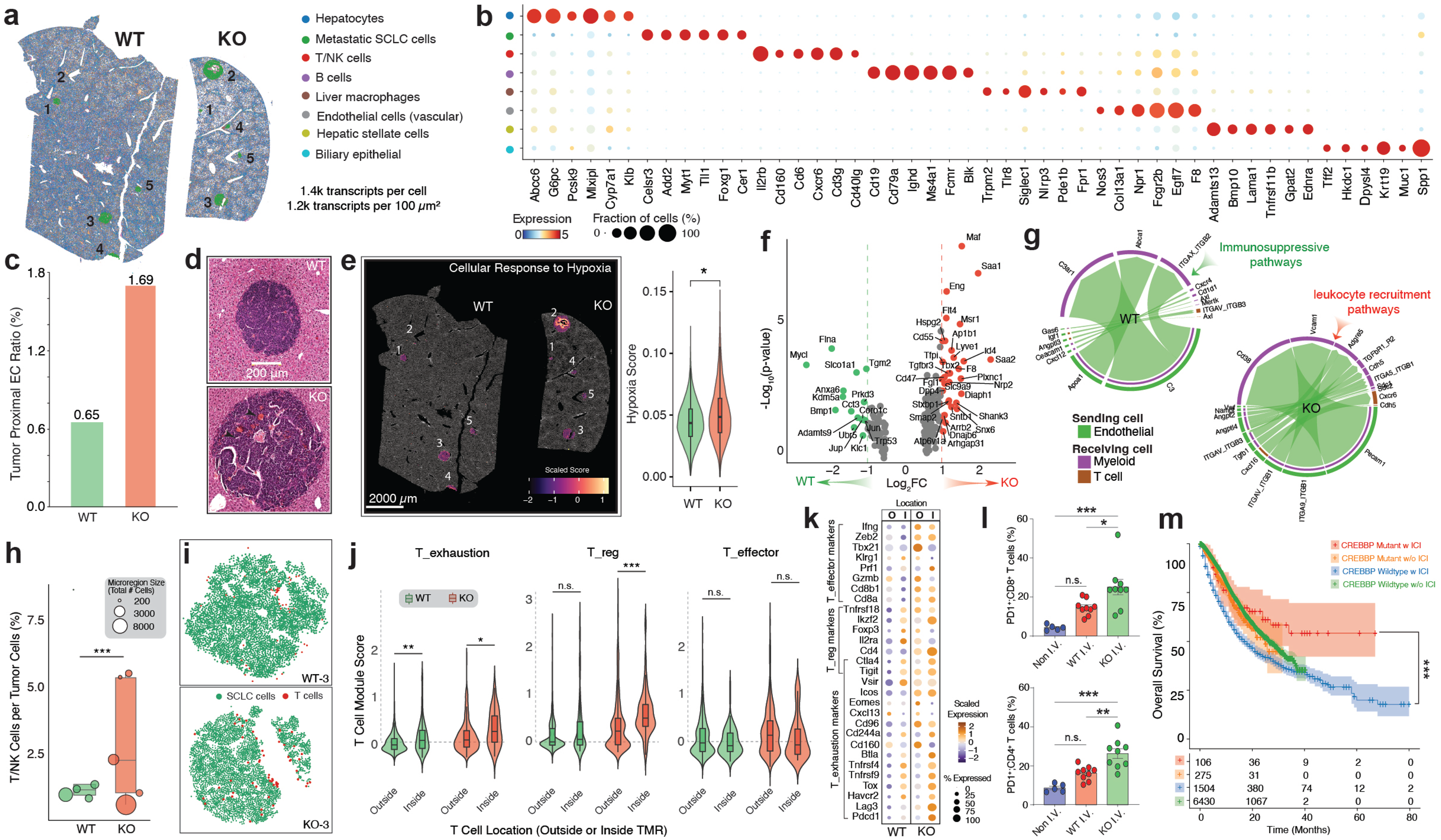
*Crebbp* loss orchestrates the liver metastatic microenvironment through endothelial-immune cell crosstalk. **a.** Spatial transcriptomic profiling of *Crebbp*-WT and *Crebbp*-KO liver metastases. Xenium spatial maps reveal high-resolution cellular architecture, showing SCLC cells and 11 major stromal cell types within metastatic liver tissue. Each cell expresses an average of over 1,400 transcripts. SCLC tumor microregion ROIs (TMRs) are outlined in green and labeled with unique ID numbers. **b.** Dot plots showing signature genes of major cell types within the liver metastatic microenvironment. Circle color indicates scaled (SCTransformed) gene expression, while circle size represents the proportion of cells expressing each gene in the corresponding cell population. **c.** Proximity analysis reveals increased endothelial cell (EC) abundance in *Crebbp*-KO metastases. ECs within 20 µm of tumor cell centroids were defined as tumor-proximal, and their proportion among total ECs was quantified, showing enrichment in *Crebbp*-deficient lesions compared to wild type. **d.** *Crebb*p-KO liver metastases exhibit vascular abnormalities. Representative H&E-stained images of *Crebbp*-WT and *Crebbp*-KO liver metastases show highly vascularized lesions with irregular vessel architecture and red blood cell extravasation (arrows). Scale bar, 200 µm. **e.** *Crebbp*-KO liver metastases activate hypoxia response pathways. Scaled hypoxia scores are shown on Xenium spatial maps for individual *Crebbp*-WT and *Crebbp*-KO metastatic lesions (left) and summarized across all cells within metastatic lesions by violin plot (right). **f.** Differential expression in endothelial cells within tumor microregions (TMRs). Volcano plot showing genes differentially expressed in ECs from TMRs of *Crebbp*-WT versus *Crebbp*-KO liver metastases. Dashed lines indicate significance and effect-size (log2FC) thresholds. ECs in *Crebbp*-KO metastatic TMRs express elevated inflammatory markers, whereas ECs in *Crebbp*-WT TMRs elevated stromal and ECM-remodeling markers. **g.** ECs from *Crebbp*-KO TMR exhibit a pro-inflammatory phenotype. Cell–cell interaction analysis (CellChat) reveals distinct ligand–receptor modules enriched between ECs and immune cells in *Crebbp*-WT and *Crebbp*-KO metastases. WT TMRs show activation of immune-suppressive signaling, whereas KO TMRs show activation of leukocyte recruitment and pro-inflammatory pathways. **h.** *Crebbp*-KO TMRs show increased T and NK cell infiltration. The ratio of T/NK cells to tumor cells is plotted for each TMR from *Crebbp*-WT and *Crebbp*-KO metastases. Dot size corresponds to TMR area. **i.** Spatial proximity of T cells and tumor cells in *Crebbp*-WT and *Crebbp*-KO tumor microregions. Representative Xenium spatial maps show T cells (red) and SCLC tumor cells (green) in paired *Crebbp*-WT and *Crebbp*-KO TMRs of comparable size. *Crebbp*-KO tumors display increased T cell infiltration within the tumor core, whereas T cells in *Crebbp*-WT tumors are primarily localized at the tumor-stroma interface. **j.** T cells within TMRs exhibit enhanced exhaustion and regulatory phenotypes, particularly in *Crebbp*-KO metastases. Violin plots show T cell module scores for exhaustion, regulatory (Treg), and effector programs, comparing T cells located inside versus outside TMRs in *Crebbp*-WT and *Crebbp*-KO metastases. **k.** Intra-tumoral T cells in SCLC metastases display exhaustion-associated gene signatures. Marker genes defining effector, regulatory, and exhausted T cell states are shown for T cells located inside or outside *Crebbp*-WT and *Crebbp*-KO liver metastases. Dot color represents scaled (SCTransformed) expression, and dot size indicates the proportion of T cells expressing each marker. T cells within *Crebbp*-KO tumors show markedly elevated expression of exhaustion markers compared with those outside *Crebbp*-KO lesions. **l.** *Crebbp*-KO liver metastases are associated with increased PD1 □ T cells. Flow cytometry analysis of CD8D and CD4D T cells from control (non-I.V.), *Crebbp*-WT (WT I.V.), and *Crebbp* –KO (KO I.V.) metastasis-bearing livers. Each dot represents one mouse; data are shown as mean ± SD. n.s., not significant; *p < 0.05; **p< 0.01; **\***p < 0.001. **m.** Kaplan–Meier analysis of overall survival in the MSK-IMPACT cohort stratified by *CREBBP* mutation status. Patients with *CREBBP*-mutant tumors show improved survival following immune checkpoint inhibitor treatment compared with those harboring *CREBBP* wild-type tumors.

## Discussion

Metastasis is a complex, multistep process shaped by the interplay between tumor-intrinsic programs and microenvironmental cues. Yet, how individual factors contribute to each stage of this cascade remains poorly defined. Here, we developed the MOBA-seq technology for high-accuracy, scalable analysis of genetic drivers of metastasis at single-colony resolution. Using SCLC as a model, we redefined and quantitatively measured the role of the canonical driver *Nfib* in metastatic seeding, dormancy escape, and clonal expansion. Screening over 400 recurrently mutated SCLC genes across liver, lung, and brain revealed metastatic seeding as the dominant determinant of overall burden, with immune surveillance, primarily through innate immunity, acting mainly at this stage. Tissue context and sex further modulated these immune effects. Finally, we identified *Crebbp* as a potent metastatic suppressor whose loss enhances metastatic fitness through Cdx2-dependent transcriptional reprogramming and formation of a T cell-exhausted microenvironment.

MOBA-seq, through heritable DNA barcoding, quantifies tumor colony size at single-cell to 10 cells resolution. In contrast, even ultra–high-resolution MRI detects lesions only above ∼0.1 mm in diameter (∼1,000 cells), making MOBA-seq ∼100-fold more sensitive for identifying micro-metastatic colonies in preclinical cancer models. Its detection limit is mainly determined by sequencing depth and sampling variability during genomic DNA extraction and library preparation, particularly in frequent metastatic tissues. When profiling rare events, such as brain micro-metastases, circulating tumor cells, or dormant disseminated cells, MOBA-seq readily achieves single-cell sensitivity. MOBA-seq depends on single-cell barcoding and clonal outgrowth. Although barcode collisions or multicellular seeding cannot be entirely excluded, they are minimized through low-MOI transduction, single-cell preparation, and computational filtering of high–copy-number barcodes. As sequencing costs continue to fall and analytical methods improve, MOBA-seq represents a scalable platform for quantitative assessment of tumor growth dynamics that could surpass traditional imaging-based assays in cancer research.

With hundreds of thousands of *in vivo* data points, MOBA-seq combined with the LETTUCE computational pipeline enables quantitative analysis of key steps in the metastatic cascade. By integrating CRISPR-based knockout screening, we systematically quantified the loss-of-function effects of hundreds of genes on metastatic seeding, dormancy escape, and clonal expansion. While “metastatic seeding” reflects a dynamic equilibrium among CTC docking, trans-endothelial migration, and immune-mediated clearance, MOBA-seq captures only the net outcome, which is the overall colony number whose barcodes are detectable at the experimental endpoint. Time-course analyses confirmed stable barcode maintenance in SCLC liver metastases but revealed pronounced barcode clearance in lung metastases, indicating tissue-specific selective pressures. Currently, MOBA-seq cannot fully disentangle initial seeding from clonal survival, but integrating temporally resolved lineage-tracing barcodes could address this limitation^52^. In its current design, MOBA-seq focuses on the metastatic cascade within distant organs and is not yet optimized for studying spontaneous metastasis from primary tumors. Nonetheless, our SuperMet metric allows quantification of genotype-specific dissemination, which could be easily extended to primary tumors by tracking shared barcodes in blood and distant organs. To further resolve whether metastatic colonies arise from the same or distinct primary clones, future iterations of MOBA-seq will incorporate dynamic barcoding strategies. Finally, seeding and clonal expansion emerge as the principal determinants of metastatic burden, but their relative contributions vary across organs. In highly vascularized tissues such as liver and lung, intrinsic cancer properties dominate metastatic seeding, whereas tissue context more strongly influences dormancy escape, clonal outgrowth, and immune clearance. In contrast, successful colonization of the brain requires cooperative interactions between cancer-intrinsic drivers and the brain microenvironment, making seeding the rate-limiting step in brain metastasis. These findings establish a framework for understanding how cancer-intrinsic and environmental determinants cooperatively shape the metastatic cascade.

Immune surveillance represents the primary barrier for newly disseminated cancer cells to establish residency in distant organs. By analyzing hundreds of thousands of metastatic colonies in both immune-competent and immune-deficient hosts, we generated a comprehensive blueprint of how and when immune surveillance acts during the metastatic cascade. Our data reveals that immune elimination is most effective during early metastatic seeding and dormancy induction but becomes markedly less efficient once a lesion is established. This may explain why immunotherapies are highly effective in hematologic malignancies where cancer cells exist primarily as single cells, but show limited efficacy in solid tumors with well-formed clones. Intriguingly, highly proliferative tumors tend to harbor larger dormant populations under immune pressure, suggesting a dynamic balance between proliferation and immune evasion that warrants further study. We further demonstrate that the innate immune system, rather than the adaptive arm, plays a dominant role in metastatic suppression, consistent with prior studies highlighting NK cell–mediated control of SCLC liver metastasis^26^. While genotype-specific immune responses also shape clonal expansion, the precise immune cell types and molecular programs governing immune remodeling and escape remain to be defined. Targeting innate immune components such as NK cells may therefore offer a pan-cancer strategy for metastasis prevention. Finally, tissue-specific immune milieus impose distinct selective pressures on metastasis. Notably, successful SCLC brain metastasis appears to require an intact immune microenvironment, potentially by co-opting immune infiltration signals to breach the blood–brain barrier. This insight may guide the development of immune-competent spontaneous brain metastasis models for SCLC and other cancer types. Overall, our findings emphasize that the timing and cellular context of immune engagement are critical determinants of both immune surveillance and therapeutic efficacy.

Both our study and previous work on SCLC metastasis highlight the central role of epigenetic regulators in metastatic adaptation ^11^. This is likely because metastasis is a complex, multistep process in which effective drivers must confer broad advantages across diverse biological programs, including cell migration, proliferation, metabolic adaptation, and immune interaction. Epigenetic modifiers and master transcription factors are particularly suited for this role due to their capacity to modulate broad and diverse downstream targets. Interestingly, while Crebbp functions as a potent metastatic suppressor, its paralog Ep300 exhibits minimal impact in our study. Although the underlying mechanisms remain to be elucidated, we hypothesize that this divergence may arise from their differential regulation of key downstream targets such as Cdx2. Whether additional Crebbp-specific targets contribute to metastatic suppression, and how Cdx2 precisely mediates this regulation in the metastatic cascade, will require further study. Moreover, Crebbp loss drives a phenotypic shift in SCLC from a neuroendocrine to a Myc-high subtype, which is known to exhibit elevated metastatic potential in both mouse models and human patients^53^. Whether this Myc-high phenotype, and the interaction between Crebbp-Cdx2 signaling leading to vascular and immune remodeling and T-cell exhaustion, act as coordinated mechanisms of metastasis remains an open question. Nonetheless, our findings provide a conceptual framework for targeting master regulatory pathways to enable more effective combination therapies for metastatic cancer.

In summary, we have developed MOBA-seq, a high-throughput platform that quantifies genotype-specific effects across the metastatic cascade with single-colony resolution. MOBA-seq is a powerful tool to measure tumor fitness with exceptional accuracy and scalability. Using this approach, our study establishes a conceptual framework in which cancer-intrinsic drivers cooperate with microenvironmental determinants to shape distinct stages of the metastatic cascade. Finally, we identified Crebbp as a master metastatic suppressor in SCLC, offering new mechanistic insight and potential therapeutic avenues for preventing and treating this lethal disease.

## ACKNOWLEDGEMENTS

We dedicate this work to the memory of Gui Zhang, an exceptional collaborator and dear friend who helped build the technical foundation of this study for the Tang lab. We thank the staff at Division of Comparative Medicine (DCM) at Washington University for expert animal care, Anatomic and Molecular Pathology (AMP) Core Labs, Siteman Flow Cytometry Core (SFC), and the WashU Genome Technology Access Center (GTAC) for experimental support. We thank Monte Winslow’s lab at Stanford University for support in the generation of SCLC metastasis mouse models for the initial MOBA-seq experiments. We thank Kate Sutherland for sharing RP48 and RP116 cells. We thank Monte Winslow, Julien Sage, Kate Sutherland, David DeNardo, and Hong Chen for helpful comments. This work was supported by NIH R00-CA256039 (to R.T.), Cancer Research Foundation Young Investigator Award (to R.T.), the Phi Beta Psi Sorority Research Grant (to R.T.), the American Lung Association Innovation Award (to R.T.), U01CA294532 (to L.D., F.C), U54AG075934 (to L.D., F.C), and R01CA260112 (to L.D., F.C). The funders had no role in study design, data collection and analysis, and decision to publish or preparation of the manuscript.

## AUTHOR CONTRIBUTIONS

R.T. conceived the project and designed the experiments. C.R., A.X. and R.T. led experimental data production with contributions from A.V., X.F., C.W.P., X.Q., I.C., M.D., D.A.G., Y.Y., I.G., Z.Y., T.W., A.H.K., A.S., G.L., F.C., and L.D.. A.X., A.V., X.Q., I.C., and R.T. led the data analysis. A.X. and I.C. performed the MOBA-seq analysis. A.V., X.Q., and A.X. performed Xenium data analysis. R.T. oversaw the project. C.R., A.X. and R.T. wrote the manuscript with input from all authors.

## Figure Legends

**Supplementary Figure 1.** Establish the MOBA-seq pipeline. **a.** Schematic of the MOBA-seq construct design, including cloning of the barcoded insert downstream of the U6 promoter and incorporation of sgRNA oligo pools. PCR amplification using flanking restriction sites (XbaI, Xho1) enables efficient library assembly and integration of random barcodes in the bovine U6 promoter. Golden gate cloning enables sgRNA oligo library insertion into barcode vectors. **b.** Flow cytometry validation of Cas9 cutting efficiency in H82, RP48, and RP116 cells. Self-targeting GFP/sgGFP reporters are used to transduce Cas9D and Cas9D cells. Cas9 cutting efficiency is normalized by percentage of GFPD cells in transduced Cas9D cells. **c.** Distribution of total cell counts per barcode illustrating barcode complexity and evenness across the library. The red dashed line marks the 95th percentile threshold used to filter high-abundance barcodes prior to downstream analysis. **d.** Quantification of mCherry-labeled H82 seeding efficiency via different injection routes in liver and lung. Percentage of mCherryD H82 cells in liver and lung are measured 2 hours and 48 hours post-transplantation. Error bars represent mean ± s.d. IC: intra-cardiac; IV: intra-veinous; ISp: intra-splenic. **e.** Overview of the MOBA-seq library amplification workflow. Barcoded genomic DNA from target organ is subjected to MOBA 1^st^-round PCR (20 cycles), column-purified, diluted, and followed by an NGS-compatible 2^nd^-round PCR (15 cycles). **f.** Linear regression between relative NGS read counts and expected barcode frequency for spike-in cells across samples. R²=0.98 demonstrated a robust reproducibility for colony size calculation. **g.** Stacked distribution of R² values from linear regression analyses used to calculate colony sizes for samples derived from bone marrow, liver, lung, and blood. The consistently high R² values across tissues indicate strong confidence and robustness in colony size quantification for diverse sample types. **h.** Flowchart summarizing the LETTUCE computational pipeline for MOBA-seq data analysis, including read merging, filtering, UMI correction, cutoff selection, barcode collapse, and final metastasis-associated statistics analysis. **i.** Histogram of the number of metastatic colonies detected per mouse across experiments in C57BL/6 and NSG backgrounds. Mean colony number per mouse is indicated by dashed blue lines. **j.** Proportion of single-mouse-seeding SCLC clones identified in C57BL/6 and NSG mice following transplantation of barcoded RP48 cells via tail vein injection. The majority of barcoded clones seeded into only one mouse, indicating minimal multi-seeding events from the same barcode and ensuring high clonal assignment fidelity. **k.** Genotype-specific metastatic seeding is reproducible across individual mice. Absolute numbers of liver metastatic seeding events associated with each sgRNA genotype were compared between two biological replicates. Metastatic seeding frequencies were highly concordant between mice (*R*² = 0.94), demonstrating reproducible genotype-specific effects on liver metastatic seeding. **l.** Reproducibility of metastatic seeding profiles across biological replicates. Heatmap showing pairwise *R*² values for genotype-specific metastatic seeding profiles across six individual mice. Pairwise correlations ranged from *R*² = 0.65 to 0.94, demonstrating consistent metastatic seeding patterns across independent biological replicates. **m.** High-sensitivity detection of circulating tumor cells in C57BL/6 and NSG mice. Numbers of unique circulating tumor cell (CTC) colonies detected per milliliter of blood (*Top*) and total CTC cell numbers per milliliter of blood (*Bottom*) are shown for individual mice. Dashed lines indicate the mean across mice. These data demonstrate robust detection of metastatic CTCs in peripheral blood.

**Supplementary Figure 2.** MOBA-seq quantifies tissue-specific metastatic effects of *Nfib*. **a.** Schematic of the MOBA-seq experimental design to quantify tissue-specific metastatic effects of *Nfib*. RP48 SCLC Cas9D cells were transduced with pooled barcoded lenti-sgRNAs targeting either Safe controls or *Nfib*, followed by intravenous injection into NSG mice. After metastatic colonization in liver, lung, and brain, tissues were collected at defined timepoints and processed for barcode-based metastatic clonal reconstruction and metastatic effects analysis. **b.** Stacked bar plots showing genotype-specific metastatic cell numbers across liver and lung at serial timepoints. Each bar depicts the proportional contribution of metastatic SCLC clones within each sgRNA group. Loss of *Nfib* reduced clonal representation in both liver and lung over time. **c.** Quantification of relative metastatic burden for sgSafe and *Nfib*-targeting clones over time in liver and lung. *Nfib* inactivation significantly reduces metastatic burden in liver and lung comparing to sgSafe control. **d.** Quantification of relative metastatic seeding for *sgSafe* and *Nfib*-targeting clones in liver and lung. *Nfib* inactivation led to a sustained reduction in initial seeding efficiency in the liver and progressively decreased lung metastatic colony numbers toward sgSafe control levels over time. **e.** Quantile analysis of relative fitness among liver and lung metastatic colonies. Loss of *Nfib* produced significant negative fitness effects in larger liver colonies £75th percentile) at 3 weeks, whereas no significant fitness changes were observed in lung colonies. FDR < 0.05 are indicated. **f.** Density distributions of colony sizes for *sgSafe* and *sgNfib* clones in liver and lung. Vertical lines denote the modal colony size (PeakMode) for each genotype at the indicated timepoints. *Nfib* inactivation shifts the distribution toward markedly smaller colonies in the liver, whereas no detectable effect on colony-size distributions is observed in the lung. **g.** MOBA-seq identifies cutoffs for dormant colony sizes and the valley mode between dormant and expanding colonies across liver, lung, and brain. The distribution of log2-transformed tumor cell number of the pooled tumors across liver, lung, and brain at the 3-week time point was plotted, and the dormant mode and valley mode from the pooled distribution was applied across all three tissues to determine their dormant populations. This qualitative framework allows for the unbiased calculation of dormancy across tissues, time points, and sgRNAs. **h.** Gaussian mixture modeling (GMM) illustrating the fraction of dormant colonies in liver and lung for *sgSafe* and *Nfib*-targeting metastases. Unexpectedly, *Nfib* inactivation decreased the proportion of dormant colonies in the liver but not in the lung. **i.** Comparison of genotype-specific SuperMet colonies across time in the lung. Left: Scatterplots display colony size for *sgSafe* and *Nfib*-targeting metastases in the lung. SuperMet colonies that detected in the blood were labeled in red. Right: Colony size distributions showing SuperMet colony size from *sgSafe* and *sgNfib* metastases. *Nfib* inactivation has no effect in SuperMet colony size in the lung.

**Supplementary Figure 3.** MOBA-seq identified known and novel SCLC metastasis regulators. **a.** Summary plot of candidate genes included in the MOBA500 library. Circle size reflects mutation frequency in human SCLC patients, and point color denotes T-statistics derived from prior CRISPR screening in SCLC cell lines. The top 10 genes in each category are labeled. **b.** MOBA-seq identifies tumor suppressors in SCLC lung and brain metastasis. Metastatic colonies from each genotype are ranked by their mean colony size, which is indicated by a red dot. The dashed line represents the mean colony size of all control colonies combined (sgInerts). **c.** MOBA-seq identifies known and novel genetic regulators in SCLC metastasis in the lung and brain. Ranked plot of each genotype highlights top candidate suppressors (red, Log2FC>1, FDR<0.05) and drivers (blue, Log2FC<-1, FDR<0.05) of SCLC lung (left) and brain (right) metastatic burden. Grey error bars represent 95% confidence interval. sgRNAs that generate wild-type metastases are shown in green. **d.** MOBA-seq identifies metastatic fitness landscape in whole mouse. Ranked plot of each genotype highlights top candidate suppressors (red, Log2FC>1, FDR<0.05) and drivers (blue, Log2FC<-1, FDR<0.05) of whole mouse metastatic burden. Grey error bars represent 95% confidence interval. sgRNAs that generate wild-type metastases are shown in green. **e.** Comparison of mutation frequency in human SCLC tumors versus whole-mouse metastatic fitness derived from the MOBA-seq screen. Genes harboring high-frequency mutations in patients that also exhibit reduced metastatic fitness *in vivo* (red) represent putative metastasis suppressors with strong clinical relevance (e.g., *PTEN* and *CREBBP*). **f.** Comparison of T-statistics from prior CRISPR screening in SCLC cell lines with whole-mouse metastatic fitness from the MOBA-seq experiment. Genes showing strong negative or positive T-statistics and large metastatic fitness effects include known SCLC essential genes (e.g., *ASCL1*) as well as several newly identified metastasis-associated genes (red).

**Supplementary Figure 4.** Metastatic seeding is a reproducible and size-independent metric of metastasis. **a.** Metastatic seeding is independent of pre-transplantation cell abundance. Absolute pre-transplantation cell number and relative liver metastatic seeding fold change were plotted for each sgRNA. Pre-transplantation cell abundance, which was used to normalize metastatic seeding and also reflects genotype-specific proliferation during in vitro culture, showed no correlation with metastatic seeding (R² = 0.00), indicating that the enhanced metastatic phenotype cannot be explained solely by differences in cell proliferation before transplantation. **b.** Metastatic seeding is independent of peak-mode metastatic colony size. Relative liver metastatic seeding fold change was compared with relative peak-mode metastatic colony size for each sgRNA. The two measurements showed essentially no correlation (R² = 0.00), indicating that genotype-specific effects on metastatic seeding are distinct from effects on subsequent metastatic colony expansion. **c.** Metastatic seeding is independent of metastatic colony size across the colony-size distribution. For each colony-size percentile, R² values were calculated across sgRNAs between relative liver metastatic seeding and relative metastatic colony size. Correlations remained minimal across the full range of colony-size percentiles, demonstrating that metastatic seeding and metastatic outgrowth represent largely independent phenotypes. **d.** Genotype-specific metastatic seeding estimates are robust to metastatic colony-size thresholds. Heatmap showing relative liver metastatic seeding fold change for each sgRNA calculated using metastatic colony-size thresholds ranging from 1 to 1,000 cells. Genotype-specific seeding effects remained highly consistent across a broad range of colony-size cutoffs, demonstrating that metastatic seeding is a robust metric that is not dependent on the definition of a metastatic colony.

**Supplementary Figure 5.** MOBA-seq illustrates relationships between diverse metastatic metrics. **a.** Metastatic seeding is the primary factor driving overall metastatic burden in the lung. Plot of genotype-specific metastatic seeding and metastatic burden for all genotypes revealed a good linear correlation (R2=0.68). Outlier suppressors from linear regression are marked in purple. Grey error bars represent 95% confidence interval. **b.** Regression analyses evaluating the relationship between metastatic burden and multiple metastasis metrics in the lung, including 90th percentile colony size, peak mode, dormancy escape, and SuperMet percentage. Outlier from linear regression is marked in purple. **c.** Regression analyses evaluating the relationship between metastatic burden and multiple metastasis metrics in the liver, including 90th percentile colony size, peak mode, dormancy escape, and SuperMet percentage. Outlier from linear regression is marked in purple. **d.** Large metastatic colonies primarily determine outlier distance rather than metastatic burden. Metastatic colony size at different percentiles in the liver is used to model linear regression with outlier distance, metastatic burden, or metastatic seeding. The R² values for each regression are plotted against the corresponding percentile. Note that high correlations (R² > 0.7) are observed only at high percentiles (>90^th^) when modeling with outlier distance. **e.** Regression analyses evaluating the relationship between dormancy escape and multiple metastatic metrics in the liver and lung, including metastatic burden, 90th percentile colony size, and metastatic seeding. Dormancy escape shows a positive association with 90th colony size in the lung, whereas in the liver it is negatively correlated with 90th percentile colony size, highlighting tissue-specific differences in how clones exit dormancy and expand during metastatic progression.

**Supplementary Figure 6.** Metastatic suppressors and essential genes have distinct effects at different steps of the metastatic cascade. **a.** Ranked metastatic burden (log □ FC) for all genotypes in SCLC liver metastases. Inactivation of metastatic suppressors (red) increased overall metastatic burden, while inactivation of essential genes from the Rac-WRC-ARP2/3 pathway (blue) decrease overall metastatic burden. Error bars represent 95% confidence intervals. **b.** Ranked metastatic seeding (log □ FC) for all genotypes in SCLC liver metastases. Inactivation of most metastatic suppressors (red) increased metastatic seeding, with notable exceptions on *Tsc1* and *Tsc2*. In contrast, loss of essential genes within the Rac–WRC–ARP2/3 pathway (blue) consistently decreased metastatic seeding. Error bars indicate 95% confidence intervals. **c.** Ranked 90th percentile colony size (log □ FC) for all genotypes in SCLC liver metastases. Inactivation of most metastatic suppressors (red) increased colony size, with notable exceptions such as *Gata6* and *Csnk1a1*, which did not enhance clonal expansion. In contrast, loss of essential genes in the Rac– WRC–ARP2/3 pathway (blue) had minimal effects on 90th percentile colony size, with the exception of *Rac1*, which significantly reduced large-clone outgrowth. Error bars represent 95% confidence intervals. **d.** Ranked peak mode colony size (log □ FC) in SCLC liver metastases. Metastatic suppressors exhibit diverse effects on the exponential clonal expansion mode, reflecting heterogeneous control of metastatic growth. Loss of essential genes in the Rac–WRC–ARP2/3 pathway did not yield sufficient colonies to reliably calculate peak-mode colony size. **e.** Ranked dormancy escape percentage (log □ FC) in SCLC liver metastases. Metastatic suppressors display diverse effects on the ability of dormancy escape. Notably, inactivation of strong suppressors such as *Tsc2* and *Crebb***p** markedly reduced dormancy escape in the liver. Loss of essential genes in the Rac–WRC–ARP2/3 pathway did not yield sufficient colonies to reliably calculate dormancy escape. **f.** Ranked SuperMet percentage (log □ FC) in SCLC liver metastases. Inactivation of most metastatic suppressors (red) increased SuperMet percentage. In contrast, loss of essential genes within the Rac– WRC–ARP2/3 pathway (blue) consistently decreased SuperMet percentage.

**Supplementary Figure 7.** MOBA-seq identifies genotype-specific metastasis effects in immune deficient mice. **a.** Genotype-ranked metastatic metrics in liver metastases from NSG mice, including metastatic burden, metastatic seeding, 90th percentile colony size, SuperMet percentage, peak mode size, and dormancy escape (all as in log □ FC relative to sgSafe). **b.** Genotype-ranked metastatic metrics in lung metastases from NSG mice, including metastatic burden, metastatic seeding, 90th percentile colony size, SuperMet percentage, peak mode size, and dormancy escape (all as in log □ FC relative to sgSafe). **c.** Genotype-ranked metastatic metrics in brain metastases from NSG mice, including metastatic burden, metastatic seeding, 90th percentile colony size, and SuperMet percentage (all as in log □ FC relative to sgSafe). Brain metastasis did not yield sufficient colonies to reliably calculate peak mode size, and dormancy escape. **d.** Targeting the Rac1-WRC-ARP2/3 pathway consistently affects metastatic burden and seeding but has little effect on colony size in both C57BL/6 and NSG mice. The error bars represent 95% confidence interval. **e.** MOBA-seq identified diverse patterns for metastasis suppression in both C57BL/6 and NSG mice. Radar plots stratify genetic suppressors by their effects on six metastatic metrics throughout the metastatic cascade.

**Supplementary Figure 8.** G*p*atch8 suppresses brain metastatic outgrowth in an immune-dependent manner. **a.** Inactivation of *Gpatch8* in RP48 cells increased brain metastatic burden (*left*), metastatic seeding (*middle*), and 90th-percentile metastatic colony size (*right*) in immunocompetent C57BL/6 mice relative to safe-targeting controls, supporting *Gpatch8* as a brain metastasis suppressor. Data points outlined in black indicate statistically significant comparisons (FDR < 0.05). **b.** Inactivation of *Gpatch8* in RP48 cells increased brain metastatic seeding (*middle*) but reduced overall brain metastatic burden (*left*) and 90th-percentile colony size (right) relative to safe-targeting controls in immunodeficient NSG mice. These divergent effects suggest that *Gpatch8* loss promotes initial brain seeding but does not enhance subsequent metastatic outgrowth in the absence of an intact immune system, supporting an immune-dependent role for Gpatch8 in suppressing brain metastatic expansion. Data points outlined in black indicate statistically significant comparisons (FDR < 0.05).

**Supplementary Figure 9.** Genotype-specific regulation of cross-tissue SuperMet dissemination. **a.** Heatmaps showing cross-tissue dissemination of shared metastatic colonies in immunodeficient NSG mice. For each sgRNA genotype, the percentage of colonies originating in each tissue that were also detected in each destination tissue was calculated. Colors indicate the log2 fold change relative to control guides. Inactivation of *Crebbp*, *Tsc1*, and *Tsc2* increased cross-tissue dissemination of SuperMets, whereas loss of *Kctd10* and *Zeb1* reduced dissemination across tissues. **b.** Heatmaps showing cross-tissue dissemination of shared metastatic colonies in immunocompetent C57BL/6 mice. For each sgRNA genotype, the percentage of colonies originating in each tissue that were also detected in each destination tissue was calculated. Colors indicate the log2 fold change relative to control guides. Inactivation of *Crebbp*, *Gpatch8*, and *Pten* increased dissemination of SuperMets between specific tissue pairs.

**Supplementary Figure 10.** Recipient sex modulates immune surveillance of SCLC metastasis. **a.** Schematic of the MOBA-seq experimental design used to evaluate sex-specific metastatic phenotypes. Cas9D SCLC cells (RP48 and RP116) transduced with barcoded lenti-sgRNAs were injected intravenously into male or female immune-deficient NSG or immune-competent C57BL/6 mice (N = 5 per sex per mouse genotype). Following metastatic colonization in liver, lung, and brain, barcode-resolved clonal analysis was performed to quantify genotype-specific metastasis behavior. **b.** RP48 and RP116 cells exhibited distinct metastatic growth phenotypes, whereas female recipient mice showed stronger suppression of metastatic seeding. Syngeneic SCLC cells of male (RP48) and female (RP116) origin were transplanted into both male and female recipient mice with immune proficient (C57BL/6) and deficient (NSG) backgrounds. **Left**: Barcoded liver metastatic colonies from five mice are plotted, with each point representing one colony. Color indicates the sex of mice, and shape indicates the sex of cancer cells. Note that male cells grow better, and female mice have a stronger immune response. **Right**: Metastatic colony size distribution of each mouse-cell sex combinations. Note that female but not male SCLC cells are more sensitive to immune suppressions evidenced by a decreased clonal expansion peak mode. **c.** Heatmaps of genotype-specific metastatic burden (log □ FC) in the liver. Each row corresponds to an sgRNA-defined genotype, and each column represents an individual mouse annotated by sex and immune background. Strong sex-dependent differences in metastatic suppression emerge, particularly in female C57BL/6 mice. Color bars indicate gene categories. Top panel: male-derived RP48 cells; bottom panel: female-derived RP116 cells. **d.** Sex– and immune-stratified significance matrices for genotype-specific metastatic burden, metastatic seeding, dormancy escape, and percentile colony sizes (90th, 50th, and 20th) in liver and lung. Each square represents the measured difference in a genotype-specific metastatic metric for a given sex– or immune-state comparison, with significant associations (*FDR* < 0.05) indicated by a star within the box. Top panel: liver metastasis; bottom panel: lung metastasis.

**Supplementary Figure 11.** C***r***ebbp is a potent metastasis suppressor. **a.** *Crebbp* inactivation increases SCLC metastatic burden at three weeks. Bar plot showing the relative proportions of metastatic tumor cells by their genotype. Shown is the representation of SCLC genotypes recovered from liver, lung, brain, and bone marrow (BM) tissues of both NSG and C57BL/6 mice after three weeks of growth, compared with the pre-transplantation sample. Three *sgCrebbp* genotypes are highlighted. **b.** *Crebbp* inactivation increases SCLC metastatic seeding across multiple tissues. Metastatic colonies recovered from liver, lung, brain, and bone marrow at three weeks are shown, with each dot representing an individual barcode-defined colony and dot size proportional to colony cell number. Total colony counts and mean colony sizes are indicated for *sgNT3* and *Crebbp_sg2*, which exhibited comparable cell numbers in the pre-transplantation pool. Across all organs and timepoints, *Crebbp* inactivation consistently increased the number and size of metastatic colonies relative to controls. **c-h.** *Crebbp* inactivation promotes liver metastasis in both RP116 and RP48 models across immune backgrounds. In RP116 cells (**c-e**), *Crebbp* inactivation increased relative metastatic burden (**c**), metastatic seeding (**d**), and 90th-percentile colony size (**e**) in the liver of both NSG and C57BL/6 mice compared with control sgRNAs. In RP48 cells (**f-h**), Crebbp inactivation increased relative metastatic burden (**f**) and metastatic seeding (**g**) in the liver in both NSG and C57BL/6 mice, whereas it increased 90th-percentile colony size (**h**) in NSG mice but not in C57BL/6 mice. For each time point, values are plotted as fold change relative to control sgRNAs. Data points outlined in black indicate statistically significant comparisons (FDR < 0.05).

**Supplementary Figure 12.** Crebbp is a shared metastasis seeding suppressor across multiple tissues. **a-d**. Comparison of genotype-specific RP48 cell metastatic seeding fold changes between C57BL/6 and NSG mice in the liver (**a**), lung (**b**), brain (**c**), and bone marrow (**d**). Dashed lines indicate the wild-type reference level. Genotypes that significantly increased metastatic seeding in both immune backgrounds are highlighted as shared metastasis suppressors, whereas genotypes that significantly decreased metastatic seeding in both backgrounds are highlighted as shared metastatic drivers. *Crebbp* loss consistently increased metastatic seeding across multiple tissues, including liver, lung, brain, and bone marrow, identifying *Crebbp* as a broad metastasis suppressor whose effect is conserved across distinct metastatic tissue and immune contexts.

**Supplementary Figure 13.** cNMF analysis reveals that Crebbp loss shifts SCLC cell-state programs toward an SCLC-I-like state. **a.** Selection of the optimal number of cNMF components based on solution stability and reconstruction error across the indicated numbers of components. Four components were selected for downstream analysis. **b.** Gene expression programs (GEPs) identified by cNMF. Dot size and color indicate the relative contribution of individual genes to each program, highlighting distinct transcriptional features associated with GEP 1 to GEP 4. **c.** Association of each cNMF program with canonical SCLC subtype signatures. Heatmap shows enrichment of representative SCLC-A, SCLC-N, SCLC-P and SCLC-I marker genes across GEP 1 to GEP 4. Note that GEP 3 showed the strongest enrichment for the SCLC-I-associated program, including Vim, Axl and Zeb2, whereas GEP 1 was enriched for SCLC-A features. **d.** Comparison of cNMF program usage between Crebbp-KO and Crebbp-WT cells. Crebbp loss significantly altered usage of all four transcriptional programs, including increased usage of the SCLC-I-associated GEP 3 and decreased usage of GEP 2, consistent with a shift toward an SCLC-I-like transcriptional state. Box plots show the distribution of program usage across individual cells.

**Supplementary Figure 14.** Loss of *Crebbp* alters SCLC cell state through regulation of *Cdx2*. **a.** Gene set enrichment analysis (GSEA) of snRNA-seq from *Crebbp*-WT and *Crebbp*-KO RP48 SCLC liver metastases. Heatmaps show normalized enrichment scores (KO vs. WT) across major cancer– and stroma-associated transcriptional programs. *Crebbp* loss leads to upregulation of MYC– and cell-cycle–associated signaling pathways. **b.** Transcriptome-wide correlation between differential gene expression measured by bulk RNA-seq and snRNA-seq. A moderate Pearson correlation (r = 0.51) indicates substantial concordance between mRNA– and pre-mRNA based expression changes across platforms. **c.** Heatmap of bulk RNA-seq profiles for the top 500 differentially expressed genes from sorted *Crebbp*-WT and *Crebbp*-KO RP48 SCLC liver metastases. Clustering reveals the emergence of distinct transcriptional status upon *Crebbp* loss, including marked downregulation of *Cdx2* in *Crebbp*-KO cells. **d.** Pie charts summarizing differential H3K27ac ChIP-seq peaks identified in *Crebbp*-WT and *Crebbp*-KO cells. **e.** Doxycycline-induced CDX2 re-expression in *Crebbp*-KO RP48 cells suppresses proliferation and growth advantage in co-culture with *Crebbp*-WT cells. *Crebbp*-WT-TetON-CDX2 and *Crebbp*-KO-TetON-CDX2 RP48 cells were mixed at a 1:1 ratio and treated with 100 ng/mL doxycycline in co-culture for 0, 4, and 8 days. GFP/mCherry cell ratios were quantified by flow cytometry. n= 6 wells **f.** Doxycycline-induced CDX2 re-expression in *Crebbp*-KO RP48 cells suppresses transwell migration in co-culture with *Crebbp*-WT cells. *Crebbp*-WT-TetON-CDX2 and *Crebbp*-KO-TetON-CDX2 RP48 cells were mixed at a 1:1 ratio and treated with 100 ng/mL doxycycline for 4 days for transwell migration. GFP/mCherry cell ratios were quantified by flow cytometry. n= 6 wells **g.** *Crebbp*-mediated metastatic seeding suppression is CDX2-dependent. MOBA-seq analysis of genotype-specific metastatic seeding in control and CDX2-overexpressing RP48 cells that metastasized to the lung and liver. *Crebbp* inactivation increased metastatic seeding in control RP48 cells but had no effect in CDX2-overexpressing RP48 cells. **h.** *Crebbp*-mediated metastatic clonal expansion is CDX2-independent. MOBA-seq analysis of genotype-specific clonal expansion (90^th^ percentile colony size) in control and CDX2-overexpressing RP48 cells that metastasized to the lung and liver. *Crebbp* inactivation increased metastatic clonal expansion in both control and CDX2-overexpressing RP48 cells.

**Supplementary Figure 15.** Loss of *Crebbp* alters liver metastatic microenvironment. **a.** Quality metrics of Xenium5k spatial transcriptomics data from *Crebbp*-WT and *Crebbp*-KO liver and lung metastases. Histograms show gene counts per cell. **b.** Spatial maps of ASCL1 □ cells in *Crebbp*-WT and *Crebbp* –KO liver and lung metastases. *Crebbp* –KO lesions show a marked increase in ASCL1 □ cell abundance in both liver and lung. **c.** UMAP visualization of spatially profiled cells from WT and KO liver (top) and lung (bottom) metastases. Color-coded clusters reveal extensive transcriptional reorganization in *Crebbp*-KO liver tumors, including pronounced remodeling of immune and stromal niches. **d.** Comparison of vascularized metastatic colony numbers between WT and KO liver metastases based on H&E staining. *Crebbp* loss significantly increases the number of vascularized metastatic colonies (Wilcoxon p = 0.045). **e.** Spatial mapping of glucose catabolism signatures across liver metastases. KO tumors show strong regional upregulation of glucose metabolic activity compared with WT. Right: Violin plots quantify signature scores for WT versus KO conditions. **f.** Spatial mapping of lipid metabolism signatures across liver metastases. KO tumors demonstrate increased lipid metabolic activity, suggesting metabolic rewiring following Crebbp inactivation. Right: Corresponding violin plots compare WT versus KO conditions. **g.** Representative image of *Crebbp*-KO liver metastases. Regions enriched for cancer-stroma interactions are highlighted. **h.** Representative spatial transcriptomic images showing (left) immune-cell recruitment pathways localized at the cancer–endothelial–immune interface and (right) T-cell exhaustion markers concentrated at the cancer–immune interface. **i.** Representative immunofluorescence of WT and KO liver metastases. KO tumors show increased infiltration of T cells (CD3), NK cells (NK1.1), and macrophages (F4/80). **j.** Quantification of CD3 □ T cells, F4/80 □ macrophages, and NK1.1 □ natural killer cells in WT versus KO liver metastases. KO tumors exhibit significantly elevated infiltration across multiple immune subsets. **k.** Flow cytometry quantification of major immune subsets in WT versus KO liver metastases. *Crebbp* loss increases T cells, NK cells, but not B cells. **l.** Analysis of human tumor mutation burden (TMB) from CREBBP-mutant versus CREBBP-wild-type cancers in the MSK-IMPACT cohort. CREBBP-mutant lesions display significantly increased TMB (Wilcoxon p < 2.2 x 10D’D).

**Supplementary Figure 16.** C*R*EBBP mutation is associated with improved survival specifically in patients treated with immune checkpoint inhibitors. Multivariable Cox proportional hazards analysis of overall survival in the pan-cancer MSK-IMPACT cohort restricted to ICI-treated patients, including CREBBP mutation status, age, sex and metastatic status as covariates. *CREBBP*-mutant tumors were associated with significantly improved survival compared with CREBBP-wild-type tumors, whereas age, sex and metastatic status were not significantly associated with survival. These data identify *CREBBP* mutation as an independent correlate of favorable outcome among ICI-treated patients.

## METHODS

### Mouse research

The use of mice for the current study has been approved by the Institutional Animal Care and Use Committee at Washington University, protocol number 23-0283. Sex-balanced cohorts of mice aged 8-15 weeks were used at the time of tumor initiation. Mice were housed under a 12-h light/12-h dark cycle at 68–73 °F and 40–60% relative humidity.

### Cells and Reagents

NCI-H82 (HTB-175) and HEK293T (CRL-3216) were originally purchased from ATCC; RP48 and RP116 small cell lung cancer cells were generated in the Sutherland Lab; H82 cells were cultured in RPMI1640 medium containing 10% FBS, 100 units/mL penicillin and 100 μg/mL streptomycin. RP48 and RP116 cells were cultured in DMEM/F12 (Thermo Fisher), 10% (vol/vol) FBS, 6 ng/mL Mouse EGF (FUJIFILM IRVINE SCIENTIFIC INC), and 4 µg/mL Hydrocortisone (Cayman Chemical). All cell lines were confirmed to be mycoplasma negative (MycoAlert Detection Kit, Lonza). All plasmids used in this study were listed in Supplementary Table 1 and are available from our laboratory upon request.

### Design, generation, barcoding, and production of lentiviral vectors

sgRNA sequences targeting the putative tumor suppressor genes were designed using CRISPick (https://portals.broadinstitute.org/gppx/crispick/public). All sgRNA sequences are shown in Supplementary Table 1. The generation of the barcode fragment containing the degenerate random barcode, and subsequent cloning of sgRNA oligo pool into the vectors, were performed as previously described in supplementary figure 1.

Lentiviral vectors were produced using polyethylenimine (PEI)-based transfection of 293T cells with delta8.2 and VSV-G packaging plasmids in 150 mm cell culture plates. Sodium butyrate (Sigma Aldrich, B5887) was added 8 hours after transfection to achieve a final concentration of 20 mM. Media was refreshed 24 hours after transfection. 20 mL of virus-containing supernatant was collected 36, 48, and 60 hours after transfection.

### SCLC transplantation via tail vein injection

For intravenous transplantations, 2.5 x 10^6^ SCLC cells were injected into one of the lateral tail veins. Mice were sacrificed 21 days post-injection and lung, liver, brain, bone marrow, and blood cells were collected for gDNA extraction. For subcutaneous transplantations, 3× 10^6^ of SCLC cells were re-suspended in 200 µL Matrigel^®^ Basement Membrane Matrix (Corning, 354234) and injected into two parallel sites per mouse. Mice were sacrificed 28 days post-injection. Both primary SubQ tumors and metastases-bearing liver were dissected and the weight, height, width, and length, of each tumor was measured.

Maximal tumor size/burden permitted by Institutional Animal Care and Use Committee (IACUC) is 1.75 cm^3^, the maximal tumor size/burden was not exceeded in our study. IACUC approved all animal studies and procedures.

### MOBA-seq library generation

Genomic DNA was isolated from bulk tumor-bearing tissues from each mouse as previously described^21^. Briefly, benchmark monoclonal control cell lines were generated from B16F10 cells transduced by a barcoded Lenti-sgDummy/PuroR vector (sgDummy: a control sgRNA with dual Bsmb1 cloning sites, AGAGACGCTCGAGCGTCTCT) and selected for monoclonal culture. Eight to ten benchmark control cell lines with varying number of cells were added to each mouse tissue sample prior to lysis to enable the calculation of the absolute number of cancer cells in each tumor from the number of sgID-BC reads. Following homogenization and overnight protease K digestion, genomic DNA was extracted from the tissue lysates using standard phenol-chloroform and ethanol precipitation methods. Subsequently, Q5 High-Fidelity 2x Master Mix (New England Biolabs, M0494X) was used to amplify the sgID-BC region from 40 μg of genomic DNA in a total reaction volume of 400 μl per sample for 1^st^ round PCR amplification. The primers used were Forward: ATA ATG TGT GTG GTA CAA AAG GTC and Reverse: GAA TTC CAT GTT AAT TAA GCC ATA GGC. The PCR products were purified with the QIAquick PCR Purification Kit (Qiagen, 28104). Q5 High-Fidelity 2x Master Mix (New England Biolabs, M0494X) was used to amplify the sgID-BC region from 10% of the purified PCR product in a total reaction volume of 200 μl per sample for the NGS PCR amplification. The unique dual-indexed primers used were Forward: AAT GAT ACG GCG ACC ACC GAG ATC TAC AC-8 nucleotides for i5 index-ACA CTC TTT CCC TAC ACG ACG CTC TTC CGA TCT-4 to 7 random nucleotides for increased diversity-AGG GTT ACA GTT TAG TCA CCA TA and Reverse: CAA GCA GAA GAC GGC ATA CGA GAT-6 nucleotides for i7 index-GTG ACT GGA GTT CAG ACG TGT GCT CTT CCG ATC T-9 to 6 random nucleotides for increased diversity-CGA CTC GGT GCC ACT TTT TC. The PCR products were purified with the QIAquick PCR Purification Kit (Qiagen, 28104). The concentration and quality of the purified libraries were determined using the Invitrogen 1X dsDNA High Sensitivity kit (Invitrogen, Q33231) on the Invitrogen Qubit 4 Fluorometer (Invitrogen, Q33238). The libraries were assigned one to ten million reads based on tissue type to ensure adequate sequencing coverage, purified with the QIAquick Gel Extraction kit (Qiagen, 28704), and sequenced (read length 2×150bp) on the NovaSeq X Plus platform (GTAC@MGI, Novogene).

### Generation of Stable Cell Lines

Parental cells were seeded at 50% confluency in a 6-well plate the day before transduction (day 0). The cell culture medium was replaced with 2 mL fresh medium containing 8 µg/mL hexadimethrine bromide (Sigma Aldrich, H9268-5G), 20 µL ViralPlus Transduction Enhancer (Applied Biological Materials Inc., G698) and 40 µL concentrated lentivirus, then cultured overnight (Day 1). The medium was then replaced with complete medium and cultured for another 24 hours (Day 2). Cells were transferred into a 100 mm cell culture dish with appropriate amounts of antibiotic (Blasticidin doses: RP48: 50 µg/mL; RP116: 20 µg/mL; H82: 25 µg/mL; Puromycin doses: RP48: 3 µg/mL; RP116: 3 µg/mL; H82: 1 µg/mL) and selected for 48 hours (Day 3).

### Western Blot

Cells were lysed in RIPA buffer (50 mM Tris-HCl (pH 7.4), 150 mM NaCl, 1% Nonidet P-40, and 0.1% SDS) and incubated at 4 °C for 15 minutes, followed by centrifugation at 13,000 × rcf for 10 minutes at 4 °C. The supernatant was collected, and the protein concentration was determined by BCA assay (Thermo Fisher Scientific, 23250). Histones were acid extracted using Histone Extraction Kit (Active Motif, 40028), neutralized, and concentration measured according to protocol. Protein extracts (20 μg) and histone extracts (2^g) were separated on 4-12% SDS-PAGE and transferred onto PVDF membranes activated by methanol for 60s. The membranes were blocked with 5% non-fat milk in TBS with 0.1% Tween 20 (TBST) at room temperature for one hour, cut according to the molecular weight of the target protein (with at least two flanking protein markers), followed by incubation with primary antibodies diluted in TBST at 4 °C overnight. After three 10-minutes washes with TBST, the membranes were incubated with the appropriate secondary antibody conjugated to HRP diluted in TBST (1:10000) at room temperature for 1 hour. After three 10-minutes washes with TBST, protein expression was quantified with enhanced chemiluminescence reagents (Fisher Scientific, PI80196).

Antibodies used in this study: GAPDH (Cell Signaling, 5174S, 1:2000), CREBBP (Cell Signaling, 7389S, 1:200), H3 (Cell Signaling, 4499S, 1:1000), CDX2 (Invitrogen, MA5-35215, 1:2000), H3K27ac (Cell Signaling, 8173S, 1:1000), Goat-anti-Rabbit IgG Antibody, HRP-conjugate (Sigma-Aldrich, 12-348), Goat-anti-Mouse IgG Antibody, HRP-conjugate (Thermo Fisher Scientific, 62-6520).

### H3K27ac ChIP-seq

H3K27ac ChIP-seq was performed using the SimpleChIP Plus Enzymatic Chromatin IP Kit (Magnetic Beads; Cell Signaling Technology, #9005) according to the manufacturer’s protocol. Crebbp-WT and Crebbp-KO RP48 cells were crosslinked with 1% formaldehyde, and chromatin was fragmented by micrococcal nuclease digestion. Chromatin was immunoprecipitated overnight with H3K27ac antibody (Cell Signaling Technology, #8173), captured with Protein G magnetic beads, and purified after crosslink reversal. ChIP and matched input DNA were used for sequencing library preparation. Two biological replicates were analyzed per genotype.

### Flow Cytometry

Metastases bearing mouse tissues were dissected, cut into small pieces and digested in 5 mL tissue digest media (3.5 mL HBSS-Ca2+ free, 0.5 mL Trypsin-EDTA [0.25%], 5 mg Collagenase IV [Worthington], 25 U Dispase [Corning]) for 30 min in hybridization chamber at 37°C with rotation. Digestion is then neutralized by adding 5 mL ice cold Quench Solution (4.5 mL L15 media, 0.5 mL FBS, 94 µg DNase). Single-cell suspensions were generated by filtering through a 40 µm cell strainer, spinning down at 500 rcf for 5 min and washed with PBS twice.

Digested single-cell suspensions were subjected to a 30% Percoll gradient for separation of cancer and immune cells from hepatocytes and fat. In short, a stock isotonic Percoll (SIP) solution was made combining one-part 10X HBSS (14065056, ThermoFisher Scientific) with nine-parts of Percoll (P1644, Millipore Sigma). Digested liver samples were centrifuged at 300g x 8 minutes to pellet. Upon removal of the supernatant, pellets were resuspended in 3mL of SIP and 7mL of 1X HBSS to create a 30% Percoll solution. Samples were centrifuged at room temperature for 30 minutes at 1000g; acceleration:9, brake:1. After centrifugation, the top layer of the Percoll gradient and interphase was removed, leaving behind a heterogeneous cell pellet that was washed with 1X HBSS to remove leftover Percoll. Post wash, cells were counted and plated in a 96-well U-bottom plate at a concentration of 1×10^6^ cells of one sample, per well, for staining.

Once plated, samples were centrifuged (500g x 5 min.) to pellet cells, supernatant was removed, and samples were resuspended in 100uL of Cell Staining Buffer (420201, BioLegend) containing LIVE DEAD Blue staining (1:400 dilution) and CD16/CD32 monoclonal antibody FC Block (1:200 dilution) and incubated at room temperature (RT) in the dark for 10 minutes. Samples were then washed and resuspended with the surface antibody cocktail (see antibody table below) prepared in a 1:1 mixture of Cell Staining Buffer and BD Horizon Brilliant Stain buffer (566349, BD Biosciences) for 25 minutes at room temperature in the dark. After staining, samples were washed once with Cell Staining Buffer and resuspended in 100uL buffer prior to acquiring data on a Cytek Aurora Spectral Flow Cytometer. Analysis of acquired data was performed using FlowJo_v.10.10.0.

| Marker | Fluorophore | Dilution | Catalog number & Manufacturer |
| --- | --- | --- | --- |
| Viability | LIVE DEAD Blue | 1:400 | L23105, ThermoFisher Scientific |
| CD45 | BUV737 | 1:300 | 752414, BD Biosciences |
| CD11b | BV605 | 1:300 | 101257, BioLegend |
| LY-6C | RB613 | 1:300 | 759408, BD Biosciences |
| Ly-6G | RB780 | 1:200 | 569153, BD Biosciences |
| F4/80 | RY703 | 1:300 | 770168, BD Biosciences |
| CD80 | BV650 | 1:300 | 104731, BioLegend |
| CD206 | BV711 | 1:300 | 141727, BioLegend |
| CD161 (NK1.1) | BUV563 | 1:300 | 741223, BD Biosciences |
| CD19 | RB705 | 1:300 | 570651, BD Biosciences |
| CD3 | BUV395 | 1:200 | 740268, BD Biosciences |
| CD4 | APC-Fire 810 | 1:200 | 100480, BioLegend |
| CD8 | BUV805 | 1:200 | 368-0081-82, ThermoFisher Scientific |
| CD44 | PerCP | 1:300 | 103036, BioLegend |
| CD25 | PE-CF594 | 1:200 | 562694, BD Biosciences |
| CD62L | BV510 | 1:200 | 563117, BD Biosciences |
| CD11c | Spark Blue 550 | 1:200 | 117365, BioLegend |
| MHC Class II (I-Ab) | BV480 | 1:300 | 566086, BD Biosciences |
| CD279 (PD-1) | PE-Fire 744 | 1:200 | 135269, BioLegend |
| CD274 (PD-L1) | PE-Fire 640 | 1:200 | 124345, BioLegend |
| FC BLOCK | CD16/CD32 | 1:200 | 14-0161-86, ThermoFisher Scientific |

### Analysis of MSK-IMPACT data

Clinical and genomic data were obtained from the cBioPortal for Cancer Genomics using the cBioPortalData R package. Two cohorts were analyzed: (1) the MSK-IMPACT 2017 cohort (study ID: msk_impact_2017), comprising comprehensive genomic profiling data from 10,336 patients across multiple cancer types, and (2) the MSK immunotherapy cohort (study ID: tmb_mskcc_2018), consisting of the 1,610 patients from the MSK-IMPACT assay treated with immune checkpoint inhibitors (ICI). Clinical data, including overall survival (OS) status and duration in months, were extracted for each cohort. *CREBBP* mutation status was assigned based on the presence of non-silent mutations on chromosome 16 spanning the CREBBP locus (chr16:3,775,055-3,930,727, GRCh37).

For comparative analysis of ICI treatment effects, patients were stratified into two treatment groups. The ICI-treated cohort consisted of all patients in the MSK immunotherapy dataset with matched clinical and mutation sequencing data (n=1,610), which included both patients from the MSK-IMPACT 2017 cohort and additional patients profiled from MSK-IMPACT. The non-ICI cohort comprised patients from the MSK-IMPACT 2017 dataset excluding those present in the immunotherapy cohort (n=6,705), with patient overlap verified by unique patient identifiers. 16 patients who had multiple samples with differing CREBBP mutation status were excluded to ensure homogeneous classification, and patients without survival data were also excluded, leaving a final patient number n=6,705 for the non-ICI cohort. Overall survival was analyzed using Kaplan-Meier curves with the survfit function from the survival R package with each patient providing one sample, and survival differences between groups were assessed using log-rank tests. Pairwise comparisons were performed to evaluate the effects of ICI treatment and CREBBP mutations on overall survival. All statistical analyses were performed in R version 4.4.1, and visualizations were generated using the survminer and ggplot2 packages.

### Processing of paired end reads to identify the sgID and barcode

Sequencing of MOBA-seq libraries produces reads that are expected to contain a 23-nucleotide barcode (BC) of the form GTTNNNNNNNNNNNNNNNNNATG, where each of the 17 Ns represents a random nucleotide, followed by a 17 to 21 nucleotide sgID. Each sgID has a one-to-one correspondence with an sgRNA in the lentiviral-sgRNA library (sgID dictionary). Note that all sgID sequences in the viral pool differ from each other by at least five nucleotides such that incorrect sgID assignment due to PCR or sequencing error is extremely unlikely. The random 17-nucleotide portion of the BC is expected to be unique to each lentiviral integration event and thus tags all cells in a single clonal expansion. Note that the length of the barcode ensures a high theoretical potential diversity (∼4^17^ > 10^10^ barcodes per vector), so while the actual diversity of each Lenti-sgRNA/PuroR vector is dictated by the number of colonies generated during the plasmid barcoding step, it is very unlikely that we will observe the same BC in multiple clonal expansions.

FASTQ files were parsed using regular expressions to identify the sgID and BC for each read. To minimize the effects of sequencing error on BC identification, we required the forward and reverse reads to agree completely within the 23-nucleotide sequence to be further processed. Due to PCR and sequencing errors, genuine tumor barcodes can occasionally appear mutated. These “fake” tumor colonies are expected to be rare and sporadic. To minimize their impact, each barcode was converted into a 2D array of characters (n_barcodes x n_barcodes x barcode_length), and NumPy broadcasting was used to compute all pairwise Hamming distances simultaneously, while masking redundant comparisons. For barcode pairs with distances <=2, we retained the barcode with the higher read count.

After filtering, absolute cancer cell numbers for each barcode were calculated based on the regression of spike-in read counts against their known expected cell numbers. If not all expected spike-ins were detected in the sequencing data, missing spike-ins were assigned using the average ratio of observed counts to expected cell numbers across detected spike-ins. For samples with multiple spike-ins detected, linear regression without intercept was performed using the spike-in read counts and their expected cell numbers to calculate a conversion factor based on the slope of the regression line. Spike-ins with read counts below 10, relative counts outside ±50% of expected values, or high residuals (when R² < 0.9) were excluded to improve regression quality. This conversion factor was applied to all barcodes to estimate cell numbers, and barcodes with estimated cell numbers ≤1 were excluded from downstream analysis. For pre-transplantation samples lacking spike-ins, a default conversion factor of 10 cells per read was applied. The median sequencing depth across experiments was ∼1.3 reads per cell.

### Summary statistics for metastatic related statistics

To assess the extent to which a given gene (*X*) *a*ffects metastatic clonal expansion, we compared the distribution of tumor sizes produced by vectors targeting that gene (sg*X* tumors) to the distribution produced by our negative control vectors (sg*Inert* tumors). We relied on two statistics to characterize these distributions: the size of tumors at defined percentiles of the distribution (specifically the 50^th^, 60^th^, 70^th^, 80^th^, 90^th^, and 95^th^ percentile tumor sizes), and the peak mode. The percentile sizes are nonparametric summary statistics of the tumor size distribution. The peak mode identifies the most common tumor size by determining the maximum of the kernel density estimate of log10-transformed cell numbers, computed using 512 evaluation points across the data range. This metric allows for quantification of the expansion of tumor colonies.

To quantify the extent to which each gene suppressed or promoted tumor growth, we normalized statistics calculated on tumors of each genotype to the corresponding inert statistic. The resulting ratios reflect the growth advantage (or disadvantage) associated with each tumor genotype relative to the growth of *sglnert* tumors.

For example, the relative i^th^ percentile size for tumors of genotype X was calculated as:

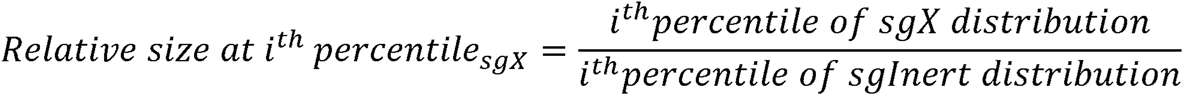

Likewise, the relative peak mode size for tumors of genotype X was calculated as:

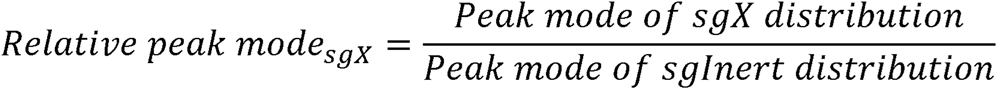

In addition to the tumor size metrics described above, we characterized the effects of gene inactivation on tumorigenesis in terms of the number of seeded tumors (“metastatic seeding”) and total neoplastic cell number (“metastatic burden”) associated with each genotype. Unlike the aforementioned metrics of tumor size, metastatic seeding and burden are linearly affected by pre-transplantation cell number and are thus sensitive to underlying differences in the representation each barcoded tumor genotype in the pre-transplantation cells. Therefore, to assess the extent to which a given gene (*X)* affects metastatic seeding, we therefore first normalized the number of sg*X* tumors to the number of pre-transplantations sg*X* cells to account for differences in initial tumor transplantation:

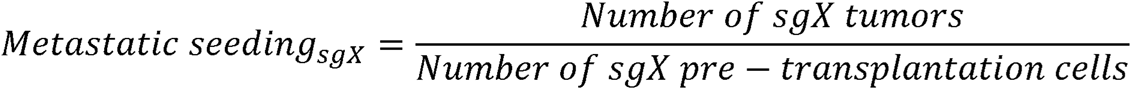

As with the tumor size metrics, we then calculated a relative metastatic seeding by normalizing this statistic to the corresponding statistic calculated using *sglnert* tumors:

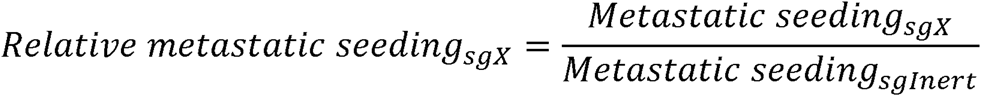

Genes that influence relative metastatic seeding alter the probability that disseminated cancer cells establish detectable metastatic colonies and/or survive during the early post-seeding phase. Relative metastatic seeding thus captures an additional and potentially important aspect of tumor suppressor gene function.

Analogous to the calculation of relative metastatic seeding, we characterized the effect of each gene on metastatic burden by first normalizing the sg*X* tumor burden to tumors to the number of pre-transplantations sg*X* cells:

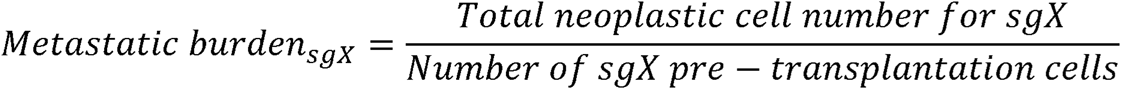

We then calculated a relative metastatic burden by normalizing this number to the corresponding statistic calculated using sgInert tumors:

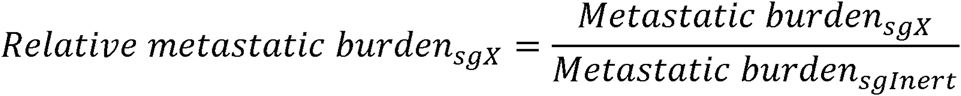

Metastatic burden is an integration over tumor size and number and thus reflects the total neoplastic load in each mouse. Metastatic burden is thus more strongly related to morbidity than are our metrics of tumor size and is closely related to traditional measurements of tumor progression such as duration of survival and tumor area. While intuitively appealing, tumor burden is notably noisier than our metrics of tumor size as it is strongly determined by the size of the largest tumors. To study the potential for aggressive metastatic colonies to release circulating tumor cells into the blood, we defined super-metastasis (“SuperMet”) as the percentage of disseminating tumors, defined as those tumor barcodes that were detected in both the target tissue and blood at matched time points within the same mouse, of all colonies:

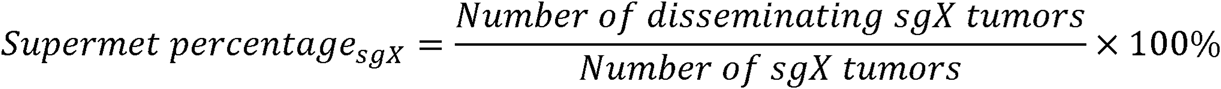

In addition, we also calculated the peak mode of disseminating tumors to characterize the size at which tumors release cancer cells. As with the tumor size metrics, we normalized the supermet percentage calculated on tumors of each genotype to the corresponding inert statistic to reflect the impact on dissemination potential of each genotype relative to that of sg*Inert* tumors:

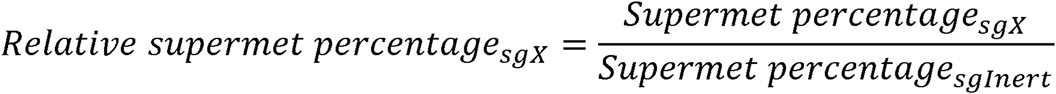

Finally, we also characterized the effects of gene inactivation on tumor arrest in the dormant phase by calculating the percentage of tumors that were found to have remained at a size considered to be dormant. The size cutoff for tumors to be considered dormant was calculated through two methods. We applied Gaussian mixture modeling to the distribution of log2-transformed tumor cell numbers within each tissue and time point. We fit two-component GMM models with unequal variances (modelNames=“V”) using the mclust package, allowing us to distinguish between dormant and expanding tumor populations. For early and intermediate time points, we employed a constrained fitting approach where the mean of the dormant component was fixed to a predetermined value based on biological expectations or data from later time points. This constraint improves model stability and ensures consistent identification of the dormant population across time. The constrained model uses an expectation-maximization (EM) algorithm with early stopping (convergence threshold: change in dormant proportion < 1×10^-6 or maximum 50 iterations) to estimate the dormant and expanding components.

As an alternative to GMM, we also classified tumors as dormant or proliferative using empirically derived cutoff values. Dormancy cutoffs were determined by calculating the kernel density estimate of log2-transformed cell numbers, identifying genes with bimodal size distributions and calculating the valley point between the first two peaks in their density distributions. The overall dormancy cutoff was computed by calculating the median of all valley points across bimodal genes, and tumors were classified as dormant if their cell number was less than or equal to the cutoff and expanding otherwise. The dormancy percentage statistic was calculated as the percentage of dormant tumors:

We then calculated a relative dormancy percentage by normalizing this number to t**h**e corresponding dormancy percentage calculated using sg*Inert* tumors:

Furthermore, the relative statistics were converted to log2 fold-change for downstream analy**sis** and visualization.

### Calculation of confidence intervals and P-values for tumor growth and number metrics

Confidence intervals and *P*-values were calculated using bootstrap resampling of tumors **to** estimate the sampling distribution of metastatic seeding, metastatic burden, and percentile sizes.. 10,0**0**0 bootstrap samples were drawn for all reported P-values. 95% confidence intervals were calculated usi**n**g the 2.5^th^ and 97.5^th^ percentile of the bootstrapped statistics. Because we calculate metrics of tumor grow**th** that are normalized to the same metrics in sgInert tumors, under the null model where genotype does n**ot** affect tumor growth, the test statistic is equal to 1. Two-sided p-values were thus calculated as followed:

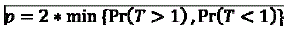

Where T is the test statistic and Pr(T>1) and Pr(T<1) were calculated empirically as the proportion **of** bootstrapped statistics that were more extreme than the baseline of 1. To account for multiple hypothe**si**s testing, p-values were FDR-adjusted using the Benjamini-Hochberg procedure as implemented in t**he** Python package stats models.

### Xenium data generation and analysis

Formalin-fixed paraffin-embedded (FFPE) blocks from metastases bearing C57BL/6 mouse liv**er** and lung were sectioned at 5 µm and mounted onto Xenium slides following a modified version of t**he** FFPE Tissue Preparation Guide (10x Genomics, CG000578, Rev D). Slides were deparaffinized wi**th** sequential xylene and ethanol washes, followed by decrosslinking using the FFPE Tissue Enhancer and diluted Perm Enzyme B (10x Genomics, CG000580, Rev E). Xenium Prime pre-designed and add-on custom priming oligos were hybridized to target RNAs in situ. After RNase treatment to release the RNA strand, a polishing step was performed, followed by overnight probe hybridization utilizing Xenium Prime 5K Mouse Pan Tissue & Pathways Panel (10x Genomics, 1000725) of probes for 5001 genes and an additional custom panel of probes for 100 genes (design ID: Q6VTXC). Probes were subsequently ligated and underwent rolling circle amplification, generating multiple copies of the gene-specific barcode for each RNA target. Cell segmentation reagents were applied during a staining workflow to label nuclei, membranes, and cytoplasmic regions for automated morphology-based segmentation. The background was quenched using an auto-fluorescence mixture, and nuclei were counterstained with DAPI to aid in sample tracking and cell boundary segmentation (10x Genomics, CG000760, Rev C).

Xenium slides were then loaded onto the 10X Xenium Analyzer instrument (10x Genomics, 1000481) for imaging and initial data processing following manufacturer’s instructions (10x Genomics, CG000584, Rev F). Fluorescently labeled oligos bind to the amplified DNA probes, and iterative rounds of probe hybridization, imaging, and stripping generated optical signatures for each barcode, which were decoded to gene identities. The H&E staining of the post-Xenium slides was performed after the run following the guidance provided (10X Genomics, CG000613, Rev B).

Following the optimized Xenium analysis pipeline^54^, we preprocessed and clustered Xenium data as follows: 1) After formatting each raw data as an Anndata structure, we then concatenated both groups into one Anndata with their group identifications. 2) Then, we filtered out the cells with a minimum of 40 transcripts and a minimum of 15 gene types. After that, we performed normalization first with CP10K and then with log1p. Then, we selected the top 3000 highly variable genes from a total of 5106 genes for each group. 3) We then used PCA to get the top 50 principal components, and performed neighbor analysis to find 30 closest neighbors in the feature matrix for each cell and construct a corresponding neighborhood graph. 4) Finally, we performed the Leiden clustering method to classify all cells into 15 clusters, and also performed UTAG and gene ranking for characterizing these clusters.

For Cell annotation, we annotated all cell clusters for their cell types based on the top 30 genes of each cluster and related marker genes of targeted cell types. The key cell types, like SCLC tumor cells and T cells, were further confirmed by their marker genes (e.g., ASCL1, CD3d/e/g).

For T cell analysis, we calculated the spatial neighbors of all tumor cells. The tumor neighbor was defined as the 20 µm distance to tumor cell centroids. All T cells within this distance were marked as proximal to tumors, and others were marked as distant from tumors. To analyze T cell subtypes, we further clustered all T cells into 4 groups: liver tissue-residue memory T cells (TRM), regulatory T cells (Treg), naïve T cells (naïve), and T cells without these specific subtype markers. The effector and exhausted states of TRMs were further calculated based on their marker genes.

### Statistical analysis for non-MOBA-seq experiments

Sample or experiment sizes were estimated based on similar experiments previously performed in our laboratory, as well as in the literature. Sample sizes are provided in the corresponding figure legends. All values are presented as mean ± SD, with individual data points shown in the figure when possible. Comparisons of parameters between two groups were made by two-tailed Student’s t-tests. The differences among several groups were evaluated by one-way ANOVA with Tukey-Kramer post hoc evaluation. p-values less than 0.05 and 0.01 were considered significant (*) or very significant (**), respectively.

### Statistics & Reproducibility

The statistical tests used for each analysis are described in detail in the sections above. All analyses of barcode sequencing data were performed in Python (3.10) and visualizations of data were performed in R (4.4.1). Sample sizes were determined based on our previous experience conducting similar experiments and, in the case of Moba-seq experiments, based on previously published power analyses^18^. For experiments using western blot as a readout, at least three independent experiments were repeated with similar results. Data collection and analysis were not performed blind to the conditions of the experiments. Analyses of barcode sequencing data used non-parametric statistics; therefore, no assumptions about the distribution of data were made. For all other analyses data distributions were assumed to be normal but this was not formally tested and individual data points are plotted to show distribution.

## DATA AVAILABILITY

The human cancer genomic data analyzed for the ICI treatment response of this manuscript were derived from the MSK-IMPACT project via cBioPortal (https://www.cbioportal.org/study/summary?id=msk_impact_2017). Processed data plotted in figures has been available in the Supplementary Information. All other data supporting the findings of this study are available from the corresponding author on reasonable request.

## CODE AVAILABILITY

The code used for data analysis in this study is available on GitHub (https://github.com/tanglab-2024)

