## Supplementary material for "Quantitative dissection of the metastatic cascade at single colony resolution": Figure S1-16

Roberts, Xu *et. al*

### Supplementary Figure 1

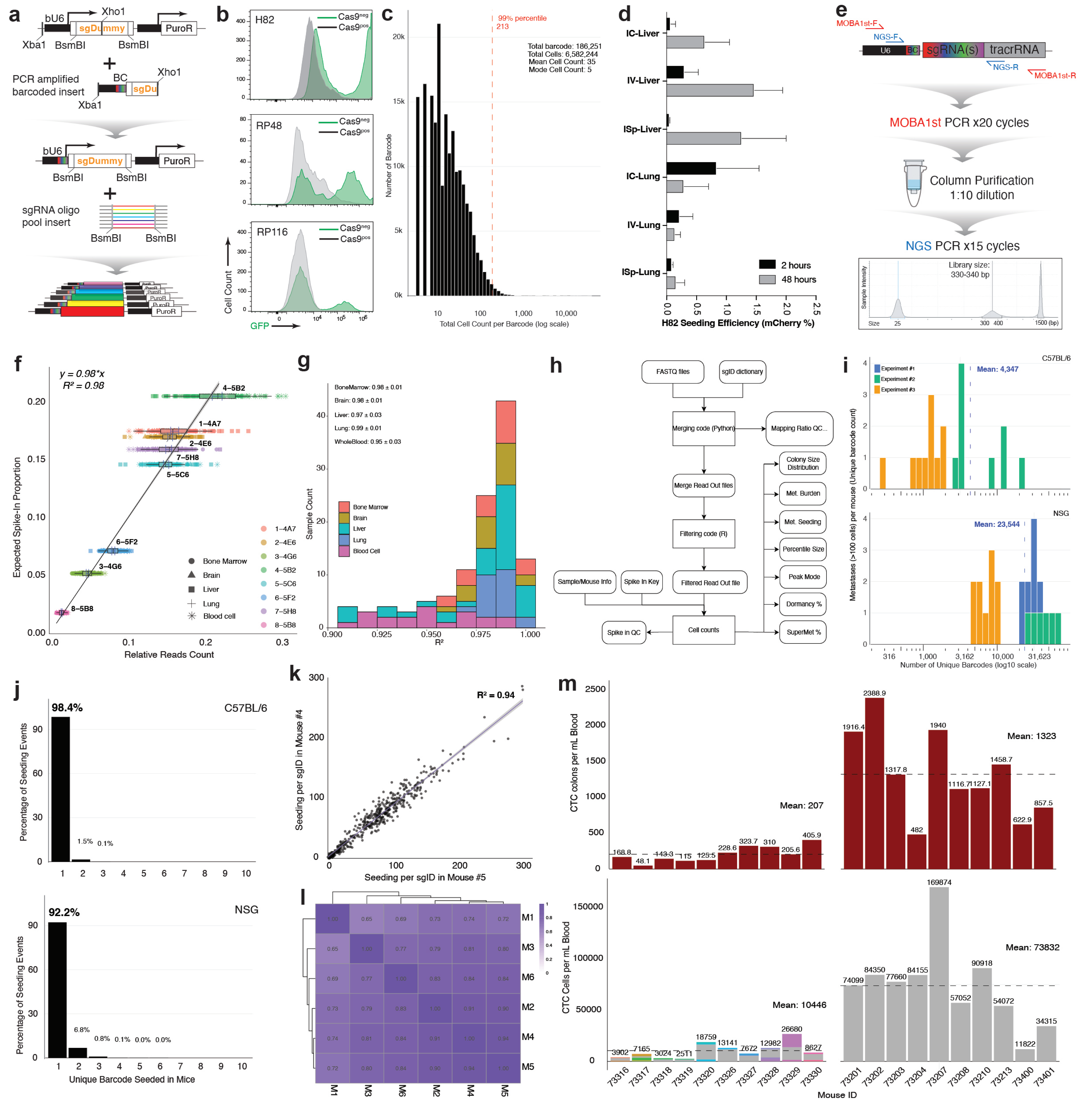

### Supplementary Figure 2

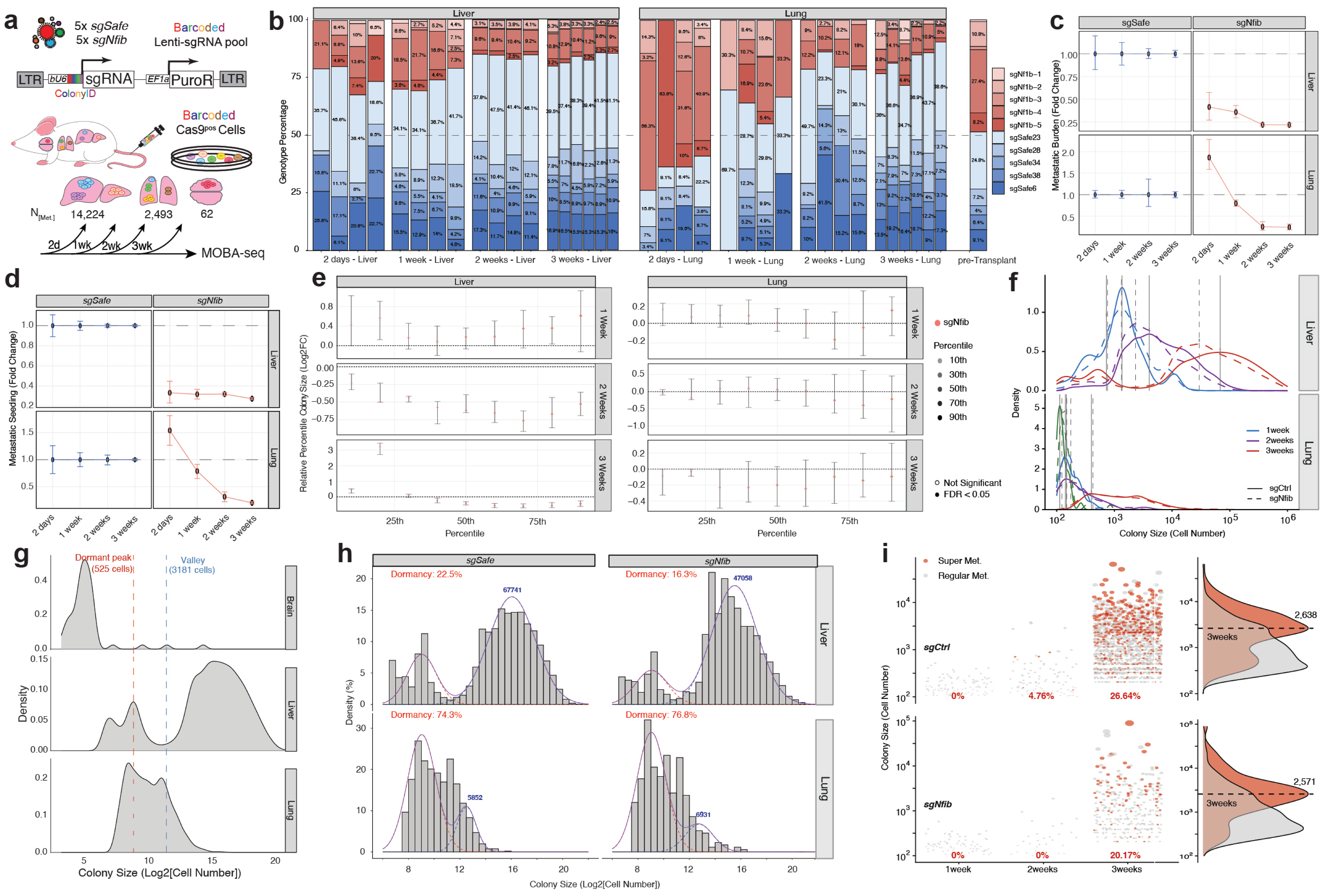

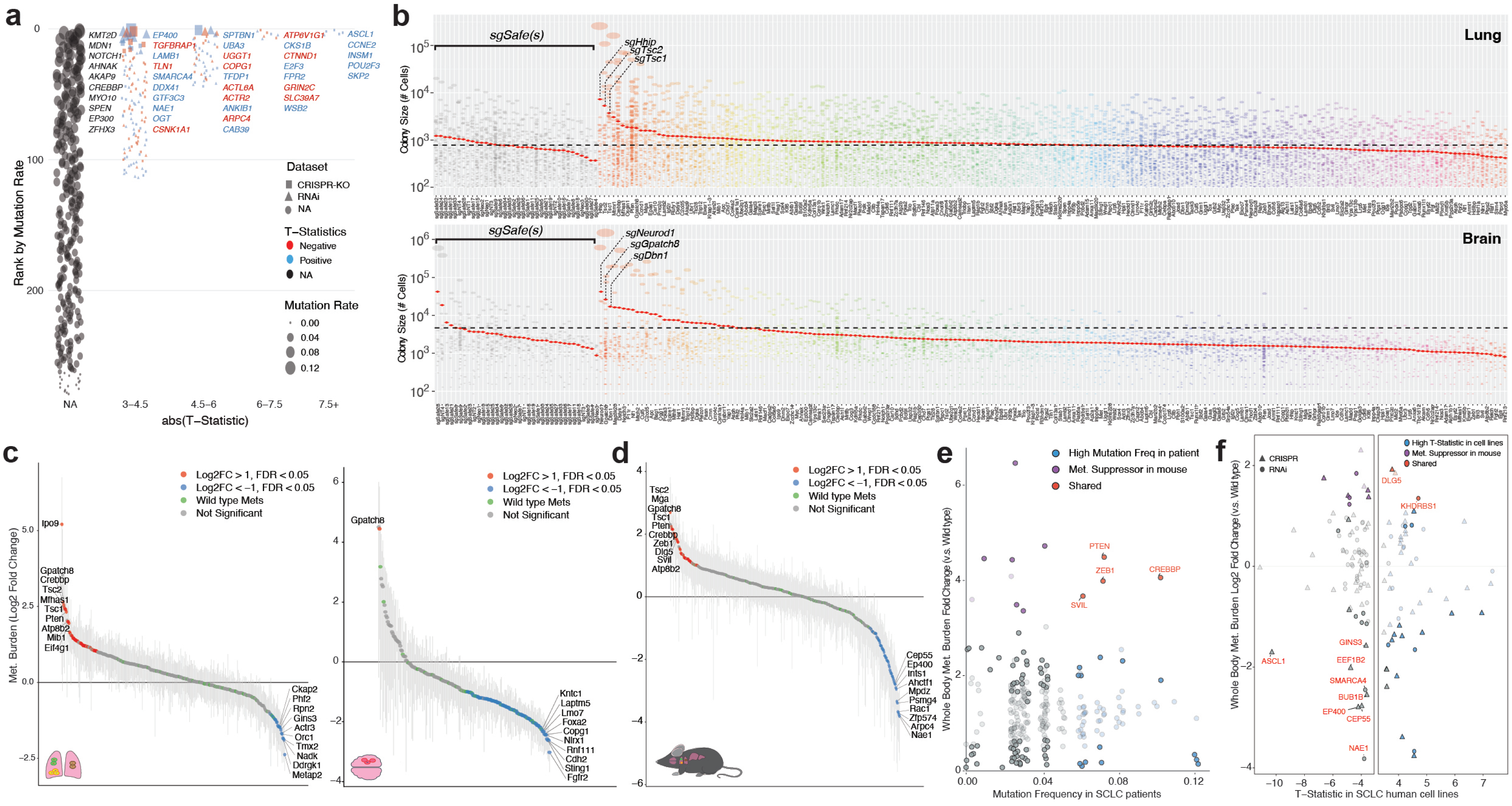
a
b
c
d
e
f
Rank by Mutation Rate
abs(T−Statistic)
NA
3–4.5
4.5–6
6–7.5
7.5+
KMT2D
MDN1
NOTCH1
AHNAK
AKAP9
CREBBP
MYO10
SPEN
EP300
ZFHX3
EP400
TGFBRAP1
LAMB1
TLN1
SMARCA4
DDX41
GTF3C3
NAE1
OGT
CSNK1A1
SPTBN1
UBA3
UGGT1
COPG1
TFDP1
ACTL8A
ACTR2
ANKIB1
ARPC4
CAB39
ATP6V1G1
CKS1B
CTNND1
E2F3
FPR2
GRIN2C
SLC39A7
WSB2
ASCL1
CCNE2
INSM1
POU2F3
SKP2
Dataset
CRISPR-KO
RNAi
NA
T−Statistics
Negative
Positive
NA
Mutation Rate
0.00
0.04
0.08
0.12
Lung
Brain
sgSafe(s)
sgHhip
sgTsc2
sgTsc1
sgNeurod1
sgGpatch8
sgDbn1
Colony Size (# Cells)
Met. Burden (Log2 Fold Change)
Log2FC > 1, FDR < 0.05
Log2FC < −1, FDR < 0.05
Wild type Mets
Not Significant
Ipo9
Gpatch8
Crebbp
Tsc2
Mfhas1
Tsc1
Pten
Atp8b2
Mib1
Eif4g1
Ckap2
Phf2
Rpn2
Gins3
Actr3
Orc1
Tmx2
Nadk
Ddrgk1
Metap2
Kntc1
Laptm5
Lmo7
Foxa2
Copg1
Nlrx1
Rnf111
Cdh2
Sting1
Fgfr2
Mga
Zeb1
Dlg5
Svil
Cep55
Ep400
Ints1
Ahctf1
Mpdz
Psmg4
Rac1
Zfp574
Arpc4
Nae1
High Mutation Freq in patient
Met. Suppressor in mouse
Shared
Whole Body Met. Burden Fold Change (v.s. Wild type)
Mutation Frequency in SCLC patients
PTEN
ZEB1
CREBBP
SVIL
CRISPR
RNAi
High T-Statistic in cell lines
Whole Body Met. Burden Log2 Fold Change (v.s. Wild type)
T−Statistic in SCLC human cell lines
DLG5
KHDRBS1
ASCL1
GINS3
EEF1B2
SMARCA4
BUB1B
EP400
CEP55
NAE1

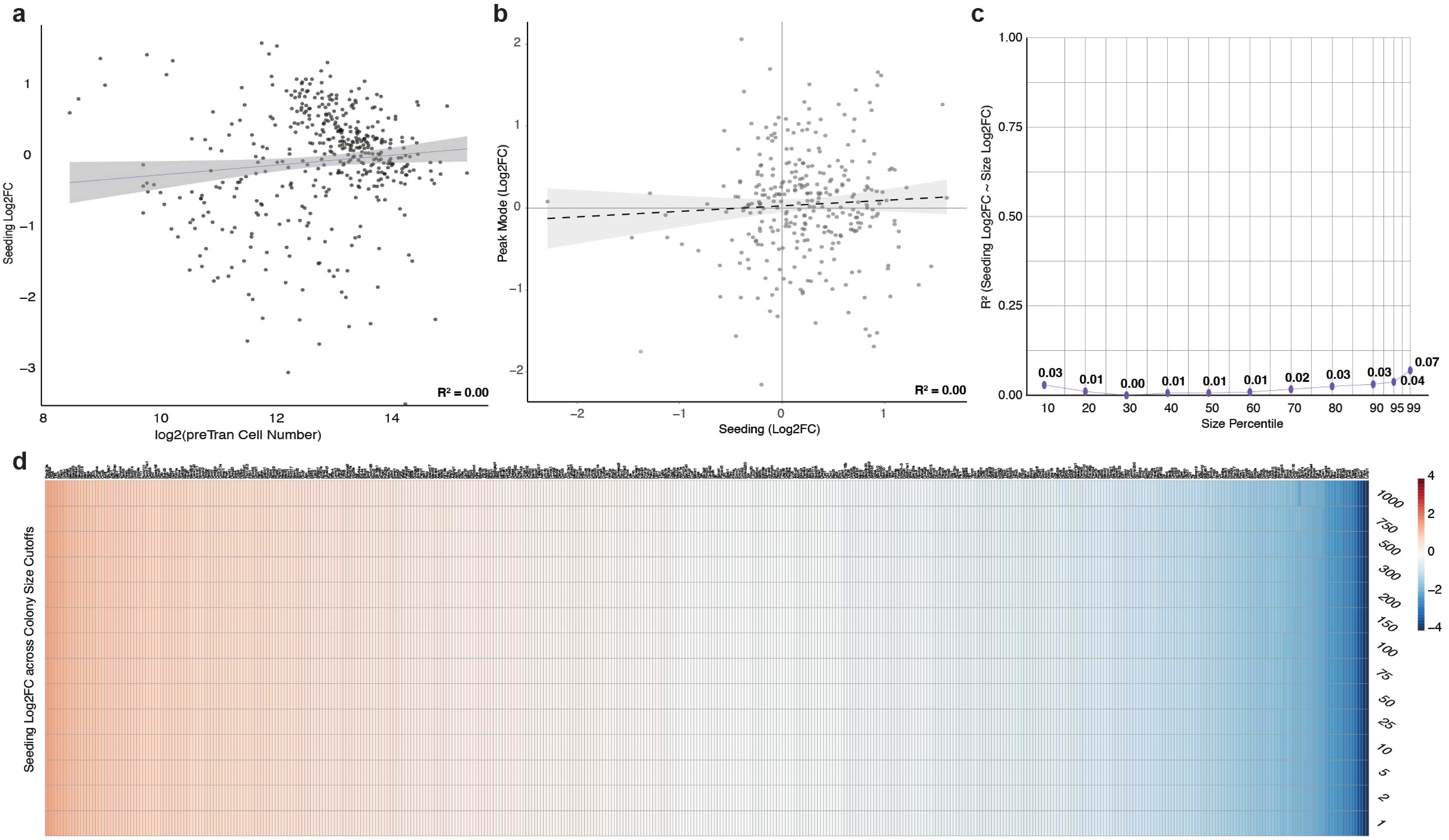
a
Seeding Log2FC
log2(preTran Cell Number)
$R^2 = 0.00$
b
Peak Mode (Log2FC)
Seeding (Log2FC)
$R^2 = 0.00$
c
$R^2$ (Seeding Log2FC ~ Size Log2FC)
Size Percentile
0.03
0.01
0.00
0.01
0.01
0.01
0.02
0.03
0.03
0.04
0.07
d
Seeding Log2FC across Colony Size Cutoffs
1000
750
500
300
200
150
100
75
50
25
10
5
2
1

Roberts, Xu *et. al*

### Supplementary Figure 5

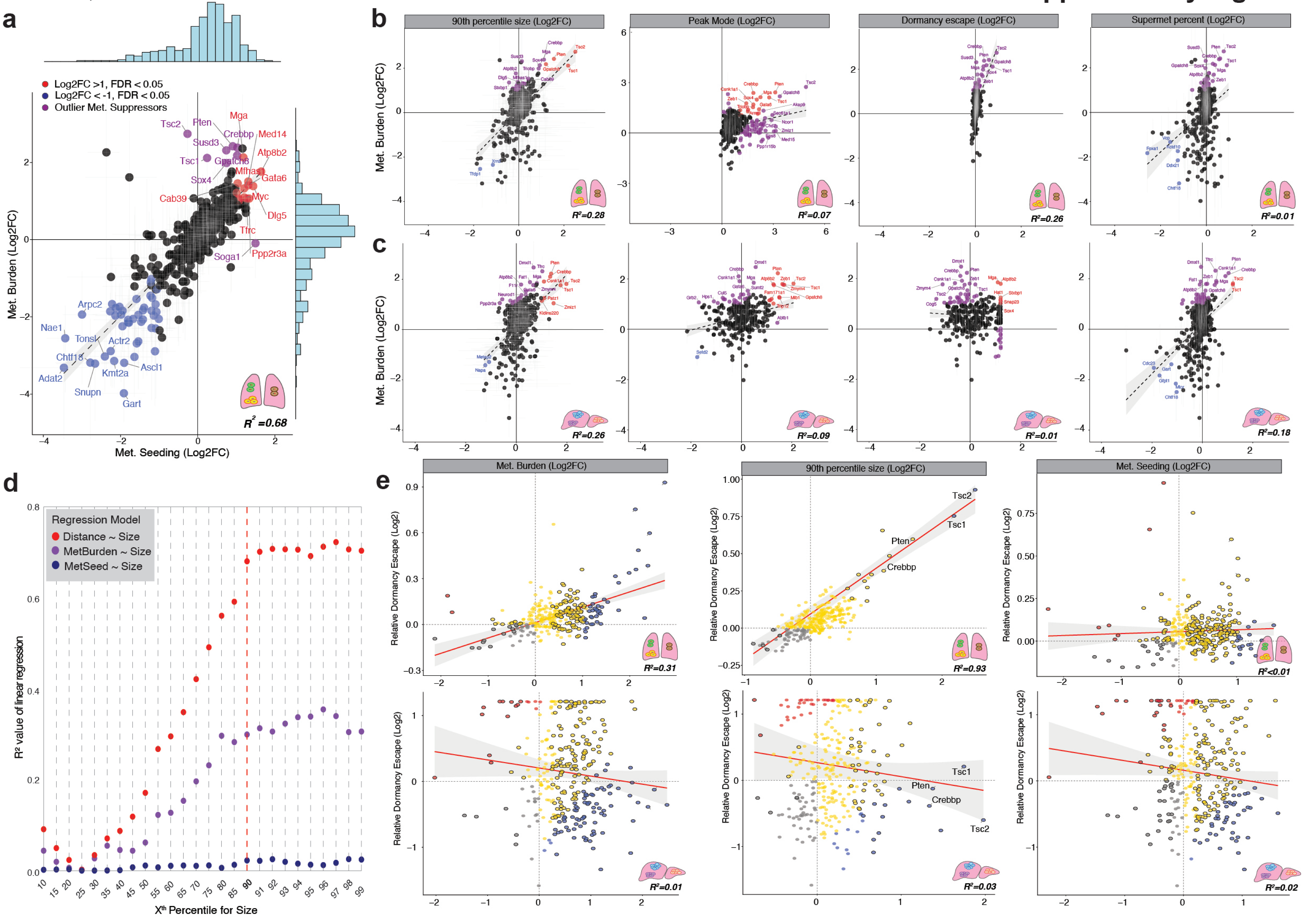

### Supplementary Figure 6

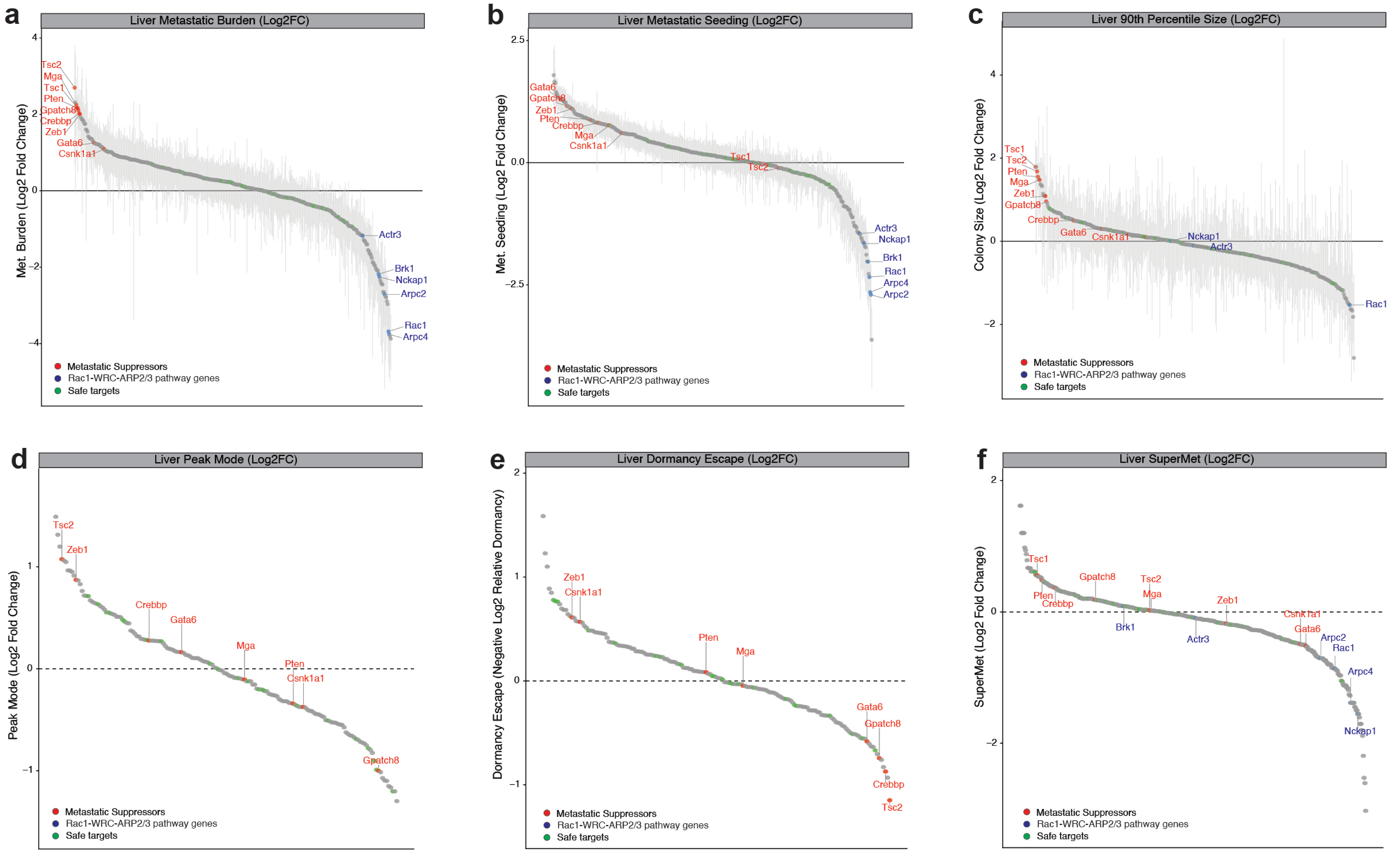

Roberts, Xu *et. al*

**Supplementary Figure 7**

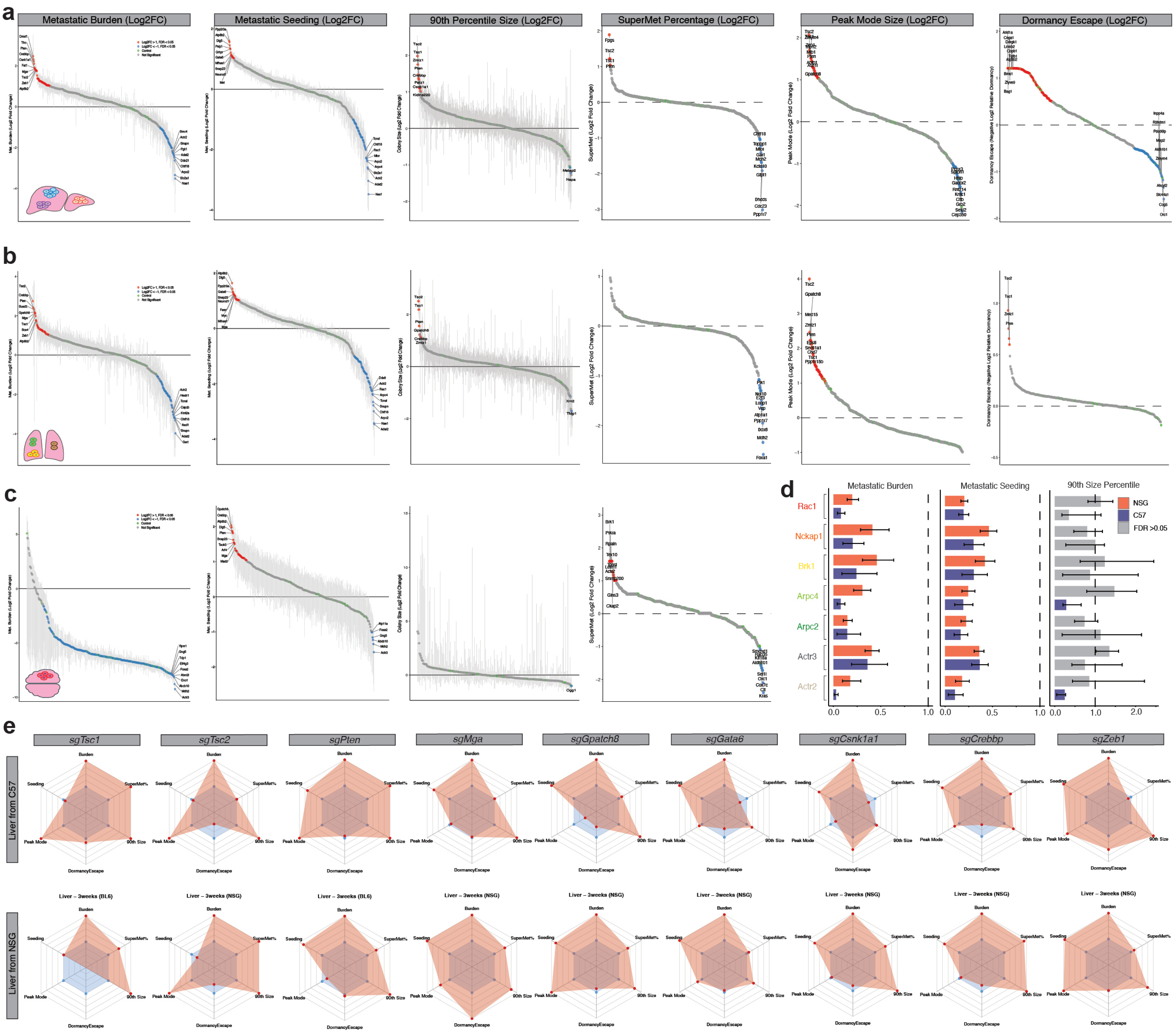

### Supplementary Figure 8

**a**

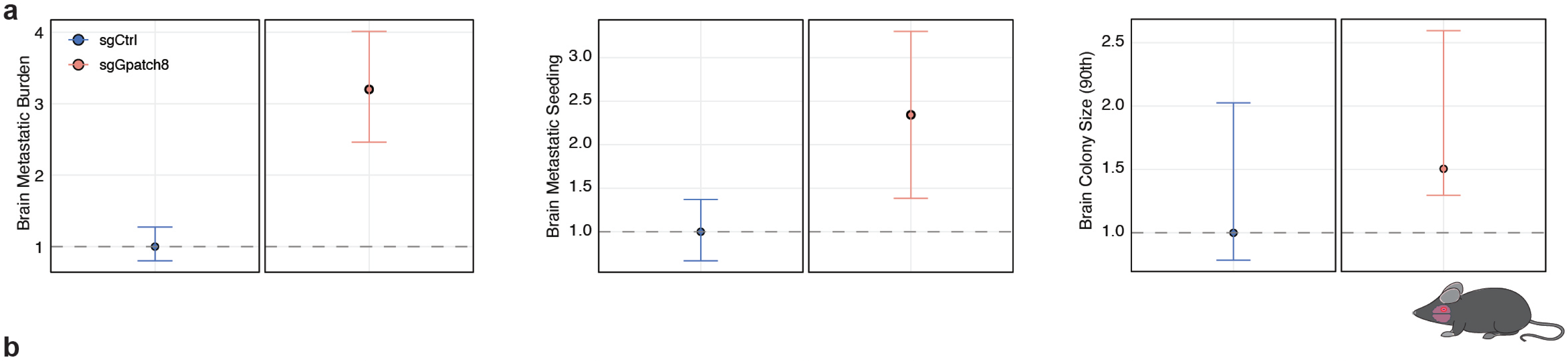


**b**

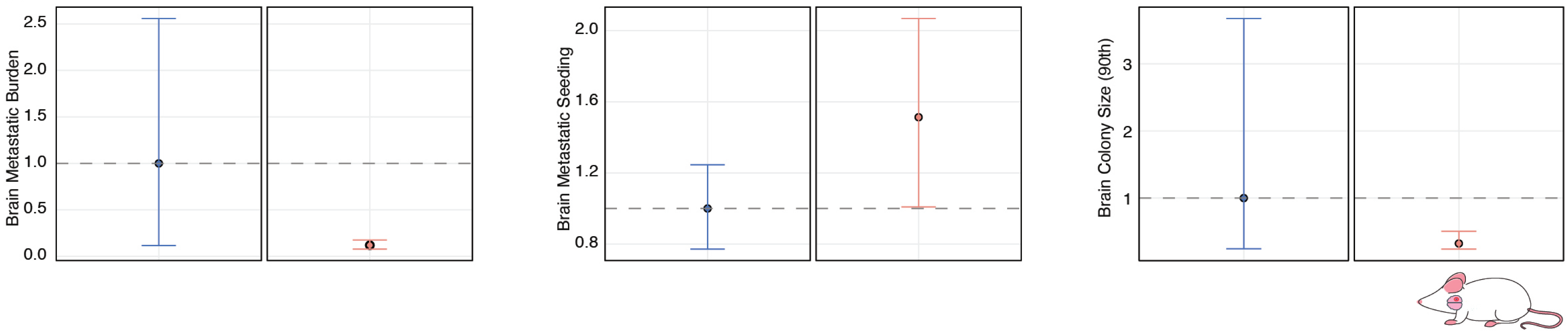

### Supplementary Figure 9

**a**

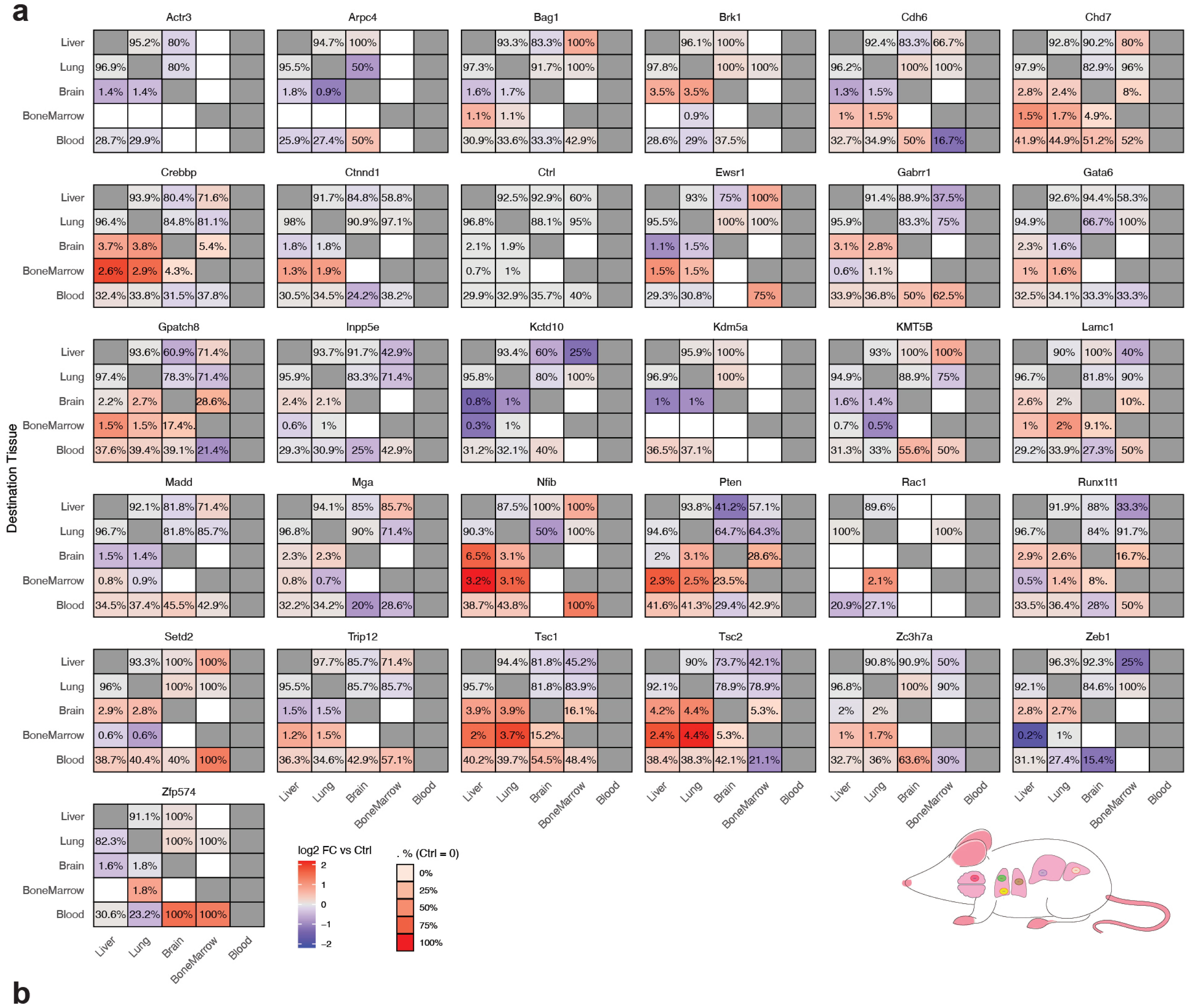


**b**

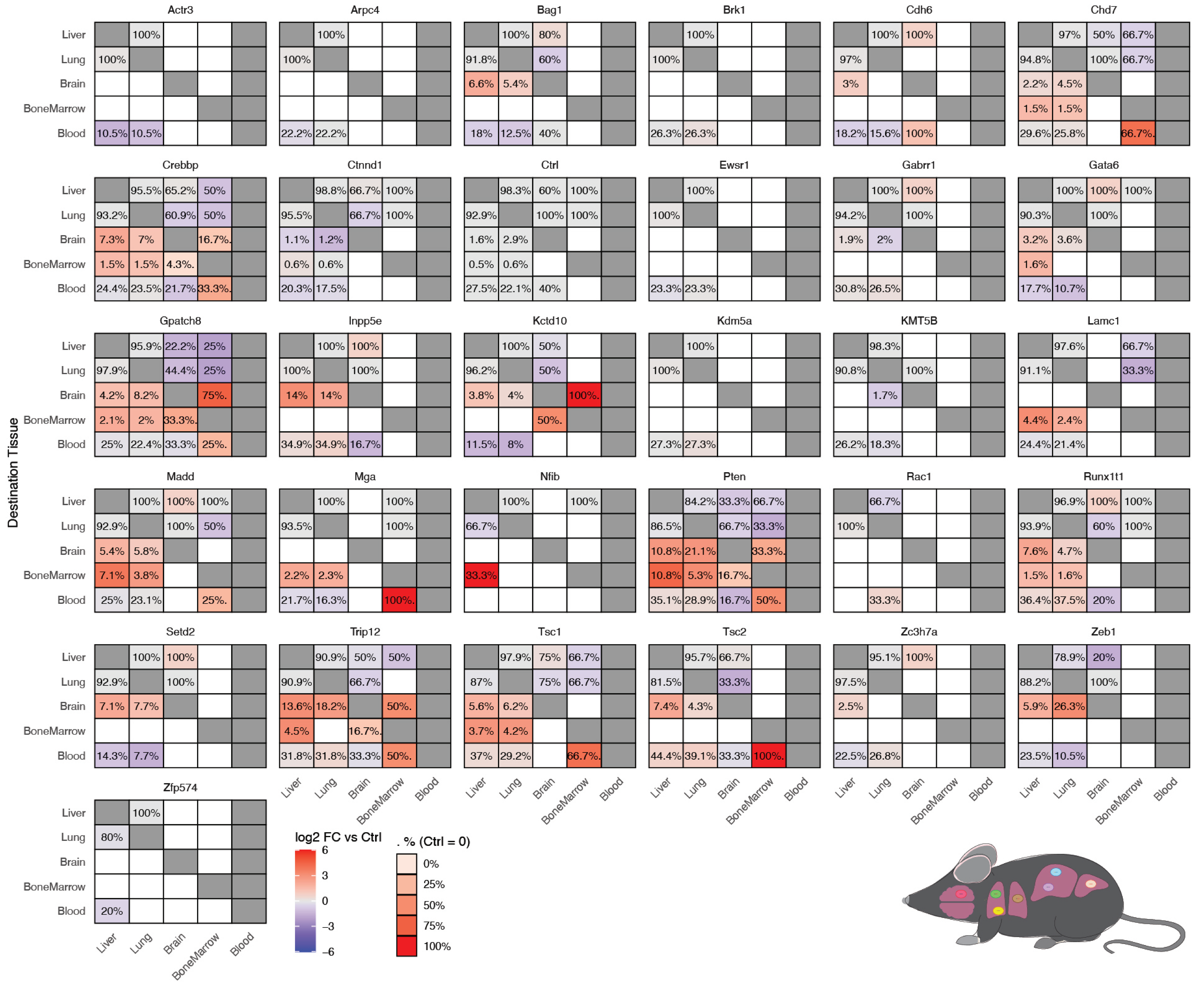

Roberts, Xu *et. al*

**Supplementary Figure 10**

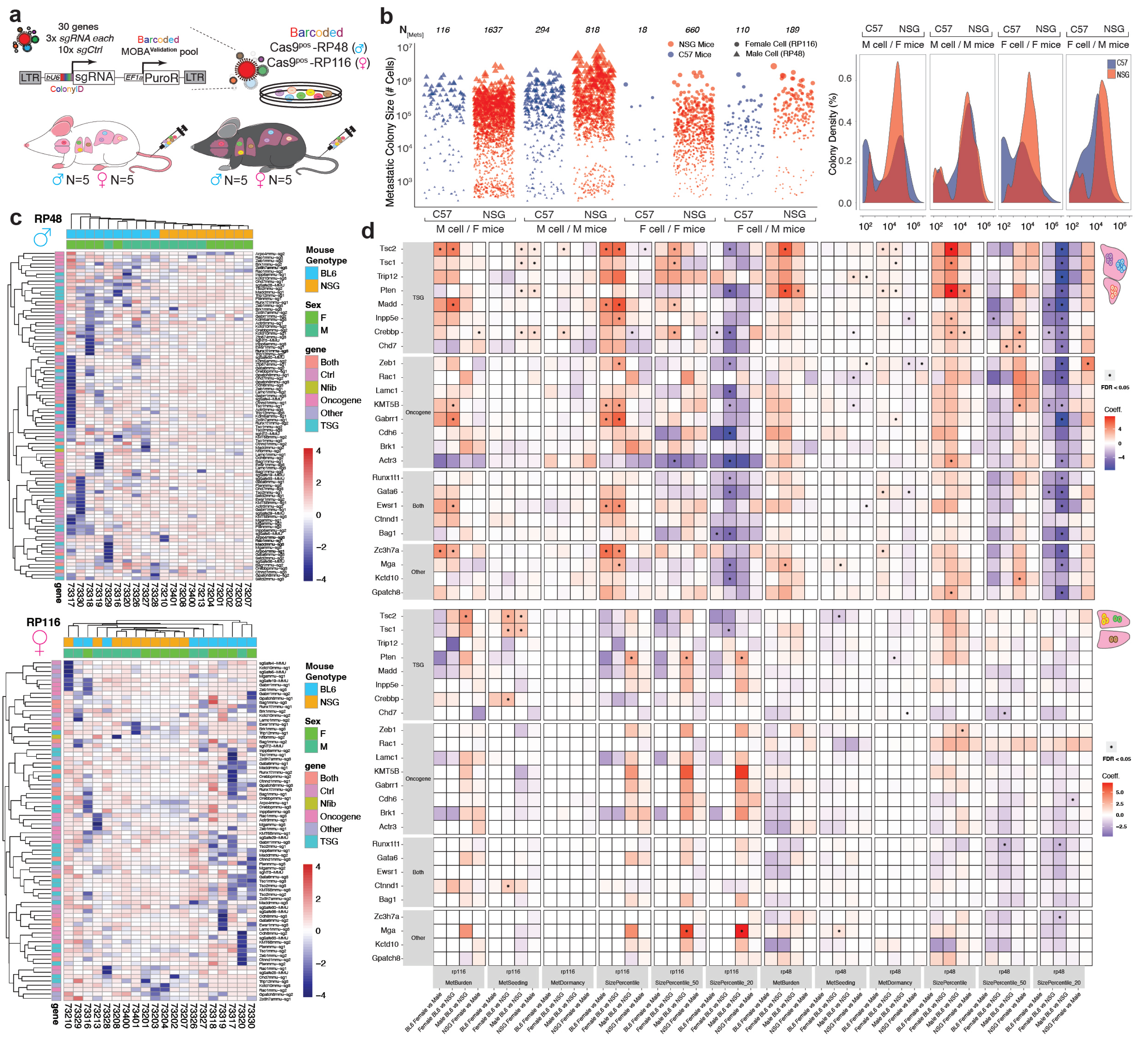

Roberts, Xu *et. al*

### Supplementary Figure 11

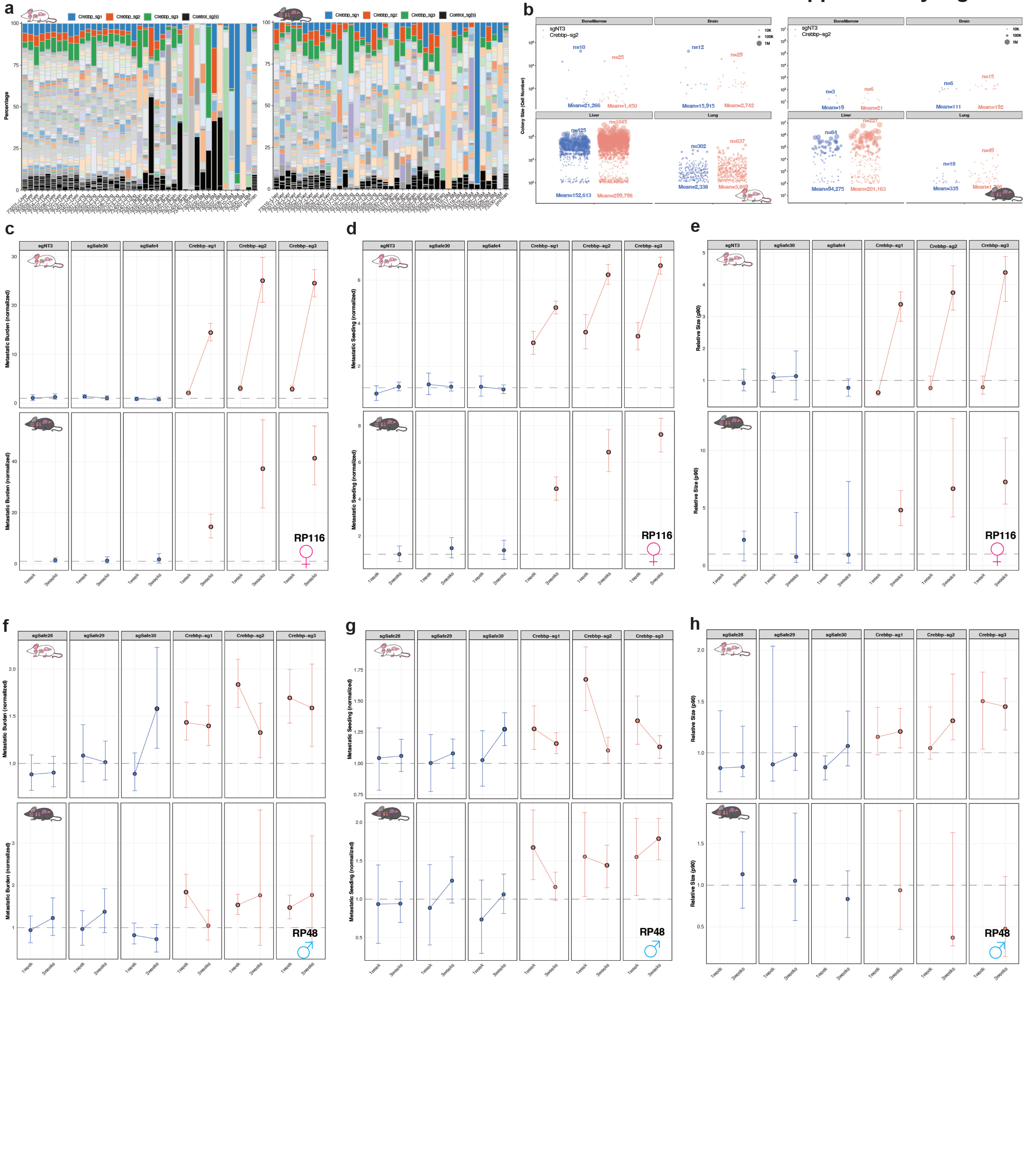

#### Supplementary Figure 12

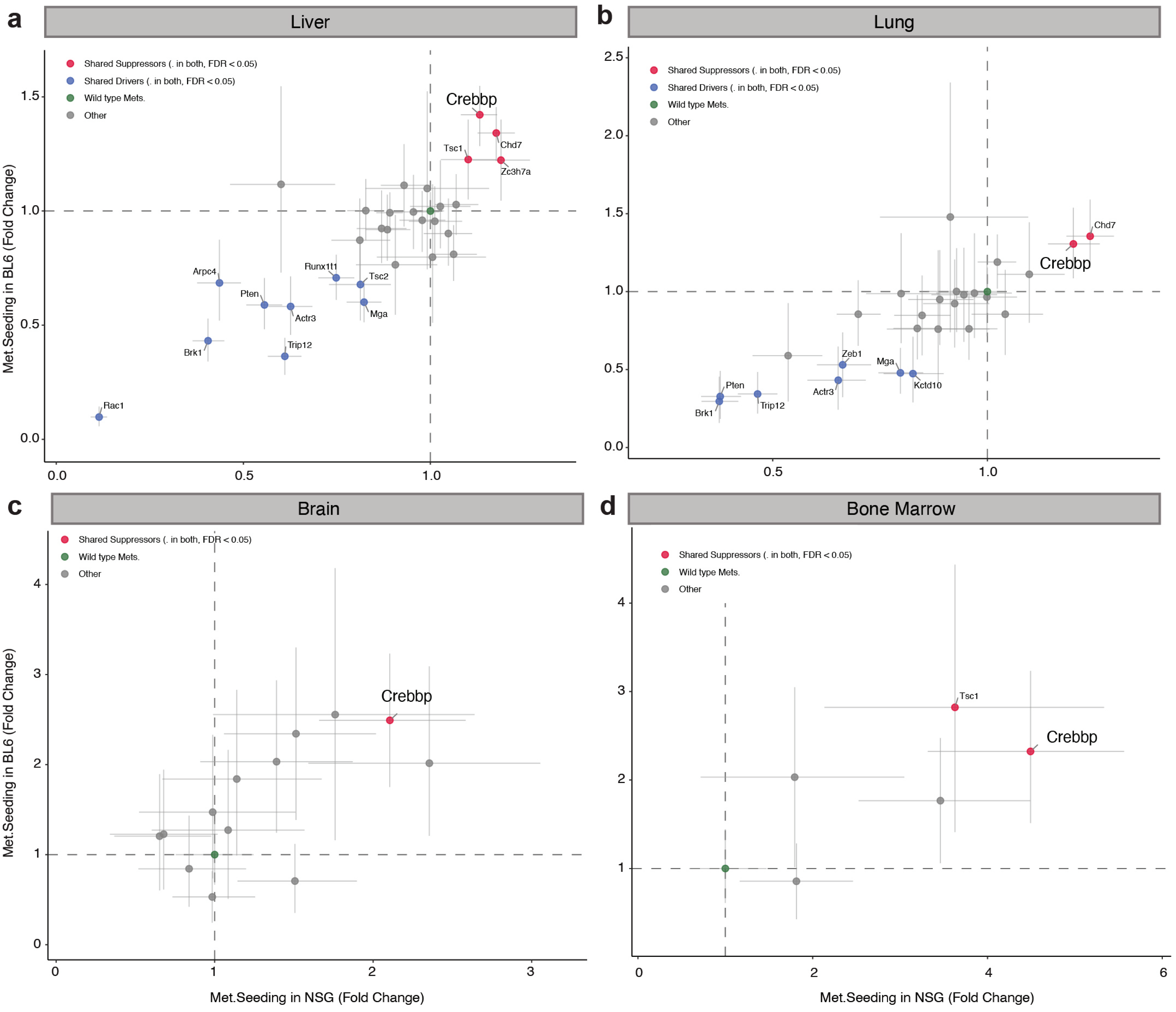

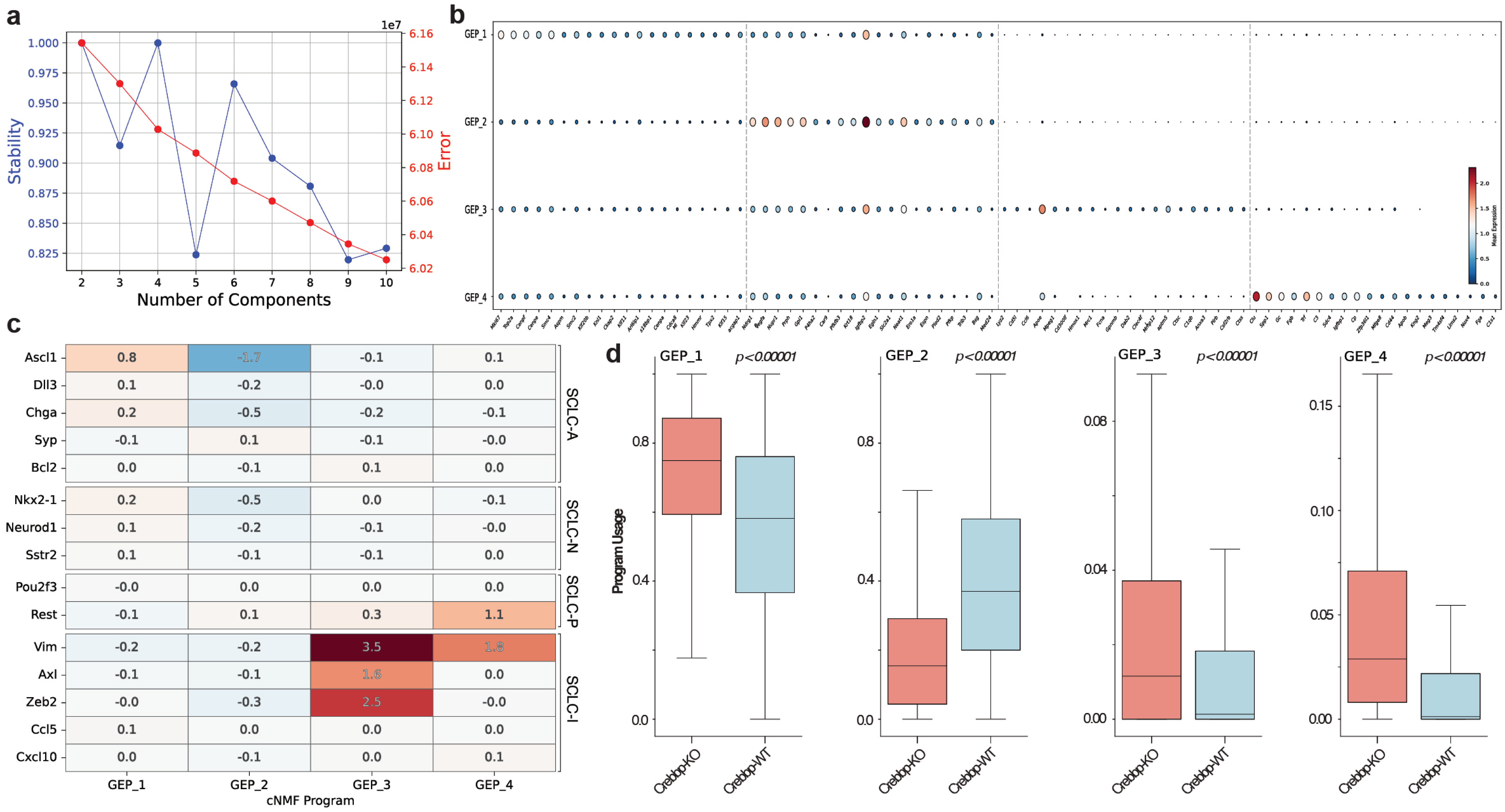
a
Stability
Error
1e7
Number of Components
b
GEP_1
GEP_2
GEP_3
GEP_4
Mean Expression
c
Ascl1
Dll3
Chga
Syp
Bcl2
Nkx2-1
Neurod1
Sstr2
Pou2f3
Rest
Vim
Axl
Zeb2
Ccl5
Cxcl10
SCLC-A
SCLC-N
SCLC-P
SCLC-I
GEP_1
GEP_2
GEP_3
GEP_4
cNMF Program
0.8 -1.7 -0.1 0.1
0.1 -0.2 -0.0 0.0
0.2 -0.5 -0.2 -0.1
-0.1 0.1 -0.1 -0.0
0.0 -0.1 0.1 0.0
0.2 -0.5 0.0 -0.1
0.1 -0.2 -0.1 -0.0
0.1 -0.1 -0.1 0.0
-0.0 0.0 0.0 0.0
-0.1 0.1 0.3 1.1
-0.2 -0.2 3.5 1.8
-0.1 -0.1 1.6 0.0
-0.0 -0.3 2.5 -0.0
0.1 0.0 0.0 0.0
0.0 -0.1 0.0 0.1
d
Program Usage
GEP_1 p<0.00001
GEP_2 p<0.00001
GEP_3 p<0.00001
GEP_4 p<0.00001
Crebbp-KO
Crebbp-WT

Roberts, Xu *et. al*

#### Supplementary Figure 14

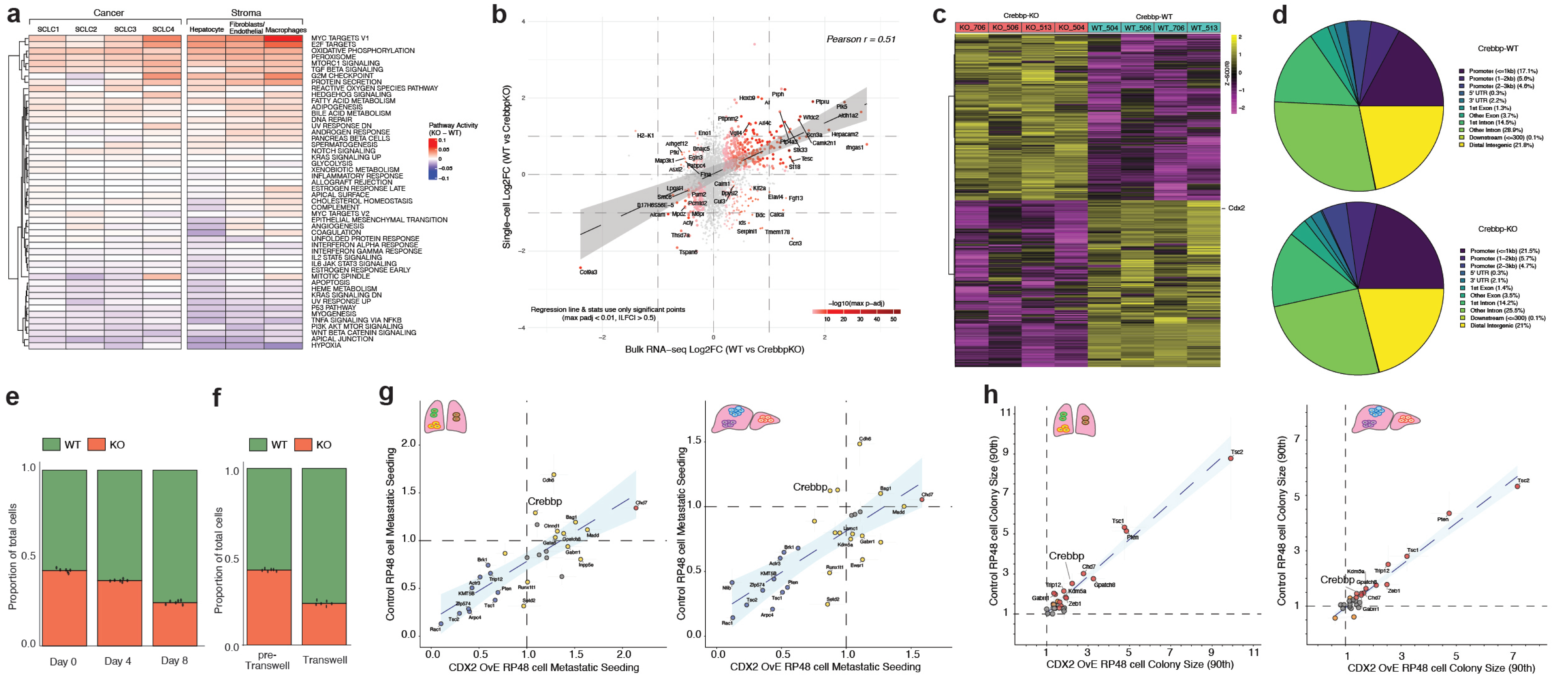

### Supplementary Figure 15

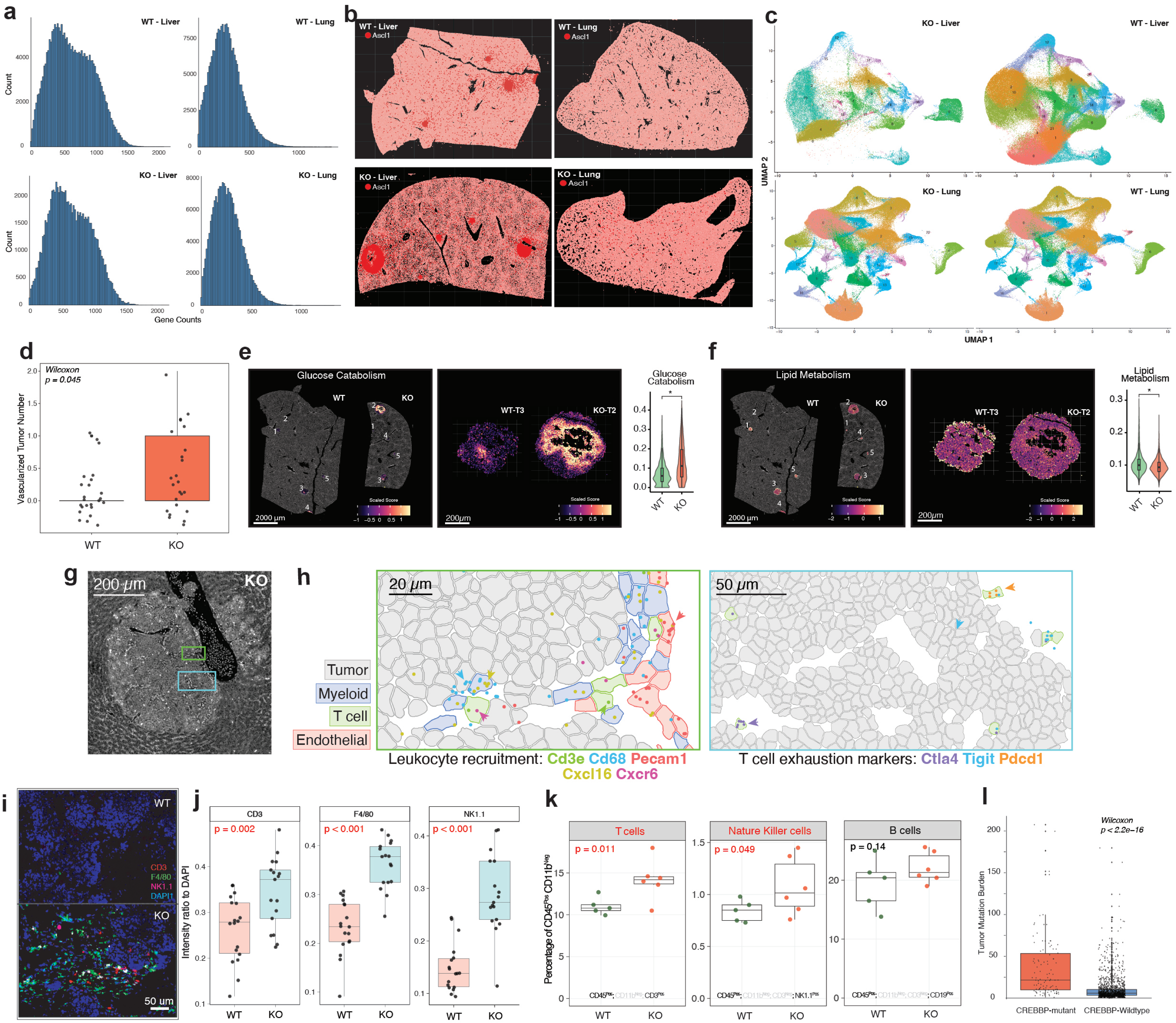

### Supplementary Figure 16

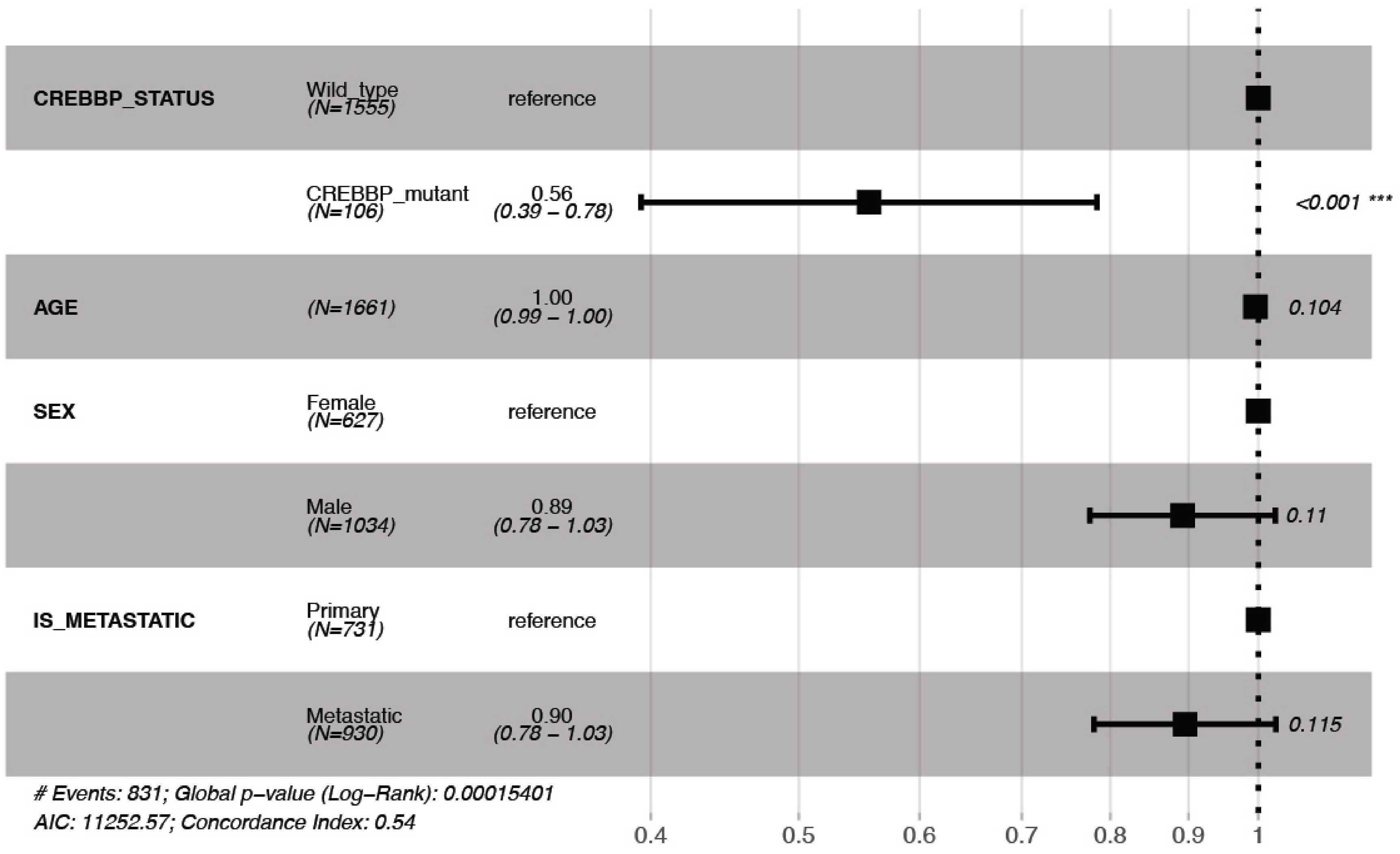
